# Improved bladder cancer antitumor efficacy with a recombinant BCG that releases a STING agonist

**DOI:** 10.1101/2023.12.15.571740

**Authors:** Peter K. Um, Monali Praharaj, Kara A. Lombardo, Takahiro Yoshida, Andres Matoso, Alex S. Baras, Liang Zhao, Geetha Srikrishna, Joy Huang, Pankaj Prasad, Max Kates, David McConkey, Drew M. Pardoll, William R. Bishai, Trinity J. Bivalacqua

**Affiliations:** Johns Hopkins University, School of Medicine, Department of Medicine, Center for Tuberculosis Research, Baltimore, USA; The Bloomberg-Kimmel Institute for Cancer Immunotherapy at Johns Hopkins, Baltimore, USA; Johns Hopkins University, School of Medicine, Department of Urology, Baltimore, USA; Department of Urology, Hyogo Prefectural Nishinomiya Hospital, Japan, 6620918; Department of Pathology, The Johns Hopkins University, Baltimore, USA; School of Medicine, Department of Surgery, University of Pennsylvania, Philadelphia, USA

## Abstract

Despite the introduction of several new agents for the treatment of bladder cancer (BC), intravesical BCG remains a first line agent for the management of non-muscle invasive bladder cancer. In this study we evaluated the antitumor efficacy in animal models of BC of a recombinant BCG known as BCG-*disA*-OE that releases the small molecule STING agonist c-di-AMP. We found that compared to wild-type BCG (BCG-WT), in both the orthotopic, carcinogen-induced rat MNU model and the heterotopic syngeneic mouse MB-49 model BCG-*disA*-OE afforded improved antitumor efficacy. A mouse safety evaluation further revealed that BCG-*disA*-OE proliferated to lesser degree than BCG-WT in BALB/c mice and displayed reduced lethality in SCID mice. To probe the mechanisms that may underlie these effects, we found that BCG-*disA*-OE was more potent than BCG-WT in eliciting IFN-β release by exposed macrophages, in reprogramming myeloid cell subsets towards an M1-like proinflammatory phenotypes, inducing epigenetic activation marks in proinflammatory cytokine promoters, and in shifting monocyte metabolomic profiles towards glycolysis. Many of the parameters elevated in cells exposed to BCG-*disA*-OE are associated with BCG-mediated trained innate immunity suggesting that STING agonist overexpression may enhance trained immunity. These results indicate that modifying BCG to release high levels of proinflammatory PAMP molecules such as the STING agonist c-di-AMP can enhance antitumor efficacy in bladder cancer.

## INTRODUCTION

Bladder cancer is the sixth most common malignancy in the United States, and the incidence of new cases of non-muscle invasive bladder cancer (NMIBC)—the most common form of bladder cancer—is approximately 63,000 per year in the US.^1,2^ Intravesical Bacillus Calmette Guerin (BCG) was introduced as an immunotherapy for NMIBC in the 1970s, and remains a first-line therapy despite the fact that BCG shortages have limited the supply in the US and other countries since 2019.^3,4^ Despite its first-line status for most forms of NMIBC, 20-40% of patients will relapse or fail to respond to BCG immunotherapy, and these patients are faced with limited therapeutic options such as cytotoxic chemotherapy or cystectomy.^5,6^ Thus, there is an unmet need for alternatives to standard BCG that may offer higher success rates and also may be effective in BCG-refractory, relapsing, or intolerant forms of NMIBC.^2^

Among responders, intravesical BCG has been shown to elicit a potent Th1 cellular immune response that is characterized by elevated production levels of IL-2, IL-12, IFNγ, and TNFα.^7,8^ BCG is also known to be rapidly internalized by phagocytic cells where it enters a phagosomal vesicle and may persist for days to weeks. Indeed, BCG is often recovered in the urine for several days following intravesical administration, and there have been reports of prolonged BCG bacteriuria for months.^9,10^ The Th1 immune response observed following BCG is associated with elevated levels of neutrophils, CD8+ T cells, NK cells and macrophage entering the urothelium, and these cells undoubtedly contribute to the antitumor activity of BCG.^11^

The cGAS-STING-TBK1-IRF3 pathway forms a key innate immune signaling pathway that was originally characterized as a component of antiviral immunity, but more recently has been appreciated to play a role in antitumor immunity.^12^ Cyclic GMP-ATP synthase (cGAS) is activated by cytosolic DNA to release the endogenous STING agonist, cyclic-GAMP (cGAMP), and activation of the pathway leads to a potent pro-inflammatory interferon response.^13^ Bacteria including BCG and other mycobacteria make low levels of related cyclic dinucleotides such as cyclic-di-AMP (c-di-AMP) and cyclic di-GMP (c-di-GMP) which serve as second messengers for bacterial processes.^14,15^ These bacterial-derived cyclic dinucleotides are recognized as pathogen-associated molecular patterns (PAMPs) by the STING pathway and similarly elicit potent pro-inflammatory immune responses.^16^

Small molecule STING agonists have been shown to have potent antitumor efficacy typically following intratumoral injection, and several such agents have been tested in human clinical trials as anti-cancer agents. We hypothesized that a recombinant BCG strain engineered to release high levels of its endogenous STING agonist might offer more potent antitumor efficacy than wild-type BCG (BCG-WT). We previously reported the construction of a recombinant BCG, known as BCG-*disA*-OE, in which the endogenous BCG *disA* gene (that encodes a c-di-AMP-generating di-adenylate cyclase) is fused to a strong mycobacterial promoter.^17^ Compared to BCG-WT, BCG-*disA*-OE was found to be more potent in preventing tuberculosis disease progression in the guinea pig model. A previous report by us on BCG-*disA*-OE and bladder cancer contained data irregularities discovered post-publication leading us to retract that paper.^18^ In the present work we report new unpublished data as well as portions of the previous paper that are devoid of irregularities. Herein, using two well-established animal models of bladder cancer, we show that BCG-*disA*-OE provides improved antitumor efficacy compared to BCG-WT. We also characterize the safety of BCG-*disA*-OE as compared to BCG-WT, and we profile the comparative immune responses elicited by the two versions of BCG.

## RESULTS

### Macrophages that engulf BCG-*disA*-OE elicit a greater pro-inflammatory cytokine response when compared to BCG-WT

BCG-*disA*-OE is a genetically engineered BCG strain capable of intracellular delivery of a STING agonist, cyclic-di-AMP. This is achieved by fusion of an endogenous di-adenylate cyclase gene, *disA*, to a strong promoter, leading to a 300-fold overexpression of *disA* and a 15-fold increase in production of cyclic di-AMP (**Fig. 1**, **Fig. S1a**).^15^ This excess cyclic di-AMP production greatly enhances the STING pathway response via IRF3 induction (**Fig S1b**). To account for the fact that numerous BCG strains are used worldwide and variability in their clinical efficacies have been described, we generated two versions of BCG-*disA*-OE and their corresponding wild-type parental strains (BCG-WT): one using BCG-Tice and one using BCG-Pasteur. We did not detect major differences between the Tice and Pasteur versions.

**Figure 1.**
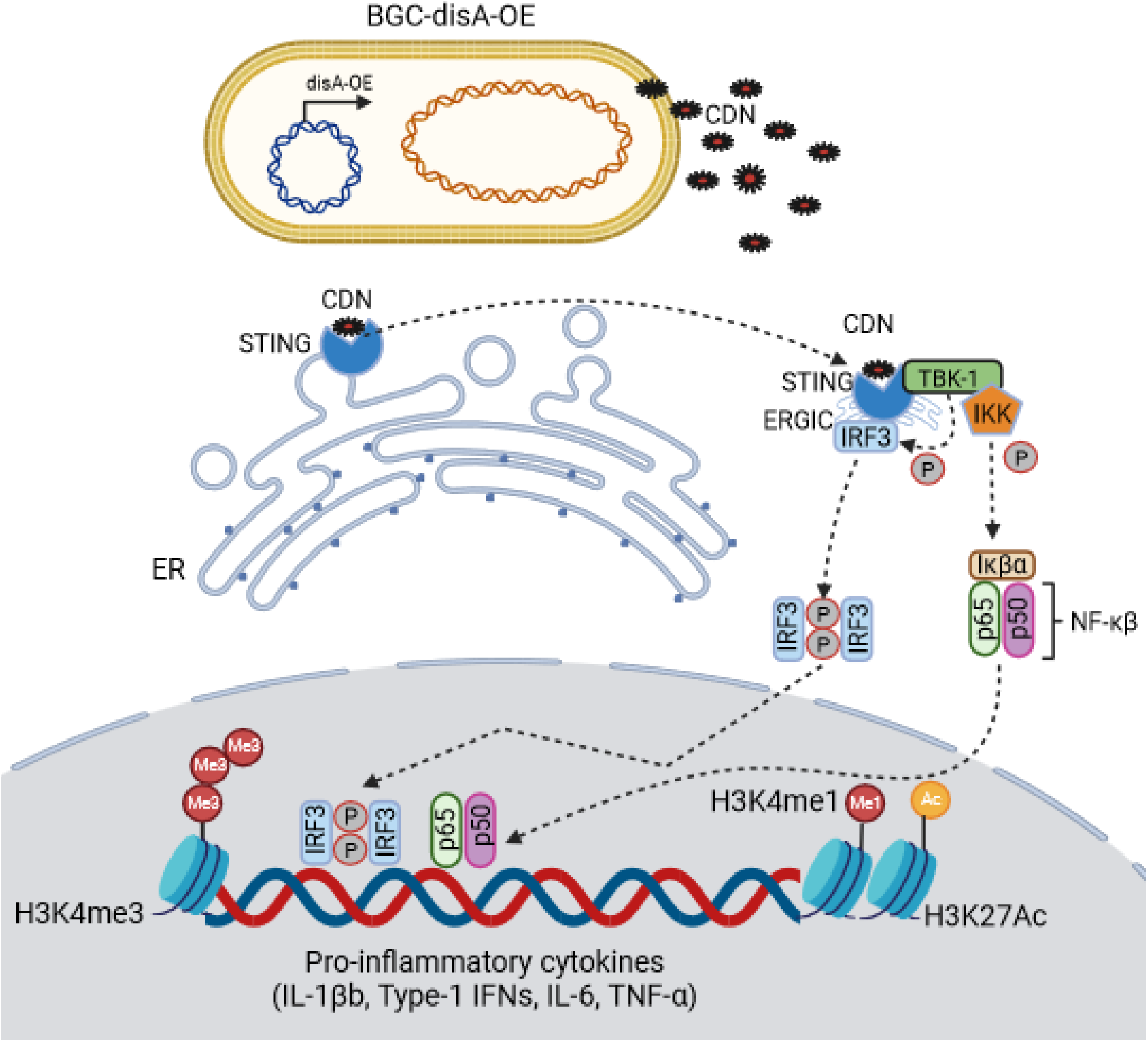
**A schematic diagram** of BCG-*disA*-OE’s intra-cellular delivery of cyclic di-nucleotides (CDNs) and subsequent binding to the STING homodimer; STING-CDN trafficking to the Golgi via ER-Golgi intermediate compartment (ERGIC) and then TBK-1 recruitment and subsequent phosphorylation of IRF3 and NFκβ for downstream transcriptional activation of pro-inflammatory cytokines.

### Antitumor efficacy of BCG-*disA*-OE compared to BCG-WT in the rat heterotopic, carcinogen (MNU)-induced bladder cancer model

Since the mid-1970s, BCG has served as a first-line immunotherapy for the treatment of NMIBC. Recent studies indicate that BCG exerts its antitumor effects via a trained immunity mechanism.^11^ We sought to determine if augmenting BCG with excess cyclic di-AMP release may improve bladder cancer outcomes relevant animal models. We tested BCG-*disA*-OE versus BCG-WT in the rat orthotopic, carcinogen-induced bladder cancer model in which intravesical therapies can be introduced into the bladder as they are in humans with NMIBC. The rat *N*-methyl-*N*-nitrosourea (MNU) model of bladder cancer (BC) is schematized in **Fig. 2a**.^19,20^ In this model urothelial dysplasia develops at 14 weeks after the final intravesical instillation of MNU; by week 24 rats display different forms of urothelial cancer severity including carcinoma-*in-situ* (CIS), papillary Ta (superficial), or higher-grade T1-T2 urothelial carcinoma with histopathologic and immunophenotypic features similar to those observed in human bladder cancer.^20–22^ Following carcinogen-mediated tumor induction with 4 weekly cycles of MNU (week 0, week 2, week 4, week 6), groups of rats were treated with 6 weekly doses of intravesical BCG-*disA*-OE, BCG-WT, or mock treatment from week 18-23 as is done for BCG induction therapy for humans with NMIBC. Upon necropsy at week 24 we divided the rat urinary bladders into portions for (i) RT-qPCR analysis, and (ii) histologic analysis including tumor staging by a blinded genitourinary pathologist. Transcriptional analysis of the whole excised bladders at week 23 showed that compared with BCG-WT, BCG-*disA*-OE elicited increased levels of mRNA for IFN-β, IFN-γ, IL-1β CXCL10, Mcp-1, MIP-1α, and Nos2 transcription. While mRNA levels of the immunosuppressive cytokines IL-10, TGF-β were not altered by both BCG strains; when BCG or BCG-*disA*-OE were compared to mock, we found BCG-*disA*-OE elicited a trend towards increased levels of mRNA for TNF-α (**Fig. 2b**). Correspondingly, we found a significant decrease in highest pathology grade (**Fig. 2c**), tumor involvement index (**Fig. 2d**) and highest tumor stage (**Fig. 2e**) in MNU rats treated with BCG-*disA*-OE over BCG-WT when compared to mock. By tumor involvement index, BCG-*disA*-OE was significantly superior to mock (p < 0.04), whereas BCG-WT showed non-significant trend towards improvement over mock. Importantly, the highest tumor stage observed in BCG-*disA*-OE-treated rats was Cis, whereas it was T1 in those receiving BCG-WT, and T2 in mock treated rats. 53.33% of BCG-*disA*-OE-treated rats were cancer free (p=0.0074), while only 31.25% of BCG-WT-treated rats were cancer-free (p=0.0598) compared to 0% of mock (**Fig. 2f**). Immunohistochemical (IHC) analyses revealed a significant reduction in Ki67 staining in BCG-*disA*-OE-treated MNU rat bladders when compared to mock (p < 0.001) and to a lesser yet significant extent in BCG-WT (p < 0.05) suggesting reduced tumor proliferation (**Fig. 2g**). CD68 IHC staining of rat bladders showed a slight trend towards higher levels of macrophage recruitment (**Fig. 2h**). We did not see a significant difference in IHC staining for pro-inflammatory CD86+ macrophages (M1-like) (**Fig. 2h**). However, we observed a significant reduction in IHC staining for CD206+ immunosuppressive (M2-like) macrophages that are associated with tumor promotion in the BCG-*disA*-OE-treated rats compared with mock controls (**Fig. 2h**). These observations indicate that the enhanced induction of type I IFN and other proinflammatory signatures in bladders of tumor-bearing rats treated with BCG-*disA*-OE correlated with the enhanced antitumor activity of the recombinant BCG strain.

**Figure 2.**
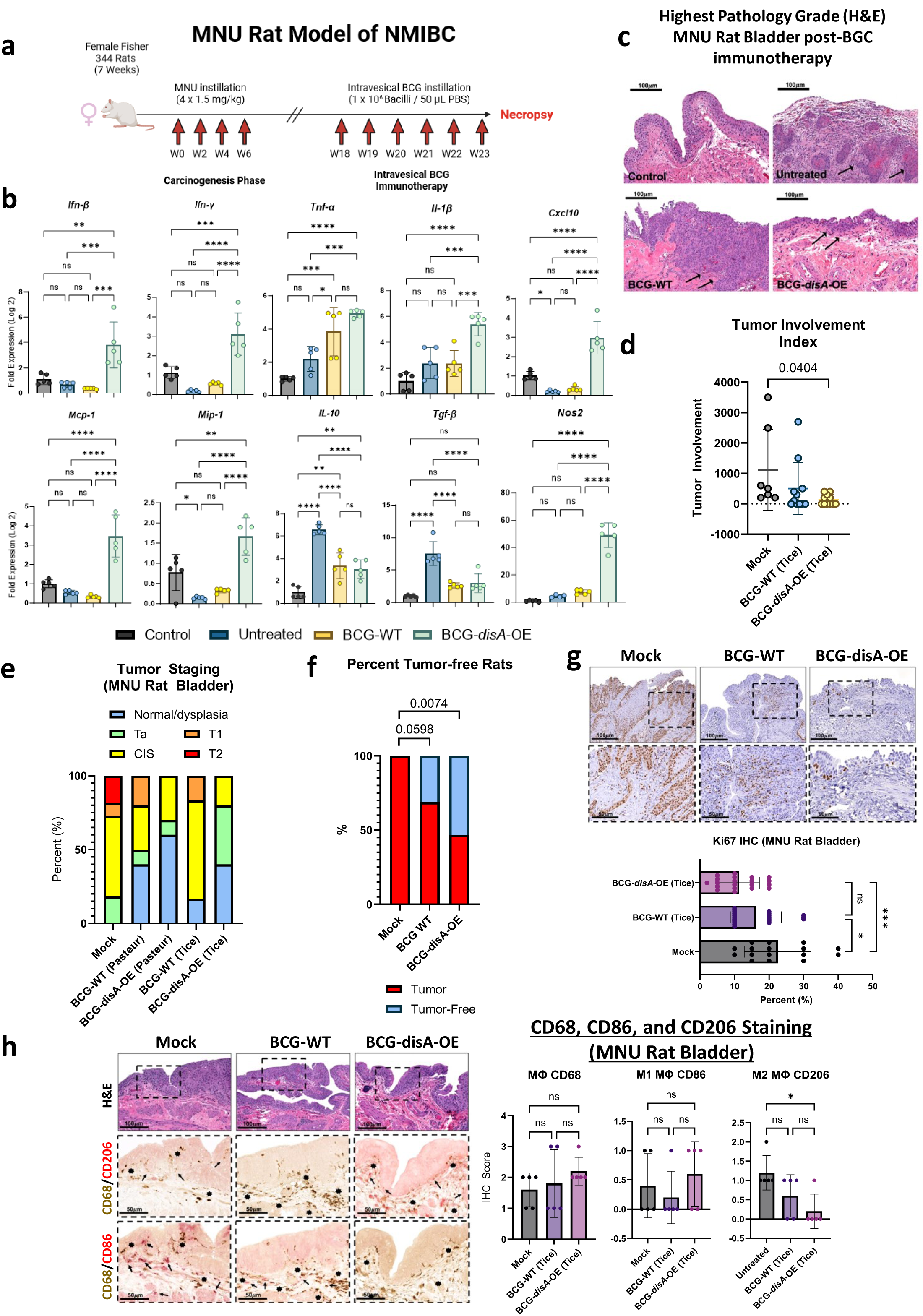
BCG-*disA*-OE elicits improved antitumor efficacy over BCG-WT in an orthotopic carcinogen-induced MNU rat model of urothelial cancer. **a.** Schematic diagram of the MNU rat model of NMIBC. **b.** mRNA levels for proinflammatory cytokines (IFN-β, IFN-γ, TNF-α, IL-1β), regulatory chemokines (CXCL10, Mcp-1, MIP-1α), immunosuppressive M2-like macrophage cytokines (IL-10, TGF-β), and the M1-like tumoricidal effector (Nos2) in whole bladders at necropsy (wk 23) measured by RT-qPCR relative to GAPDH (n= 5 animals / group). **c.** Representative H & E staining showing highest pathology grade for each group (control, untreated MNU bladder). **d.** Tumor involvement values at necropsy **e.** Tumor stage at necropsy. **f.** Percent of rats which were cancer-free at necropsy; BCG-WT (Pasteur and Tice), and BCG-*disA*-OE (Pasteur and Tice). g. Representative immunohistochemistry and bar graph of rat bladder tissue at necropsy stained for Ki67. **h.** Representative immunohistochemical co-staining and graph for CD68 (brown), CD86 (M1-like macrophages; red) and CD206 (M2-like macrophages; red) in rat bladder tissues at necropsy. Tumor staging and involvement index was performed by a pathologist trained urothelial cancers who was blinded to sample identities. The MNU model was conducted twice with BCG strains from the Tice background and the Pasteur background. Data shown represent pooled results from the two studies (n = 11-16 animals per group). Data are represented as mean values ± S.D. Statistical analyses were done using one-way ANOVA with Tukey’s test for multiple comparisons in panels **b, g,** & **h**; one-way ANOVA with Dunnett’s Test for multiple comparisons in panel **d**; two-sided Fisher’s Exact test in panel **f** (* p < 0.05, ** p < 0.01, *** p < 0.001, ****p < 0.0001).

### Antitumor efficacy of BCG-*disA*-OE compared to BCG-WT in the mouse heterotopic, syngeneic MB49 bladder cancer model

We also tested the functional efficacy of BCG-*disA*-OE in a murine heterotopic, syngeneic cancer model using MB49 urothelial cancer cells. Following flank engraftment with MB49 tumor cells, mice received four intra-tumoral treatments over 12 days as shown in **Fig. 3a**. In this model BCG-*disA*-OE also showed significantly more antitumor efficacy than BCG-WT as measured by tumor volumes after intra-tumoral injection of BCG-*disA*-OE when compared with BCG-WT (**Fig. 3b** and **Fig. S2a**). Histopathology demonstrated extensive necrosis and congestion in MB49 tumors treated with BCG-*disA*-OE when compared to BCG-WT and untreated (**Fig. S2b**).

**Figure 3.**
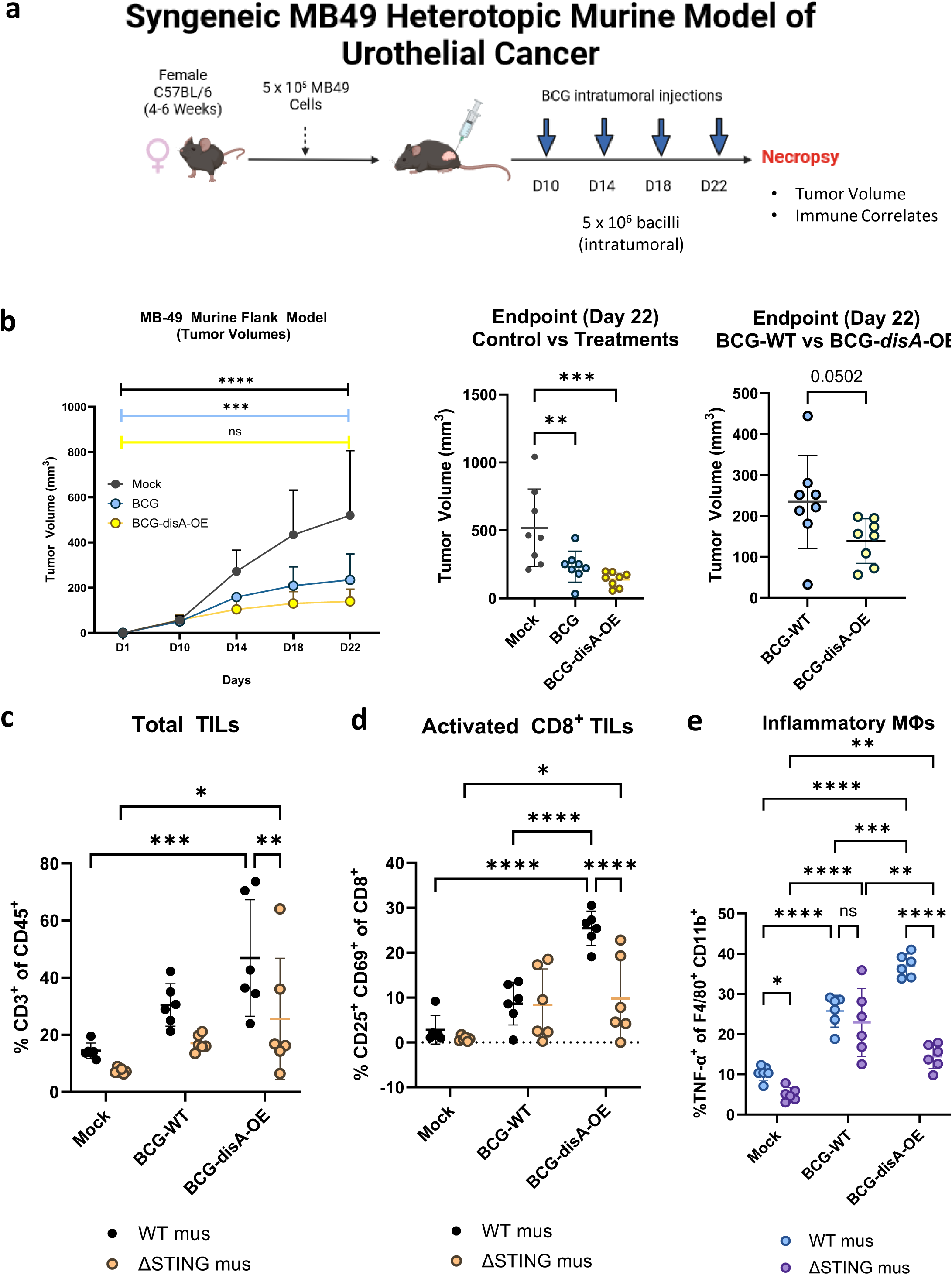
BCG-*disA*-OE elicits improved antitumor efficacy over BCG-WT in the syngeneic MB49 heterotopic mouse model of urothelial cancer. **a.** Schematic diagram of the MB49 syngeneic mouse model of urothelial cancer. **b.** MB49 tumor volumes and at time of necropsy on day 22 (8 animals/group). **c.** Tumor infiltrating lymphocytes (TILs, percent CD3+ of CD45) at necropsy, **d.** Activated CD8+ TILs (percent CD25+ CD69+ of CD8+), and e. inflammatory macrophages (percent TNFα+ of F4/80+ CD11b+). The flow cytometry experiments were performed with treatment on days 10, 14, 17, and 21, with necropsy on day 22 as shown in Fig. **S3a** (6 animals/group). Data are represented as mean values ± S.D. Statistical analyses done using one-way ANOVA with Dunnett’s multiple comparisons test in panel **b** (control vs treatments); two-tailed student’s T-test in panel b (BCG-WT vs BCG-*disA*-OE); two-way ANOVA with Tukey’s multiple comparisons test in panels **b, c, d,** & **e** (* p < 0.05, ** p < 0.01, *** p < 0.001, ****p < 0.0001).

We further characterized the impact of the treatments on recruitment of activated T cells and macrophage polarization and in the tumor microenvironment (TME) using the MB49 model. Compared with BCG-WT, BCG-*disA*-OE significantly increased the abundance of total tumor infiltrating lymphocytes (TILs) (**Fig 3c**), activated (CD25+, CD69+ CD8+) TILs (**Fig. 3d**) in the MB49 model. We also observed T cells with other activation markers including IFNγ+ CD8 cells and CD69+ CD38+ CD8 cells more abundantly with BCG-*disA*-OE than with BCG-WT in the MB49 tumors. We also considered myeloid cells in the MB49 TME and observed that BCG-*disA*-OE elicited significantly higher numbers of inflammatory macrophages (TNF-α^+^, MHCII^+^) than BCG-WT (**Fig. 3e**, **S3b-e**). Similarly, TNF-expressing M2-like macrophages (CD206^+^, CD124^+^) were also more abundant with BCG-*disA*-OE than with BCG-WT. When we compared the ability of BCG-*disA*-OE to recruit each of these cell types to tumors we found the effect to be STING dependent with considerably lower percentages of cells being seen in STING knockout mice (**Fig. 3c-e**, **Fig. S3b-e**). Thus, BCG-*disA*-OE not only showed superior antitumor efficacy than BCG-WT in the MB49 model, but it also recruited greater percentages of activated lymphocytes and macrophages to the tumors.

### Safety studies: BCG-*disA*-OE is less pathogenic than BCG-WT in two mouse models

To address concerns that the enhanced pro-inflammatory immune responses elicited by BCG-*disA*-OE might lead to adverse effects, we evaluated safety in two separate mouse models. We used an immunocompetent BALB/c mouse model of aerosol exposure and measured the lung bacillary burden after four weeks when adaptive immune responses are maximal (**Fig. 4a**). While the day 1 implantation of the two BCG strains was equivalent at 2.6 log10 colony forming units (CFUs), we observed that BCG-*disA*-OE (Tice) proliferated in murine lungs to a significantly lower degree than BCG-WT (Tice) by a margin of 0.43 log10 CFUs at 4 weeks post-challenge (**Fig. 4b**). This same experiment performed with BCG-*disA*-OE (Pasteur) versus BCG-WT (Pasteur) gave virtually identical results (**Fig. S4a-b**). We also tested the two strains in immunocompromised SCID mice for which infection with BCG leads to fatal systemic disease (**Fig. 4c**). Using a low dose aerosol exposure model that implanted 1.1 log10 colony forming units in the lungs (**Fig. 4d**), we observed a statistically significant survival prolongation with a mean time to death for BCG-*disA*-OE (Tice) of 148 days compared to 112 days for BCG-WT (Tice) (**Fig. 4e**). Virtually identical results were obtained for BGC-*disA*-OE (Pasteur) versus BCG-WT (Pasteur) in the SCID mouse time-to-death experiment (**Fig. S4c-d**). Thus, despite eliciting more profound inflammatory signatures in numerous model systems, BCG-*disA*-OE is less pathogenic than BCG-WT in these two murine model systems.

**Figure 4.**
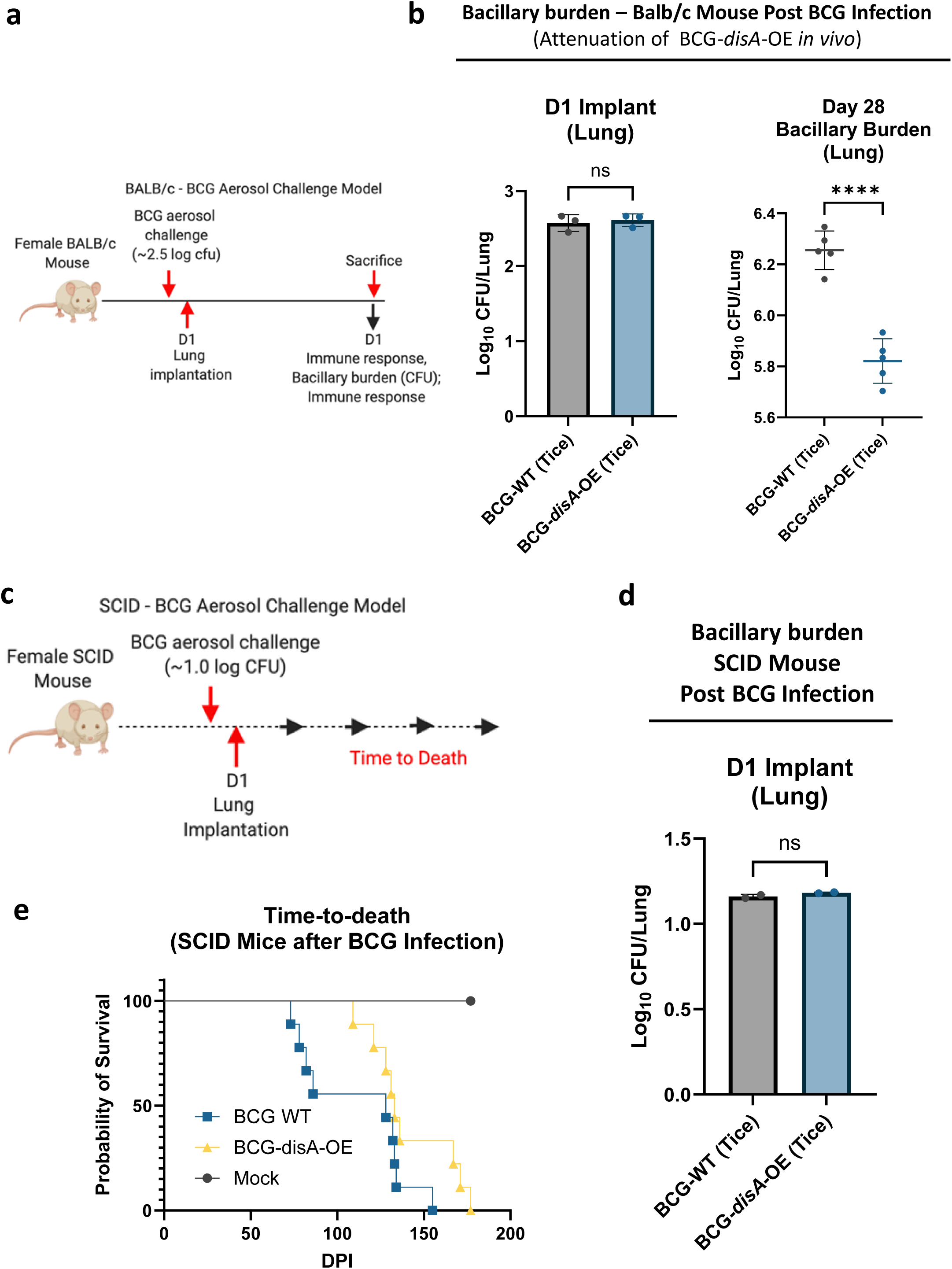
BCG-*disA*-OE is less pathogenic than BCG-WT in two mouse models. **a.** Schematic diagram of the immunocompetent BALB/c mouse challenge model. **b**. BALB/c lung colony forming unit (CFU) counts at day 1 (n= 3 animals/group) and day 28 (n= 5 animals/group). **c.** Schematic diagram of the immunocompromised SCID mouse challenge model. **d.** SCID mouse lung colony forming unit (CFU) counts at day 1 (n= 2 animals/group). **e.** Percent survival of SCID mice following low dose challenge (n=10 animals/group). The experiment was performed with BCG strains in the Tice background. Similar results were obtained using the Pasteur background as shown in **Fig. S4**. Data are represented as mean values ± S.D. Statistical analyses done using 2-tailed Student’s t-test in panels **b** and **d**, Kaplan-Meier survival curve in panel **e** (**** p < 0.0001).

### BCG-*disA*-OE elicits greater interferon-β (IFN-β) response than BCG-WT in primary murine and human macrophages *in vitro*

To better characterize the nature of the immune responses elicited by BCG-*disA*-OE and its BCG-WT parent strain, we exposed monocytes or macrophages to the BCG strains and characterized their phenotypic responses. First, we studied a recombinant reporter cell line (RAW Lucia ISG cells) that gives a luminescence signal upon activation of the STING pathway leading to interferon-stimulated gene (ISG) up-regulation. Compared to BCG-WT, BCG-*disA*-OE significantly increased activation of the STING pathway in RAW Lucia ISG macrophages as measured by relative light unit induction (**Fig. 5a**, **Fig. S1b**). Since STING pathway activation is associated with strong up-regulation of Type I interferon responses, we next characterized the interferon-β (IFN-β)-inducing potential of BCG-*disA*-OE versus BCG-WT. Next, we evaluated induction of IFN-β expression in primary murine bone marrow-derived macrophages (BMDM), a murine macrophage cell line (J774.1) and in primary human monocyte-derived macrophages (HMDMs). We found consistent induction of IFN-β in all myeloid cell types in response to BCG-*disA*-OE that was siginficantly higher than that seen with BCG-WT-exposed cells (**Fig. 5b-c**). The increased expression of IFN-β by BCG-*disA*-OE over BCG-WT was strictly STING-dependent as confirmed using BMDM from STING^-/-^ mice (ΔSTING) (**Fig. 5b**). This effect was greatly potentiated when pre-treated with IFN-γ (**Fig S5a-c**)

**Figure 5.**
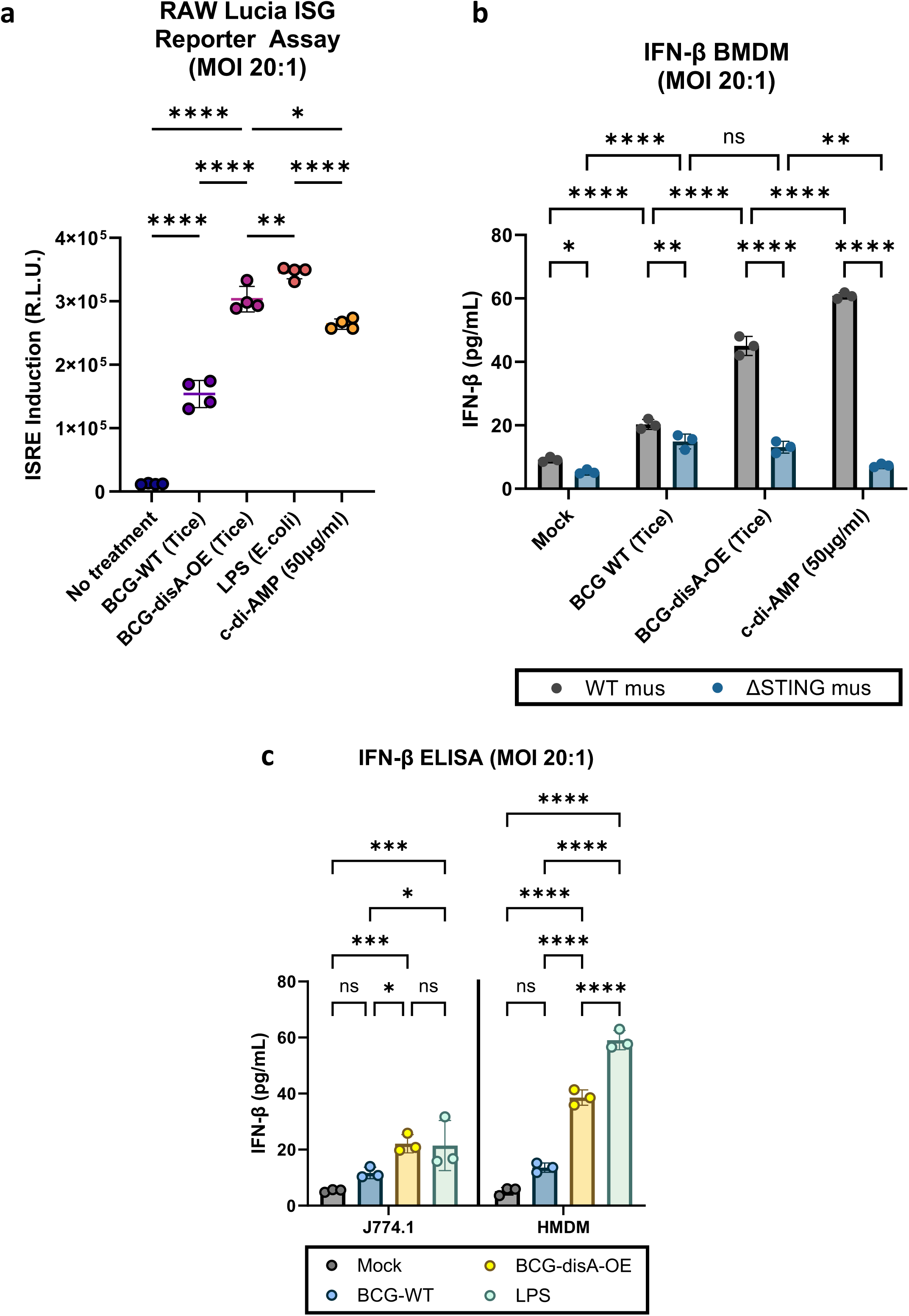
BCG-*disA*-OE elicits greater interferon-β (IFN-β) responses than BCG-WT in primary murine and human macrophages *in vitro*. **a.** IRF3 induction measured in RAW-Lucia ISG reporter murine (Balb/c) macrophages. **b.** IFN-β levels in murine BMDM from wild-type and STING^-/-^ mice (C57BL/6 background). **c.** IFN-β levels in J774.1 macrophages and human monocyte-derived macrophages (HMDM) following exposure to BCG strains. Cytokine levels were measured by ELISA after 24 hours exposure at and MOI of 20:1. Data are presented as mean values ± SD (n=3 biological replicates). Statistical analyses done using one-way ANOVA w/Tukey’s multiple comparisons test in panel a; two-way ANOVA with Tukey’s test for multiple comparisons in panels **b** and **d** (* p < 0.05, ** p < 0.01, *** p < 0.001, **** p < 0.0001).

### M1 Pro-inflammatory polarization of macrophages is greater with BCG-*disA*-OE than with BCG-WT

Trained immunity is associated with polarization of macrophages towards inflammatory phenotypes with a concomitant shift away from anti-inflammatory states.^23^ To investigate macrophage polarization, we used flow cytometry to monitor phenotypic shifts of both murine and human primary macrophages following a 24 h exposure to BCG-*disA*-OE or BCG-WT (gating strategies shown in **Fig. S6-S9**). First, we focused on the MHC class II expressing CD11b^+^ F4/80^+^ murine BMDM population following *in vitro* BCG exposure as shown in **Fig. 6a**. We observed a trend towards greater expansion of TNF-α-expressing MHCII+ CD11b^+^ F4/80^+^ inflammatory murine BMDMs (M1-like) following exposure to BCG-*disA*-OE than with BCG-WT. We next gated cells expressing the immunosuppressive surface receptors CD206^+^ and CD124^+^ among CD45^+^ CD11b^+^ F4/80^+^ macrophages and observed a significant reduction of this M2-like population with BCG-*disA*-OE than with BCG-WT (**Fig. 6b**). Within this M2-like immunosuppressive cell population, there was a higher proportion of IL-10-expressing CD206^+^ CD124^+^ cells in BCG-WT-exposed macrophages, while IL-10-expressing cells were significantly reduced in response to BCG-*disA*-OE exposure (**Fig. 6c**). These results demonstrate that compared with BCG-WT, BCG-*disA*-OE exposure elicits more extensive macrophage reprogramming with expansion of pro-inflammatory macrophages displaying increased antigen presentation (MHC class II expression) and TNF-α expression and contraction of immunosuppressive macrophages expressing IL-10.

**Figure 6.**
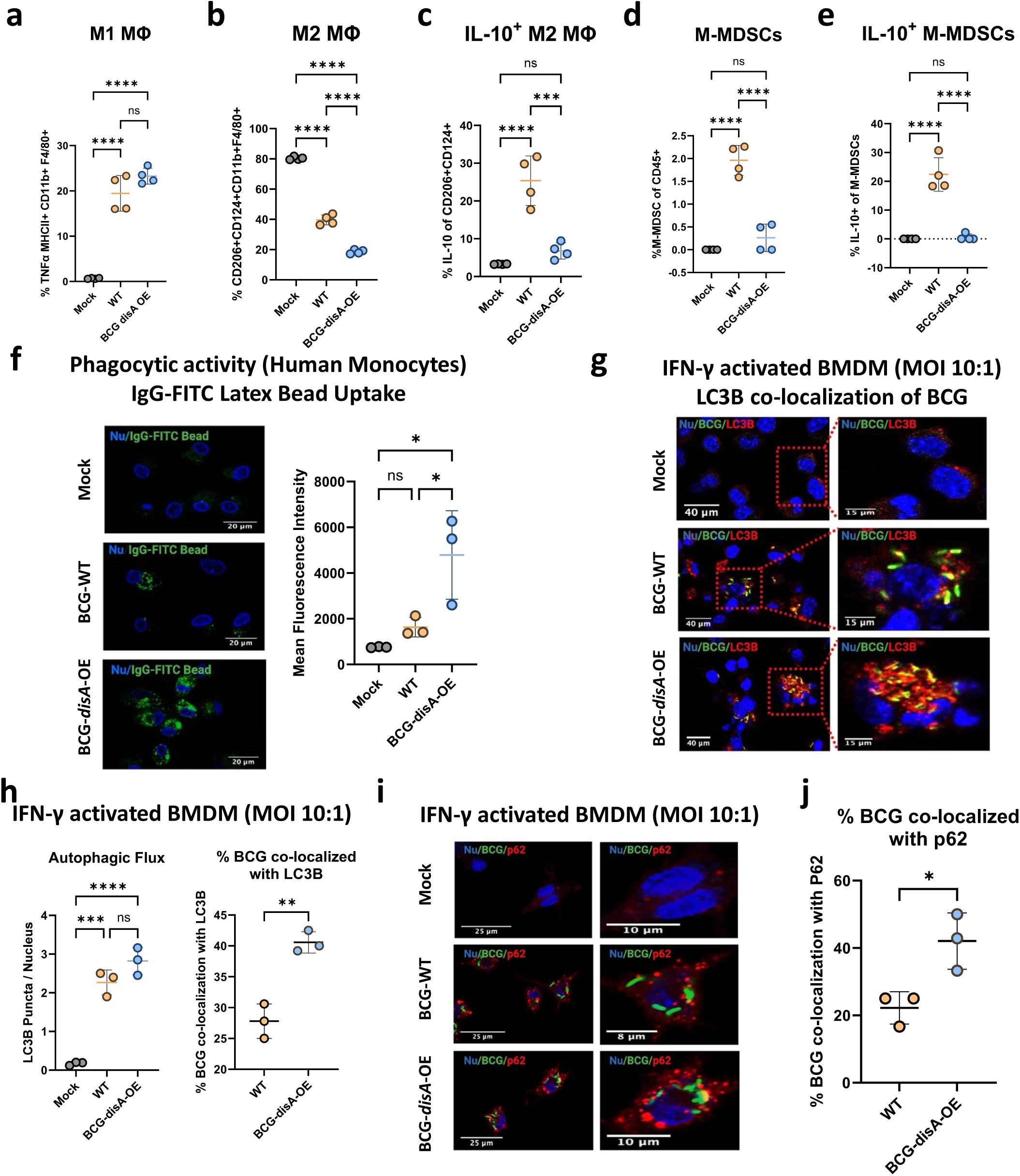
BCG-*disA*-OE elicits greater macrophage re-programming, phagocytic activity, and autophagy than BCG-WT in human and murine macrophages. Percentages of cells arising from primary murine macrophages exposed to BCG (Tice) strains at an MOI of 20:1 at 24 hours post-exposure: **a.** M1-like macrophages (TNFα-expressing of MHCII+ CD11b+ F4/80+ cells), **b.** M2-like macrophages (CD206-, CD124-expressing of CD11b+ F4/80+ cells), **c.** IL-10 expressing M2-like macrophages (IL-10 expressing of M2-like macrophages), **d.** monocytic myeloid-derived macrophages (M-MDSCs, Ly6C^hi^, Ly6G-of CD11b+ F4/80+ cells), **e.** IL-10 expressing M-MDSCs (IL-10-expressing of M-MDSCs) (FigS9b). Flow cytometry studies shown are for BCG strains in the Pasteur background. Data are presented as mean values ± SD (n = 3 biological replicates). Gating schemes and data acquisition examples are shown in **Fig. S6-S9**. **f.** Phagocytic activity in human primary macrophages in representative confocal photomicrographs showing intracellular uptake of FITC-labeled IgG-opsonized latex beads (green) with nuclei stained blue. **g.** Autophagy induction measured by LC3B puncta co-localization with BCG strains, and h. quantification of BCG-LC3B co-localization in primary murine macrophages shown by representative confocal photomicrographs. **i.** Autophagy induction measured by p62 puncta co-localization with BCG strains and p62, and **j.** quantification of BCG-p62 co-localization. FITC-labeled BCG strains are stained green, LC3B or p62 autophagic puncta (red), nuclei blue, and co-localization (yellow). Cells were fixed using 4% paraformaldehyde 6 h after infection (MOI 10:1), and images obtained with an LSM700 confocal microscope and Fiji software processing. Quantification was measured by mean fluorescence intensity. Co-localization studies shown are for BCG strains in the Tice background. Data shown for the confocal microscopy studies are mean values ± SD (n= 3 biological replicates). Statistical analyses done using one-way ANOVA w/Tukey’s multiple comparisons test in panels **a-f**, & **h**; 2-tailed Student’s t-test in panels **h** and **j** (* p < 0.05, ** p < 0.01, *** p < 0.001, **** p < 0.0001).

Myeloid-derived suppressor cells (MDSCs) are a heterogeneous population of immature myeloid cells known to foster immunosuppression.^24,25^ Accordingly, we investigated the induction of monocytic-myeloid derived suppressor cells, M-MDSCs, (Ly6C^hi^ Ly6G^-^ CD11b^+^ F4/80^-^) using primary murine BMDMs. Following BCG-WT exposure we observed a significant expansion of M-MDSCs, while in contrast this same population showed minimal expansion following BCG-*disA*-OE exposure (**Fig. 6d**). Moreover, the M-MDSCs elicited by BCG-WT exhibited higher IL-10 expression, whereas IL-10-expressing M-MDSCs were virtually absent after BCG-*disA*-OE exposure (**Fig. 6e**). These observations suggest that BCG-WT contributes to an expansion of M-MDSCs which have immunosuppressive properties; however, forced overexpression of the pro-inflammatory PAMP cyclic di-AMP by BCG prevents M-MDSC expansion.

### Macrophages exposed to BCG-*disA*-OE are more phagocytic than those with BCG-WT

Cyclic dinucleotides have been reported to recruit inflammatory macrophages which display high phagocytic potential.^26–29^ Consistent with these observations, we confirmed that HMDMs transfected with cyclic di-AMP showed increased phagocytosis and exhibited elongated dendrites compared to mock-transfected populations (**Fig. S10)**. We then evaluated the phagocytic properties of HMDMs following exposure to the different BCG strains and found significantly greater phagocytosis of IgG-opsonized FITC-latex beads by macrophages exposed to BCG-*disA-*OE compared to BCG-WT (**Fig. 6f**). In keeping with the previously established role of STING pathway activation in augmenting autophagy,^15,30,31^ we found that a significant majority of intracellular BCG-*disA*-OE bacilli were co-localized with LC3B in IFN-γ-activated primary BMDMs (**Fig. 6g-h**), while autophagy induction in BCG-WT was significantly lower. We also found significantly greater co-localization of BCG-*disA*-OE bacilli with the autophagy adapter protein p62 compared to that observed with BCG-WT (**Fig. 6i-j**). To test whether similar autophagic targeting effects might occur in non-immune cells, we tested exposed 5637 human urothelial carcinoma cells to the BCG strains. Similar to our observations with HMDMs, this cancer cell line also displayed autophagic targeting of intracellular BCG, with BCG-*disA*-OE being a significantly more potent inducer of co-localization with LC3B puncta than BCG-WT (**Fig. S11**) These results reveal BCG-*disA*-OE increases the levels of phagocytosis and autophagic processing within macrophages to a greater degree than BCG-WT, a phenomenon associated with enhanced peptide antigen presentation to MHC class-II molecules.^32,33^

### BCG-*disA*-OE reprograms macrophages epigenetically and potentiates trained immunity to a greater degree than BCG-WT

Recent studies indicate that BCG exerts its antitumor effects via a trained immunity mechanism.^11^ In light of recent data showing BCG to be a potent inducer of trained immunity through epigenetic modifications of key pro-inflammatory genes,^34–36^ we hypothesized that the addition of cyclic di-AMP overexpression to standard BCG might potentiate epigenetic modifications in primary human monocytes. First, we tested levels of TNF-α and IL-6 secretion by human monocytes exposed to the BCG strains. Using monocytes from healthy human subjects, we observed that BCG-*disA*-OE elicited higher cytokine secretion levels than BCG-WT as measured by RT-qPCR (**Fig. 7a**). The ability of traditional BCG to elicit trained immunity has been correlated with changes in epigenetic marks that increase pro-inflammatory gene expression.^37^ Thus, we asked if the enhanced induction of TNF-α and IL-6 expression elicited by BCG-*disA*-OE compared with BCG-WT is epigenetically mediated (**Fig. 7a-b**). To this end, we evaluated the promoter regions of the IL-6 gene for durable, antigen-independent epigenetic changes using an assay in which human monocytes exposed to BCG strains for 24 h were rested for five days prior to challenge with a heterologous antigen, the TLR1/2 agonist Pam3CSK4 on day 6 (**Fig. 7b**).^38^ Using chromatin immunoprecipitation-polymerase chain reaction (ChIP-PCR) assays, we quantified the activating histone methylation mark H3K4me3 present in the IL-6 promoter. We observed that exposure to BCG-*disA*-OE led to greater enrichment of this mark than BCG-WT even without the heterologous second stimulation (i.e., adding RPMI media alone at day 6). Upon secondary-stimulation with Pam3CSK4 at day 6, the abundance of the activating epigenetic mark was further increased by both BCG strains, but BCG-*disA*-OE-pretreatment yielded notably more enrichment than BCG-WT (**Fig. 7c**). Simultaneous measurement of IL-6 (as well as TNF-α) in BCG-trained culture supernatant following non-specific stimulation by Pam3CSK4 revealed that BCG-*disA*-OE-trained macrophages produced significantly higher levels of these pro-inflammatory cytokines than did those trained with BCG-WT (**Fig. 7d-e**). These results indicate that an augmented BCG which overexpresses the PAMP molecule, cyclic di-AMP, leads to significantly more robust epigenetic changes classically associated with trained immunity.

**Figure 7.**
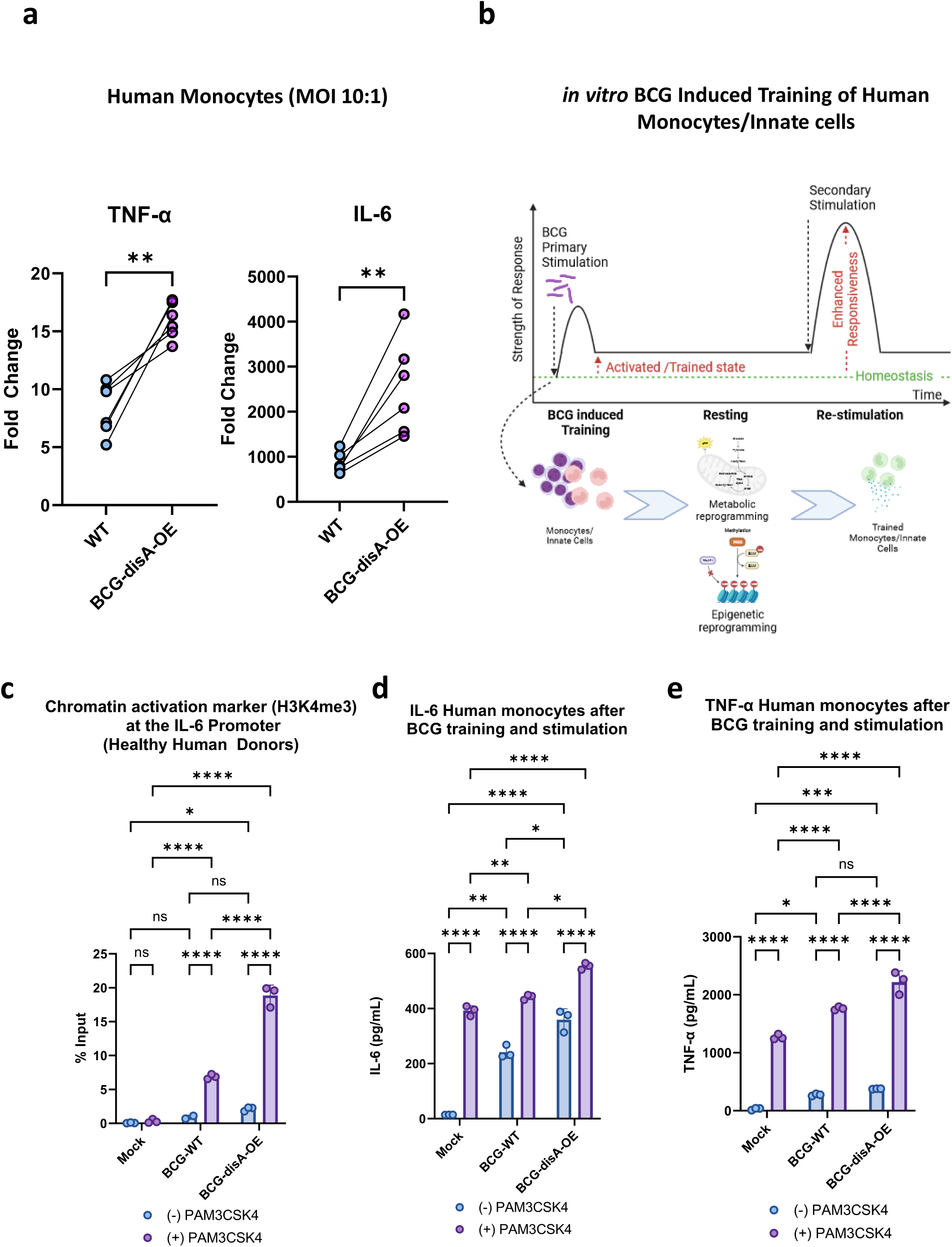
Compared with BCG-WT, BCG-*disA*-OE is a more potent inducer of epigenetic changes characteristic of trained immunity in primary human monocytes. **a.** Fold change in mRNA levels of TNF-a and IL-6 in primary human monocytes (n=6 healthy donors) relative to the RNU6A transcript after 24 hr exposures at a MOI of 10:1. **b.** Schematic diagram of *in vitro* monocyte training. **c.** Relative levels of the H3K4me3 chromatin activation mark in the IL-6 promoter region in the primary human monocytes of one healthy donor determined by ChIP-PCR assay on day 6 following initial stimulation on day 0 with no treatment (NT) or one of the BCG strains and a second stimulation on day 6 with NT or the TLR1/2 agonist PAM3CSK4. **d.** Secreted levels of the cytokine IL-6 and **e.** TNF-a from primary human monocytes (3 healthy donors) following BCG training and re-stimulation by the same protocol. Monocytes were initially challenged on day 0 with a 24 hr exposure to the BCG strains at a MOI of 10:1 followed by washing. After 5 days of rest, they were treated for 24 hr with either no treatment (RPMI) or the TLR1/2 agonist Pam3CSK4. Data are represented as mean values ± SD (n= 3 biological replicates). Statistical analyses done using a paired 2-tailed Student’s t-test on panel A; two-way ANOVA w/Tukey’s multiple comparisons test on panels **c, d,** & **e** (* p < 0.05, ** p < 0.01, *** p < 0.001, **** p < 0.0001).

### BCG-*disA*-OE is a potent inducer of macrophage immuno-metabolic reprogramming towards pro-inflammatory signatures to a greater degree than BCG-W

BCG-training has been reported to stimulate glycolysis as well as the tricarboxylic acid cycle through glutamine replenishment with accumulation of fumarate.^39^ To address whether the addition of cyclic di-AMP overexpression alters the BCG-mediated metabolomic shifts, we used LC-MS to characterize key metabolites in primary human and murine macrophages exposed to the two BCG strains. As shown in **Fig. 8a-b**, HMDMs or BMDMs exposed to BCG-*disA*-OE for 24 h showed significantly increased catabolic signatures of intracellular glucose (p < 0.01) and lactate (p < 0.05) to a significantly greater degree than with BCG-WT. Also, the TCA cycle metabolites itaconate and fumarate were also more elevated with BCG-*disA*-OE than with BCG-WT (**Fig. 8c**). These observations suggest that glycolytic carbon substrates for ATP generation (consistent with a pro-inflammatory bioenergetic profile) accumulate to a greater in macrophages infected with BCG-*disA*-OE than with BCG-WT.^37^

**Figure 8.**
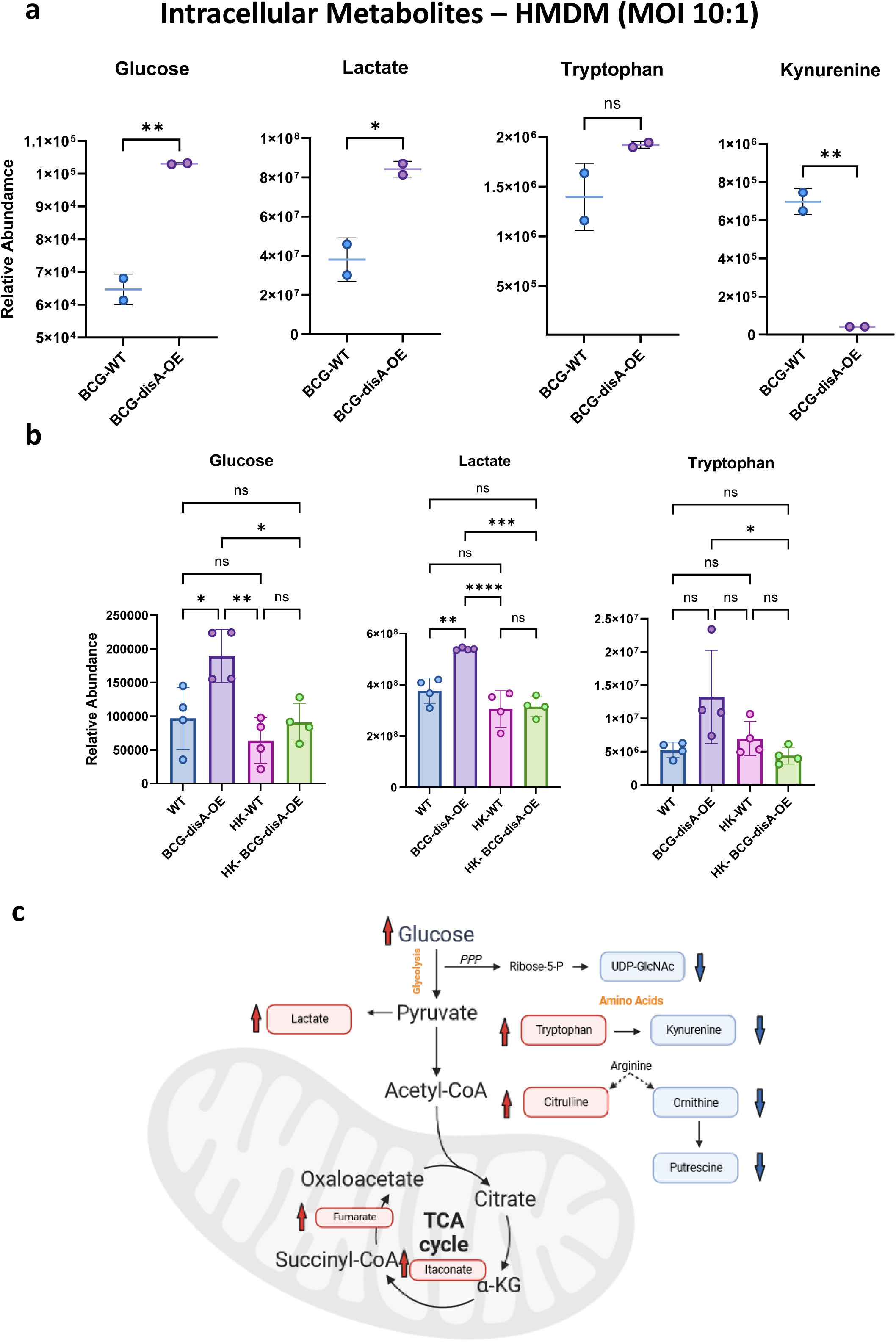
Compared with BCG-WT, BCG-*disA*-OE is a more potent inducer of metabolomic changes characteristic of trained immunity in primary human monocytes. **a-b.** Metabolite levels determined by LCMS in human or murine MDM determined 24 hr after exposure to BCG (Tice) strains or heat-killed (HK) controls. Schematic diagram **(c)** showing key metabolites significantly upregulated (red arrow upward) or downregulated (blue arrow downward) in BCG-*disA*-OE infected macrophages relative to BCG-WT infected macrophages. Data are represented as mean values ± SD (n= 4 biological replicates) Statistical analyses done using two-tailed Student’s t-test on panel **A**; one-way ANOVA w/Tukey’s test for multiple comparisons in panel **b** (* p < 0.05, ** p < 0.01, *** p < 0.001, **** p < 0.0001).

To determine whether the elevated levels of intracellular glucose were due to increased transport or to increased gluconeogenesis, we tested expression of the major glucose transporter GLUT1 in macrophages exposed to BCG-WT and BCG-*disA*-OE. As shown in **Fig. S12a**, we challenged murine BMDM with BCG or BCG-*disA*-OE for 4 hours, and after washing and recovery in glucose-free media, macrophages were treated with the fluorescent 2-deoxy-glucose analogue 2-NBDG for 2 hours and subsequently analyzed by flow cytometry for levels of GLUT1 expression or 2-NBDG uptake. As may be seen in **Fig. S12b**, exposure to BCG-WT and BCG-*disA*-OE led to a 2-fold and more than a 5-fold increase, respectively, of GLUT1 expression on BMDM compared to unexposed cells. Similarly, 2-NBDG levels were elevated by 20% or 40% following exposure to BCG-WT and BCG-*disA*-OE, respectively as shown in **Fig. S12c**. These observations strongly suggest that BCG-*disA*-OE elicits higher levels of the GLUT1 transporter and glucose uptake than BCG-WT or untreated controls resulting in greater accumulation of intracellular glucose and are consistent with earlier observations linking trained immunity and STING activation with enhanced mTOR-HIF-1α pathway activation and concomitant elevations in glucose transporter levels.^40–42^

Excess tryptophan catabolism to kynurenine by tryptophan dehydrogenase and indoleamine 2,3-dioxygenase (IDO) has been strongly associated with immunosuppression,^43^ and IDO inhibitors have shown potential as immune activators in a variety of infectious and oncologic diseases.^44^ Kynurenine levels were dramatically lower in macrophages following BCG-*disA*-OE exposure than those seen with BCG-WT (**Fig. 8a**), and as would be expected, tryptophan levels were elevated by BCG-*disA*-OE while BCG-WT led to tryptophan levels comparable to the baseline seen with heat-killed BCG controls (**Fig. 8b**). Citrulline levels were also higher while putrescine levels were lower with BCG-*disA*-OE than BCG-WT suggesting that nitric oxide synthase-mediated conversion of arginine to NO (pro-inflammatory) and citrulline was more strongly induced by BCG-*disA*-OE (**Fig. 8c**). Finally, it was of interest that itaconate, an isocitrate lyase inhibitor made by macrophages that has been shown to have antibacterial activity, was more potently induced by BCG-*disA*-OE than BCG-WT (**Fig. 8c**). Thus, compared with BCG-WT, BCG-*disA*-OE elicited a greater pro-inflammatory metabolomic signature with reduced kynurenine accumulation and increases in glycolytic metabolites, NOS products, and itaconate production.

## DISCUSSION

With high rates of recurrence or treatment failure in NMIBC even among patients who receive intravesical immunotherapy with BCG, there is an unmet need for improved NMIBC therapies.^2^ Indeed, several innovative therapies have been recently approved or are on the horizon for approval for the management of BCG-unresponsive NMIBC. These include the recently licensed nadofaragene firadenovec (adenoviral-based gene therapy),^45^ and several agents in late stage clinical trials including the oncolytic virus cretostimogene grenadeorepvec (CG0070),^46^ and a recombinant BCG agent known as VPM1002BC.^47^ VPM1002BC (also known as BCGΔ*ureC*::*hly*) is an rBCG strain that enhances phagosome permeability leading to exposure of BCG antigens to cytosolic MHC class I antigen processing,^48^ and it is currently in late stage clinical trials for tuberculosis and bladder cancer.^49^ In contrast to VPM1002BC and numerous other rBCG agents proposed for bladder cancer immunotherapy over the years which are designed to make recombinant proteins,^50^ BCG-*disA*-OE is an rBCG which is specifically re-engineered to overexpress a small molecule (c-di-AMP) which serves as a STING agonist with anticancer properties. In addition, owing to the known ability of BCG to persist in the phagosomal compartment of myeloid cells,^51^ BCG-*disA*-OE is likely to release the small molecule STING agonist endogenously in a sustained manner from the intracellular space of phagocytic cells in the urothelium.

In this study we evaluated BCG-*disA*-OE in two separate models of urothelial cancer. In the orthotopic, carcinogen-induced MNU rat model of NMIBC, we found that the highest degree of bladder cancer in BCG-*disA*-OE-treated animals was Ta with 53% of rats being cancer-free (p < 0.05); while, in contrast, invasive tumors developed in untreated rats (highest tumor grade of T2) and in BCG-WT-treated animals (highest tumor grade of T1). We also tested efficacy in the heterotopic, syngeneic mouse MB-49 bladder cancer model and found that BCG-*disA*-OE was superior to BCG-WT in reducing endpoint tumor volumes. These antitumor properties were accompanied by significantly higher recruitment of tumor infiltrating lymphocytes (TILs), activated CD8+ T cells, and inflammatory macrophages in of BCG-*disA*-OE-treated animals as compared to BCG-WT and untreated controls. In STING^-/-^ mice these cellular recruitment changes were reduced indicating that the activity of BCG-*disA*-OE is STING-dependent.

While the safety of BCG is well-established owing to its introduction as a TB vaccine in 1921 and as a bladder cancer immunotherapy in the mid-1970s, we were concerned that overexpression of the pro-inflammatory STING agonist might lead to unwanted toxicity. Hence, we tested the ability of BCG-*disA*-OE to proliferate in the tissues of immunocompetent mice and to lead to lethality in SCID mice, and by both measures we found that BCG-*disA*-OE was less pathogenic than BCG-WT. This is likely due to the fact that c-di-AMP is recognized as a PAMP by the innate immune system, and therefore cell-mediated immune responses that clear pathogenic mycobacteria are more strongly activated with BCG-*disA*-OE than BCG-WT. Indeed, related study from our lab found that *disA* overexpression in *Mycobacterium tuberculosis* was also associated with reduced pathogenicity compared to the wild-type.^15^

In recent years, BCG has been studied for its ability to elicit heterologous immunity to unrelated antigenic stimuli. This phenomenon, known as trained innate immunity, may account for the association of BCG vaccination with reduced rates of viral infections and childhood mortality.^52–54^ Cell-based studies of BCG-vaccinated human volunteers have documented the fact that BCG elicits epigenetic and metabolic reprogramming in myeloid cells.^39,55,56^ In this study we evaluated the comparative abilities of BCG-*disA*-OE and BCG-WT to stimulate trained immunity epigenetic responses (activation marks in the IL-6 promoter) and metabolic response (shift towards glycolysis), and found that BCG-disA-OE was a more potent stimulus than BCG-WT. These findings suggest that increased engagement of the STING pathway may be linked to trained immunity.

Our results demonstrate that BCG-*disA*-OE—an rBCG strain which overexpresses a small molecule STING agonist--leads to improved antitumor efficacy in two animal models of bladder cancer compared with BCG-WT, and that the elevated levels of STING agonist release does not result in higher BCG pathogenicity. Immune parameters characteristic of trained innate immunity are also more strongly induced by BCG-*disA*-OE than by BCG-WT. Overexpression of small molecules associated with antitumor activity may be a novel route towards re-engineered recombinant BCG strains with improved efficacy in bladder cancer.

## ACKNOWLEDGEMENTS

The authors gratefully acknowledge the financial support of NIH AI 37856, Tedco awards MII-4181, MII-5072, and awards from the Willowcroft Foundation and the Cigarette Restitution Fund of Maryland. The authors also thank Alok Singh, Gregory Joice, Manish Gupta and Emily Juzwiak for experimental assistance.

## AUTHOR CONTRIBUTIONS

W.R.B., and T.J.B. co-led the study through conceptualization, design, oversight, and the interpretation of results. W.R.B. and T.J.B. obtained funding for the study. P.K.U., M.P., K.A.L., T.Y., A.M., A.S.B., L.Z., G.S., J.H. and P.P. designed, conducted the experiments, and/or interpreted the results. M.L.K., D.M. and D.M.P. assisted in the design of experiments and provided key expert advice. P.K.U., G.S., W.R.B., and T.J.B. wrote the manuscript. P.K.U., M.P., K.A.L., T.Y., A.M., A.S.B., L.Z., G.S., P.P., M.L.K., D.M., D.M.P., W.R.B., and T.J.B. revised and edited the manuscript. P.K.U., W.R.B., and T.J.B. designed and produced figures for this manuscript.

## METHODS

Ethics: All protocols involving animals strictly adhered to US NIH guidelines and were approved by the Johns Hopkins Medical Institutions Animal Care and Use Committee under the protocols: MO18M58, MO20M20 and RA17M332.

### Bacterial strains and culture conditions

In this study we used *Mycobacterium bovis* (*M. bovis*) Bacillus Calmette-Guérin (BCG) Pasteur (BCG-WT Pasteur) (<u>a generous gift from Dr. Frank Collins [FDA]</u> and identical to BCG-Pasteur provided by the Pasteur Institute to the Trudeau Institute in 1967 as TMC No. 1011) and commercially available BCG-Tice (Onco-Tice^©^, Merck) for generation of c-di-AMP overexpressing recombinant BCG strains. Briefly, genomic DNA from *Mycobacterium tuberculosis* (*M. tb*) strain CDC1551 was used for PCR amplification of *disA* (MT3692/Rv3586). Single isolated bacterial colonies growing on 7H11 plates supplemented with oleic-albumin-dextrose-catalase (OADC) (Cat. B11886, Fisher Scientific) were picked and propagated in 7H9 Middlebrook liquid medium (Cat. B271310, Fisher Scientific) supplemented with (OADC) (Cat. B11886, Fisher Scientific), 0.5% glycerol (Cat. G55116, Sigma) and 0.05% Tween-80 (Cat. BP338, Fisher Scientific). Cloning experiments were performed using *E. coli* strain DH5-α (Cat. 18258012, Fisher Scientific) and was routinely maintained in LB broth. For generation of *disA* overexpressing BCG, an *E. coli*-mycobacterial shuttle vector (pSD5.hsp60) was used to clone *M.tb* gene MT3692 or Rv3586 under the strong mycobacterial promoter *hsp60*^18^. Clones were confirmed by gene sequencing and were used for bacterial transformation by electroporation method. Recombinant strains were confirmed using colony PCR against kanamycin cassette, subjected to whole genome sequencing and qPCR analyses. Details of all bacterial strains, plasmids and constructs are listed in supplementary **Table S1**.

### Mammalian cell culture

#### Cell lines

For cell-based *in vitro* infection assays J774.1 (American Type Culture Collection-ATCC® TIB67™, Manassas, VA, USA) murine macrophage cell lines were cultivated in RPMI-Glutamax (Cat. 61870-036, Fischer Scientific), supplemented with 10% heat inactivated fetal bovine serum (FBS) (Cat. 10082147, Fischer Scientific) with 1% streptomycin/penicillin at 37°C with 5% CO2. The urothelial carcinoma cell line 5637, a human high grade urothelial cancer (obtained from ATCC, HTB-9™) and MB49 cells (murine urothelial carcinoma cells, 7,12-dimethylbenz[a]anthracene, DMBA, EMD Millipore, Cat. SSC148) were maintained as monolayers in RPMI1640 medium supplemented with 10% heat inactivated fetal bovine serum (FBS) with 1% streptomycin/penicillin at 37°C with 5% CO2. The mouse fibroblast cell line NCTC clone 929 [L cell, L-929, derivative of Strain L] (ATCC® CCL-1™) was routinely maintained as monolayer in DMEM media supplemented with 10% heat inactivated fetal bovine serum (FBS) with 1% streptomycin/penicillin at 37°C with 5% CO2. All cell lines were maintained for fewer than 10 passage cycles and *Mycoplasma* testing was performed periodically while cells were in culture. The reporter mouse cell line, RAW-Lucia ISG (InvivoGen, CA, USA) was cultivated in custom prepared media as per manufacturer’s instructions.

#### Primary cells (Macrophages and Dendritic Cells)

For generation of murine bone-marrow-derived macrophages (BMDMs) and dendritic cells (BMDCs), bone marrow (BM) cells were isolated from 4-week old female wild-type (WT) C57BL/6J (Charles River laboratories, North Wilmington, Mass) and STING-KO mice (C57BL/6J-Tmem173gt/J, Jackson laboratories). The seed stock containing multiple vials of bone-marrow cells were preserved in cryopreservation media containing 10% DMSO (Cat. D2650; Sigma) and 90% heat inactivated FBS (Cat. 10082147, Fischer Scientific) in liquid nitrogen. For differentiation of BM cells into macrophages or DCs, random cryopreserved vials were chosen and differentiated for 6 days in BMDM-differentiation media made from DMEM containing 10% FBS, 1% MEM amino acids (Cat. 11130051, Thermo Fisher Scientific), 1% MEM non-essential amino acids (Cat. 11140050, Thermo Fisher Scientific), 1% sodium pyruvate (Cat. 11360070, Thermo Fisher Scientific), 1% MEM vitamin (Cat. 11120052, Thermo Fisher Scientific) and antibiotics (Penicillin-Streptomycin solution) supplemented with 30% sterile mouse fibroblast L929 (ATCC® CCL-1™) conditioned media. Differentiation of BM cells into DCs was carried out in low attachment 10 mm cell culture dish in presence of bone marrow-differentiation media in presence of recombinant murine Granulocyte-Macrophage Colony-Stimulating Factor (GM-CSF) (Cat. 315-03, Peprotech) for 48 **a.** h. Non-adherent cells were washed and loosely attached cells were allowed to differentiate into BMDCs for next 6 days. Cells were characterized for macrophage and DC markers using cell-surface staining and flow cytometry analyses. Human primary monocytes and human monocyte-derived macrophages (HMDMs) were used for cell-based *in viro* infection assays. Peripheral blood-derived mononuclear cells (PBMCs) isolated from healthy male donors (leukopacks) aged between 18-30 were used for isolation of human monocytes (HM) or human monocyte-derived macrophages (HMDM). Briefly, to separate blood constituents and isolation of buffy coat density gradient centrifugation (400 × *g* at 18°C for 30 min) of RPMI-1640 diluted blood over a Ficoll-Paque™ Plus reagent (Cat. 17-1440-02, GE Healthcare, Piscataway, NJ) was performed. Cells were washed several times using 1 x PBS and were counted using hemocytometer. Once counted CD14^+^ human monocytes were isolated from PBMCs using magnetic labeling (Monocyte Isolation Kit II, Cat. 130-091-153, Miltenyi Biotec, San Diego, CA) and magnetic columns as per manufacturer’s instructions. The purity of isolated CD14^+^ cells was confirmed using a fraction of cells stained with a fluorochrome-conjugated antibody against a monocyte marker as recommended by manufacturer and cells were analyzed using BD-LSR2 flow cytometer. Human monocytes were seeded (2.0 - 3.0 X 10^5^ cells / ml in RPMI 1640 medium supplemented with 10% FBS and 1% streptomycin/penicillin at 37°C with 5% CO2. Monolayers of CD14+ monocytes were differentiated into M1 [GM-CSF (20 ng/ml, PeproTech, Rocky Hill, NJ) and IFN-γ (20 ng/ml, PeproTech, Rocky Hill, NJ PeproTech)] or M2 [M-CSF (20 ng/ml, PeproTech, Rocky Hill, NJ) and IL-4 (20 ng/ml, PeproTech, Rocky Hill, NJ PeproTech)] for next 7 days.

#### Animals

Experimental procedures involving live animals were carried out in agreement with the protocols approved by the Institutional Animal Care and Use Committee (IACUC) at The Johns Hopkins University School of Medicine. For animal infection protocols, pathogen-free age 4-6 weeks female C57BL/6J (Charles River Laboratories, North Wilmington, Mass.), C57BL/6J-Sting1/J (STING^-/-^ Golden ticket mouse) (The Jackson Laboratory, ME, US), Fox Chase SCID mice (Charles River Laboratories North Wilmington, Mass.) and BALB/c mice (Charles River Laboratories, North Wilmington, Mass.) were purchased and housed under pathogen-free conditions at an Animal Biosafety Level-3 (ABSL3) or Biosafety Level-2 (ABSL2) animal facility without cross-ventilation. Fischer 344 female rats age 8 weeks (Harlan, avg. weight 160g) were housed at an BSL2 animal facility. Animals were housed under standard housing conditions (68-76°F, 30-70% relative humidity, 12-12 light-dark cycle) with free access to water and standard chow and were monitored daily for general behavior and appearance by veterinary specialists.

#### *In vitro* infection assays

For *in vitro* infection assays, cell lines or primary cells were seeded at required cell density in 6-well tissue culture plates or 10 mm petri dishes. For infection, log-phase wild-type and BCG-*disA*-OE strains were harvested by centrifugation and washed twice using DPBS to remove residual detergent and BSA then suspended in antibiotic-free RPMI 1640 media supplemented with 10% FBS. For infection assays, the bacteria were deposited at pre-calibrated multiplicity of infection (MOI). Infection was allowed for next 4 hours, followed by repeated washing of infected cells using warm DPBS to remove non-internalized bacteria. Infected cells were incubated until endpoints in presence of RPMI-1640 medium supplemented with 10% FBS and antibiotics.

#### Toxicity assays

The human urothelial cancer cell line 5637 was cultured at 37°C under 5% CO2 in RPMI 1640 containing 10% FBS without antibiotics. For cell toxicity assays 1500 5637 cells were seeded in a 96-well tissue-treated plate in triplicate, respectively. Twenty-four hours after seeding, cells were treated with the indicated ratio of BCG to cells for 72 hours. To measure cell viability, CellTiter-Glo Luminescent Cell Viability Assay (Promega, Madison, WI, USA) and FLUOstar OPTIMA (BMG Labtech, Ortenberg, Germany) were used according to manufacturer’s protocols. Relative cell viability was calculated by dividing the viability of the indicated ratio by that of a control.

For Annexin-PI staining, 0.5 million J774.1 cell and BMDMs were plated per well in 6-well plates for physical attachment. Cells were exposed at 1:10 MOIs for 24 hours using wild-type and BCG-*disA*-OE strains of Tice and Pasteur to determine the BCG cytotoxicity following exposure. At the endpoint of infection or treatment cells were non-enzymatically removed using 0.02% EDTA-PBS solution. Cells were washed twice with ice-cold PBS and FITC-annexin-PI was done as per manufacturer’s instruction using FITC Annexin V Apoptosis Detection Kit I (Cat. 556547, BD Biosciences). Flow cytometry was performed using a BD LSR II flow cytometer of the Flow Cytometry Core Facility at The Bloomberg School of Public Health, Johns Hopkins University). Data was processed using FACSDiva (v 9.0) and FlowJo (Tree Star v10) software.

#### Quantitative real-time qPCR

Gene expression profiling was carried out using total RNA isolated from cell lines or primary cells. For RNA isolation from rat bladders, pieces of whole bladder samples were excised, snap frozen in liquid nitrogen immediately after harvesting and stored in RNAlater (Cat. AM7021, Ambion) at -80°C. Total RNA isolation was carried out using RNeasy system (Cat. 74106, Qiagen). Real-time qPCR was performed using the StepOnePlus system (Applied Biosystems). For gene expression analyses in cell lines and primary cells, SYBR Fast green double stranded DNA binding dye (Cat. 4085612, Applied Biosystems) was used. Gene expression analyses in rat bladder tissues were performed using TaqMan gene expression assays. Gene-specific qPCR primers were purchased from Integrated DNA Technologies and all TaqMan gene expression assays were purchased from Thermo Fischer Scientific. Amplification of RNU6a, β-actin, GAPDH were used as endogenous control for RNA samples derived from human, mouse and rat cells/tissues respectively. All experiments were performed at least in triplicate and data analyses was done using 2^-ΔΔCT^ method. Details of NCBI gene identifiers and primer sequences are given in the Supplementary **Table S2**.

#### ELISA

Sandwiched ELISA was performed for cytokine (IFN-β, TNF-α, IL-6) measurements in culture supernatants. Briefly, culture supernatants were flash frozen in liquid nitrogen immediately after harvest and stored at -80 °C. Details of all ELISA kits and accessory reagents are given in supplementary table S2.

#### Multicolor confocal microscopy

Multicolor laser confocal microscopy experiments were performed to determine phagocytosis, autophagy, and colocalization studies in urothelial cancer cells and primary macrophages. Briefly, cells were allowed to adhere on sterile glass cover slips placed in 6-well tissue culture plates and infections were carried at pre-calibrated MOI. Log phase bacterial cultures were labeled using FITC (Cat. F7250, Sigma)^78^. Following infection and treatment conditions, cells were fixed, permeabilized and blocked followed by overnight incubation with a primary antibody for LC3B (Cat. NB100-2220, Novus) or p62/SQSTM1 (Cat. P0067, Sigma-Aldrich) at recommended dilutions at 4 °C. Cells were washed and incubated in the dark with Alexa Flour 647 conjugated secondary antibody (Cat. A32733, Thermo Fisher Scientific) at 4 °C for 1hour. DNA staining was carried out using Hoechst 33342 (Cat. 62249, Thermo Fisher Scientific) for 5 minutes. Images were acquired using Zeiss LSM700 single-point, laser scanning confocal microscope at 63X magnification at the Microscope Facility, Johns Hopkins School of Medicine. Image processing and analyses was carried out using open source Fiji software (https://imagej.net/software/fiji/). For LC3B or p62 quantification, perinuclear LC3B puncta (spot) was counted in a minimum 100 cells across different fields using and Imaris 9.5.0. Quantification carried out using GraphPad Prism (Prism 10.0.3) software.

#### Phagocytosis assay

IgG-FITC conjugated latex bead phagocytosis assay kit (Item No. 500290, Cayman Chemicals, USA) was used for phagocytosis studies. Briefly, HMDMs were placed on sterile glass cover slip for attachment. Infection was carried out at 5:1 (HMDM versus BCG) ratio for 3 hours followed by addition of IgG-FITC beads in warm RPMI 1640 media at 1: 400 dilutions for 3 hours. Nuclear staining was carried out using Hoechst 33342 (Cat. 62249, Thermo Scientific) and cells were visualized for bead phagocytosis using Zeiss LSM700 single-point, laser scanning confocal microscope. Quantification of beads was measured by mean fluorescence intensity (M.F.I.) calculations using open-source Fiji Software (https://imagej.net/software/fiji/).

#### Multicolor flow cytometry

The cell surface and intracellular staining was carried out on J774.1, murine BMDMs, human HMDMs and single cells derived from murine MB49 tumors and spleens. Flow cytometry panel were designed and if needed modified form murine myeloid and lymphoid cells and human myeloid cells. Details of all antibodies and the dilutions used are given in the supplementary table S2. For *in vitro* infection assays, protein transport inhibitor cocktail (Cat. 00-4980-03, eBioscience) at recommended dilution, 12 hours before harvesting monolayer of cells. At the endpoint cells were harvested using a cell-detachment buffer (ice-cold PBS - 10 mM EDTA solution). Single cell isolation was performed using animal tissues by harvesting tumors and spleens following necropsy. Briefly, tissues were manually disrupted before incubating in collagenase type I (Gibco) and DNase (Roche) in RMPI for 30 minutes at 37 °C. Tumor and spleen cells were dissociated through a 70-μm filter and washed with PBS. RBC lysis was performed for 5 minutes using ACK lysis buffer (Cat. A1049201, Thermo Fisher Scientific) at room temperature. Cells were washed twice using ice-cold PBS and stained using Zombie Aqua™ Fixable Viability Kit (Cat. 423101, Biolegend). Cells were washed and resuspended in FACS buffer (1% BSA, 2mM EDTA in PBS), Fc blocked (TruStain FcX™, Cat. 101320, and True-Stain Monocyte Blocker™ Cat. 426102 Biolegend) and stained with conjugated primary antibodies as per manufacturer’s protocol and pre-titrated antibody dilutions (supplementary table S1). Intracellular staining was performed following fixation and permeabilization (Fixation and Permeabilization Buffer Set, eBioscience). Cells were washed and resuspended in flow buffer and acquired using BD LSRII with FACSDiva Software (v 9.0). analyses were performed using FlowJo (v10) (TreeStar).

The following antibodies were used to stain myeloid and lymphoid cells:

Mouse BMDMs: Anti-CD45 (clone 30-F11), anti-CD124 (clone I015F8), anti-I-A/I-E (clone 107630), anti-Ly6C (clone HK1.4), anti-CD11b (clone M1/70), anti-F4/80 (clone BM8), anti-Ly6G (clone 1A8), anti CD206 (clone C068C2), anti-TNF (clone MP6-XT22) all Biolegend), anti-IL-10 (clone JES5-16E3 eBiosciences), and anti-Glut1 (clone EPR3915, Abcam).

Human HMDMs: anti CD16 (clone 3G8), anti-CD14 (clone 63D3), anti-HLA-DR (clone L243), anti-CD11b (clone ICRF44), anti-CD206 (clone 15-2), anti-CD163 (clone GHI/61), anti-TNF (clone MAb11), and anti-TNF (clone MAb11) all Biolegend.

Mouse macrophages (syngeneic MB49 model of urothelial carcinoma): CD45 (clone 30-F11, Biolegend), CD124 (IL-4Ra) (clone I015F8, Biolegend), I-a/I-e (clone M5/114.15.2, Biolegend), F4/80 (clone BM8, Biolegend), CD206 (clone C068C2, Biolegend), TNF (clone MP6-XT22, Thermo Fisher), IL-10 (clone JES5-16E3, Thermo Fisher).

Mouse T cells (syngeneic MB49 model of urothelial carcinoma): CD45 (clone PerCP, Biolegend), CD25 (clone PC61, Biolegend), CD3 (clone 17A2, Biolegend), CD4 (clone GK1.5, Biolegend), CD8a (clone 53-6.7, Biolegend), FOXP3 (clone MF-14, Biolegend), Mouse IFN-γ (clone XMG1.2, Biolegend) and FOXP3 (clone MF-14 Biolegend), CD69 (cloneH1.2F3, Biolgened), CD38 (clone IM7, Biolegend).

#### *In vitro* monocyte trained immunity experiment

*In vitro* training of primary human monocytes was performed according to the well-established model.^34^ Briefly, PBMCs were isolated from healthy donors (leukopaks). Following magnetic separation, CD14^+^ monocytes were seeded in 10 mm^3^ tissue culture dishes for 3 hours in warm RPMI 1640 media supplemented with 10% FBS at 37°C with 5% CO2. Non-adherent cells were removed by washing cells using warm PBS. Monolayer culture of human monocytes was infected with BCG-WT and BCG-*disA*-OE strains at 5:1 (monocyte versus BCG) MOIs for 4 hours in presence of RPMI 1640 supplemented with 10% FBS. Non-internalized bacilli were washed out using warm PBS and subsequently incubated for 24 hours. Cells were again washed using warm PBS and fresh warm RPMI 1640 media was added. For the following 5 days, cells were allowed to rest with a PBS wash and addition of fresh media every 2^nd^ day. Cells were re-stimulated on day 6 with RPMI 1640 supplemented with 10% FBS (negative control, without training) or TLR1/2 agonist, Pam3Cys (Cat. tlrl-pms, InvivoGen). Following stimulation, for 24 h, culture supernatants were collected, filter sterilized and quickly snap-frozen (-80°C) for cytokine measurement. Cells were harvested for chromatin immunoprecipitation (ChIP) experiments to measure epigenetic changes on gene promoters.

#### Chromatin immunoprecipitation (ChIP)

Human monocytes were fixed with a final concentration of 1% formaldehyde for 10 minutes at room temperature. Cell fixation was stopped using 125 mM glycine (Cat no. 50046, Sigma-Aldrich, USA), followed by sonication to fragment cellular DNA to an average size between 300 to 600 bp using Qsonica Sonicator Q125 (Cat. 15338283, Thermo Fisher Scientific). Sonicated cell lysates were subjected to immunoprecipitation (IP) by overnight incubation with recommended concentration of primary antibodies [(Histone H3K9me3 (H3K9 Trimethyl) Polyclonal Antibody cat. A-4036-100, epigentek); Anti-Histone H3 (tri methyl K4) antibody - ChIP Grade (ab8580), abcam)] in presence of magnetic Dynabeads (Cat no. 10004D, Thermo Fisher Scientific, USA) at 4°C. Non-bound material was removed by sequentially washing the Dynabeads with lysis buffer, chromatin IP (ChIP) wash buffer and Tris-EDTA (TE buffer). DNA elution was done using ChIP elution buffer. Amplification of different segments of the regulatory regions of immunity genes was carried out using qPCR using specific primers. Reactions were normalized with input DNA while beads served as negative control. Details of all primary antibodies and sequence of primers have been given in supplementary **Table. S2**.

#### Targeted Metabolite analysis with LC-MS/MS

Targeted metabolite analysis was performed with liquid-chromatography tandem mass spectrometry (LC-MS/MS)^48^. Metabolites from cells were extracted with 80% (v/v) methanol solution equilibrated at –80 °C, and the metabolite-containing supernatants were dried under nitrogen gas. Dried samples were re-suspended in 50% (v/v) acetonitrile solution and 4ml of each sample were injected and analyzed on a 5500 QTRAP triple quadrupole mass spectrometer (AB Sciex) coupled to a Prominence ultra-fast liquid chromatography (UFLC) system (Shimadzu). The instrument was operated in selected reaction monitoring (SRM) with positive and negative ion-switching mode as described. This targeted metabolomics method allows for analysis of over two hundred of metabolites from a single 25-min LC-MS acquisition with a 3-ms dwell time and these analyzed metabolites cover all major metabolic pathways. The optimized MS parameters were: ESI voltage was +5,000V in positive ion mode and –4,500V in negative ion mode; dwell time was 3ms per SRM transition and the total cycle time was 1.57 seconds. Hydrophilic interaction chromatography (HILIC) separations were performed on a Shimadzu UFLC system using an amide column (Waters XBridge BEH Amide, 2.1 x 150 mm, 2.5μm). The LC parameters were as follows: column temperature, 40 ℃; flow rate, 0.30 ml/min. Solvent A, Water with 0.1% formic acid; Solvent B, Acetonitrile with 0.1% formic acid; A non-linear gradient from 99% B to 45% B in 25 minutes with 5min of post-run time. Peak integration for each targeted metabolite in SRM transition was processed with MultiQuant software (v2.1, AB Sciex). The preprocessed data with integrated peak areas were exported from MultiQuant and re-imported into Metaboanalyst software (MetaboAnalyst (V5.0) (https://www.metaboanalyst.ca) for further data analysis including statistical and principal component analyses.

#### Glucose uptake assay

Glucose uptake measurement was performed on bone-marrow-derived macrophages (BMDMs) isolated from C57BL/6 females in an in vitro BCG infection assay. Briefly, macrophages were infected at a ratio of 1:20 (macrophage vs BCG-WT or BCG-*disA*-OE) in presence of DMEM medium devoid of glucose for 4 hours. Exogenous addition of 2-NBDG was carried out and cells were stained for cell surface markers (Glut1 and CD45) after 2 hours of incubation. Expression of Glut1 expression and 2-NBDG positivity was determined using flow-cytometry analyses using FACSDiva (v 9.0) and FlowJo (v10) (TreeStar).

#### Histologic analyses and immunohistochemistry (IHC)

For histologic analyses, a portion of bladder was formalin fixed and paraffin embedded. Sections of 5μ in thickness on glass slides were stained with hematoxylin-eosin for classification according to the World Health Organization/International Society of Urological Pathological consensus^27^. Tumor staging was performed by 2 board certified genitourinary pathologists (A.S.B., A.M.) blinded to treatment groups. Specimens were classified based on the percentage of involvement of abnormal tissue (1 = 10% involvement, 2 = 20% involvement, and so forth). For IHC staining, high-temperature antigen retrieval (18–23 psi/126 °C) was performed by immersing the slides in Trilogy (Cell Marque). Endogenous peroxidase activity was blocked for 5 min in using Dual Endogenous Enzyme Block (Cat. S2003, Dako). Primary Antibodies used included Ki67 (1:50, Cat. ab16667; Abcam), CD68 (1:250, Cat. MCA341R; Serotec), CD86 (1:100, Cat. bs-1035R; Bioss) and CD206 (1:10K, Cat. ab64693; Abcam). For Ki67, slides were stained with ImmPACT DAB (Vector Labs) for 3 min and counterstained with haematoxylin (Richard-Allen). Dual staining for CD68/CD206 and CD68/CD86 was achieved by first staining for CD68 with Impact DAB (Vector Labs) followed by secondary antigen retrieval and incubation as above with either CD86 or CD206 and visualized with ImmPACT AEC (Vector Labs). For each section, Ki67 expression was scored as a percentage of positive cells in the urothelium. Dual stains for CD68/CD86 and CD68/CD206 were scored based on positive clusters of cells for each marker (0= no staining, 1 = rare isolated cells positive, 2 = clusters of up to 10 positive cells, 3= clusters of > 10 positive cells).

### In vivo experiments

#### Intravesical BCG treatment in carcinogen induced NMIBC rat model

The induction of urothelial cancer in rats and subsequent treatment of intravesical BCG were carried out using our published protocol^21^. Briefly, N-methyl-N-nitrosourea (MNU) instillations were given every other week for a total of 4 instillations. Fischer 344 female rats age 7 weeks (Harlan, avg. weight 160g) were anesthetized with 3% isoflurane. After complete anesthesia, a 20G angiocatheter was placed into the rat’s urethra. MNU (1.5mg/kg) (Spectrum) dissolved in 0.9.% sodium chloride was then instilled and the catheter removed, with continued sedation lasting for 60 minutes to prevent spontaneous micturition and allow absorption. Eighteen weeks after the first MNU instillation, intravesical treatment with PBS or 5 x 10^6^ CFU of each BCG strain (0.3ml via a 20G angiocatheter) was administered weekly for a total of 6 doses. Animals were monitored regularly and studies were carried out in accordance with the tumor guidelines of JHU Animal Care and Use Committee. Rodents were sacrificed 2 d after the last intravesical treatment, and bladders were harvested within 48 hours of the last BCG instillation for mRNA and protein expression analysis as well as histological evaluation.

#### BCG infection of BALB/c mice and CFU enumeration

To determine the lung bacillary burden of wild-type and BCG-disA-OE strains 6-week-old female BALB/c mice were exposed using the aerosol route in a Glas-Col inhalation exposure system (Glas-Col). The inoculum implanted in the lungs at day 1 (n=3 mice per group) in female BALB/c mice was determined by plating the whole lung homogenate on 7H11 selective plates containing carbenicillin (50 mg/ml), Trimethoprim (20 mg/ml), Polymyxin B (25 mg/ml) and Cycloheximide (10 mg/ml). Following infection, mice lungs were harvested (n = 5 animals/group), homogenized in their entirety in sterile PBS and plated on 7H11 selective plates at different dilutions. The 7H11 selective plates were incubated at 37 °C and single colonies were enumerated at week 3 and 4. Single colonies were expressed at log CFU per organ.

#### SCID Mice time to death study

The virulence testing of BCG-WT and BCG-*disA*-OE strains was done in severely compromised immunodeficient mice aerosol infection model established in our laboratory. The inoculum implanted in the lungs at day 1 (n = 3 animals per group) was determined by plating the whole lung homogenate on 7H11 selective plates. For time to death analyses (n = 10 animals per group) infected animal were monitored until their death.

#### Syngeneic MB49 model of urothelial cancer

MB49 tumor cells are urothelial carcinoma line derived from an adult C57BL/6 mouse by exposure of primary bladder epithelial cell explant to 7,12-dimethylbenz[a]anthracene (DMBA) for 24 hours followed by a long-term culture^79^. Before implantation, MB49 cells were cultured as monolayers in RPMI 1640 media supplemented with 10% FBS and 1% streptomycin/penicillin at 37°C with 5% CO2. Cells were harvested using Trypsinization and cell viability was determined using Trypan blue dye. Live MB49 cells were resuspended in sterile PBS and adjusted at 1 x 10^5^ live cells per 100 μl. Female C57BL/6J mice, age 4-6 weeks (Charles River Laboratories) were subcutaneously injected with 1 x 10^5^ MB49 cells in the right flank of hind leg. Tumor growth was monitored every 2^nd^ day to observe the increase the tumor burden at the time of treatment initiation. Once palpable tumor developed (7 to 9 days, average volume ∼ 30 mm^3^ 1 x 10^6^ bacilli of BCG-WT or BCG-*disA*-OE in a total 50 μl PBS was injected intratumorally (Fig. 1h). A total of 4 intratumoral injections of BCG were given every 3^rd^ day. Tumors were measured by electronic caliper, and tumor volume was calculated using the following equation: tumor volume = length x width x height x x 0.5326. We did not allow to exceed the maximum allowed tumor volume of ∼2 cm in any dimension was based on the guidelines of our Institutional Animal Care and Use Committee for a single implanted tumor that is visible without imaging. Mice were killed at specified time, and tumors and spleens were collected after necropsy for single cell preparation.

#### Statistics and Reproducibility

MNU rat study involved a minimum sample size of 5 animals (n=5 per group each biological replicate) and were replicated for statistical significance. Treatment outcomes were determined using one-way ANOVA with Tukey’s test for multiple comparisons, one-way ANOVA with Dunnett’s test for multiple comparisons or 2-sided Fisher’s Exact test. Animal studies involving MB49 tumor studies (minimum sample size n=6 animals per group) were replicated to determine tumor volume. Two-way ANOVA with Tukey’s test for multiple comparisons was utilized to determine statistical significance between groups across time. We assessed endpoint (Mock vs Treatment groups) by one-way ANOVA with Dunnett’s test for multiple comparisons. To determine the statistical significance between BCG-WT vs BCG-*disA-* OE at endpoint, we utilized a two-tailed Student’s T-test. Immune infiltrate analyses were assessed by two-way ANOVA with Tukey’s test for multiple comparisons. Cell-based assays (cytokine quantifications, gene expression analyses and cellular phenotyping) were performed in a minimum of three (n=3) independent biological replicates to derive statistical significance (one-way ANOVA w/ Tukey’s test for multiple comparisons, two-way ANOVA w/ Tukey’s test for multiple comparisons, unpaired/paired two-tailed student’s T-test, and two-sided Fisher’s exact test). Balb/c murine experiments were performed using n=3 mice on Day 1 and n=5 mice on Day 28. Student T-tests were utilized to assess significance. SCID murine experiments were performed using n=2 mice on Day 1 (two-tailed Student’s T-test) and n=10 mice to assess survival by Kaplan-Meier analysis. Metabolite experiments were performed using n=2 and n=4 (Two-tailed student’s T-test and one-way ANOVA with Tukey’s test for multiple comparisons, respectively). All data are expressed as mean values ± S.D. The results were significant when ****P < 0.0001; ***P < 0.005; **P < 0.01; *P < 0.05 as given in the figure legends. Description of exact number of biological replicates, statistical tests employed, and P values are given in detail. GraphPad Prism (v 10.0.3) was used for analyses.

MNU rats were randomly assigned to different groups and were blinded for treatment arms. The histopathological assessment (IHCs, tumor staging and tumor involvement index) were performed by a genitourinary pathologist blinded for treatment groups. Periodic contamination testing of mammalian cells and BCG strains ensured absence of cross-contamination. None of the data was excluded from the analyses unless the recording quality was poor (e.g., absence of sufficient viable single cells, etc.) or animals developed ulcerate tumors or were moribund. All attempts at replication were successful for *in vivo*, *ex-vivo* and *in-vitro* assays. The equipment parameters, antibody dilutions, cell numbers and experimental conditions are given in the methods to ensure reproducibility.

## SUPPLEMENTARY FIGURE LEGENDS

**Supplementary Figure S1. Validation of *disA* overexpression in BCG-*disA*-OE and induction of IRF3 signaling. a.** mRNA levels of *disA* in log-phase BCG cultures relative to *M. tuberculosis sigA* (Rv2703) (n=3 independent biological replicates). **b**. IRF3 induction measured in RAW-Lucia ISG reporter macrophages. IRF3 induction was quantified using culture supernatants of macrophages infected at an MOI of 20:1 for 24 hrs (n=4 independent biological replicates). Data reflect means values ± SD. Statistical analyses done using 2-tailed student’s T-test in panel **a**; one-way ANOVA w/Tukey’s test for multiple comparisons in panel **b** (**** p < 0.0001).

Supplementary Figure S2. BCG-*disA*-OE causes reduced tumor growth and greater tumor-associated necrosis in the heterotopic syngeneic MB49 mouse model of urothelial cancer**. a.** Tumors at necropsy on day 21 (n=9 animals/group). **b.** Representative H & E staining showing necrotic area and congestion in MB49 tumors. Similar observations were made in randomly selected 3 (n=3) tumor tissue slides per group. Untreated group shows densely packed tumor cells; BCG-WT (Tice) tumor cells with moderate necrosis (below dashed line), BCG-*disA*-OE (Tice) with extensive necrosis (below dashed line) and congestion (*). (Related to **Fig. 3a-b**).

**Supplementary Figure S3**. Improved antitumor efficacy of BCG-*disA*-OE is associated with differential recruitment of T cells and macrophages to tumors and is STING-dependent in the MB49 model a. Schematic diagram of the MB49 syngeneic mouse model of urothelial tumors used in this experiment. **b.** Total CD3^+^ T cells of all CD45+ leucocytes in tumors. **c.** IFNγ^+^ tumor-infiltrating CD8^+^ T cells. **d.** activated CD8^+^ T cells (percent CD69+ CD38+ of CD8+). **e.** TNF^+^-expressing immunosuppressive macrophages (percent TNFα+ of CD206+ CD124+ F4/80+CD11b+) in MB49 tumors after necropsy. Data are presented as mean values ± S.D. (n=6 animals/group). Statistical analyses done using two-way ANOVA with Tukey’s test for multiple comparisons. (* p < 0.05, ** p < 0.01, ***p < 0.001, **** p < 0.0001).

**Supplementary Figure S4. BCG-*disA*-OE (Pasteur) is less pathogenic that BCG-WT in two mouse models. a.** Using the same experimental scheme shown in Fig. 7a, BALB/c mice were aerosol infected and lung colony forming unit (CFU) counts at day 1 are shown (n=3 animals/group). **b.** Lung CFU counts for BALB/c mice at day 28 (n=5 animals/group). **c.** Using the same experimental scheme shown in Fig. 7c, SCID mice were aerosol infected and lung colony forming unit (CFU) counts at day 1 (n = 2 animals/group). **d**. Survival of SCID mice following low dose challenge (n=10 animals/group). The experiment was performed with BCG strains in the Pasteur background. Similar results were obtained with strains in the Tice background as shown in **Fig. 4**. Data are presented as mean values ± S.D. Statistical analyses done using 2-tailed Student’s t-test (** p < 0.01).

Supplementary Figure S5. BCG-*disA*-OE elicits stronger IFN-β responses than BCG-WT in murine bone marrow-derived macrophages (BMDM). a. IFN-β levels in resting and IFN-γ primed BMDMs (n=3 biological replicates). IFN-β levels were measured by RT-qPCR after a 6 hr exposure at a MOI of 20:1. Data are presented as mean values ± S.D. Gene expression analyses for cytokines and chemokines were performed 6 hr post-exposure. Statistical analyses performed using two-way (**Fig. S5a**) and one-way (**Fig. S2b-c**) ANOVA w/Tukey’s multiple comparisons test in panel **a** (* p < 0.05, ** p < 0.01, *** p < 0.001, **** p < 0.0001).

**Supplementary Figure S6. Representative schematic of gating strategy to identify various myeloid populations in murine BMDMs.** a. Schematic of generation of BMDMs. b. Representative gating scheme for identification of different myeloid cells. Briefly, leukocyte lineage was selected by gating SSC-A against CD45^+^ populations on live cells. CD11b^+^F4/80^+^ macrophages were identified out of CD45^+^ population. CD11b^+^F4/80^+^ macrophages were divided into MHC class II (I-a/I-e) and CD124+CD206+ populations. Expression of TNFα (M1-like macrophages) and IL-10 (M2-like macrophages) were determined on MHC class II subsets and CD124^+^CD206^+^ subsets respectively. (Related to **Fig. 6a-e**).

Supplementary Figure S7. BCG-*disA*-OE induces macrophage reprogramming and favors a stronger inflammatory macrophage shift in murine BMDMs. a. Representative FACS plots for TNF-a^+^ M1-like macrophages (MHC Class II^+^CD11b^+^F4/80^+)^ corresponding to **Fig. 6a**. **b.** Representative FACS plots for M2-like macrophages (CD206^+^CD124^+^) corresponding to **Fig. 6b**. representative FACS plots. **c.** Representative FACS plots for IL-10^+^ M2-like macrophages (CD206^+^CD124^+^) corresponding to **Fig. 6c**.

**Supplementary Figure S8. Gating scheme showing identification of myeloid-derived suppressor cell populations in primary mouse macrophages after BCG exposure.** Leukocyte lineage was determined on live cells by gating SSC-A against CD45+ myeloid cells. Myeloid cells were differentiated into CD11b^+^F4/80^+^ macrophages out of which CD11b^+^F4/80^-^ myeloid population was divided into Ly6C and Ly6G. Next, the Ly6C^(hi)^Ly6G^-^ immunosuppressive myeloid-derived suppressor cell populations were looked for IL-10 positivity (Related to **Fig. 6d-e**).

**Supplementary Figure S9. Immunosuppressive monocytic-MDSCs (M-MDSCs) populations murine primary macrophages after BCG exposure. a.** Representative FACS plots for M-MDSC measurements corresponding to **Fig. 6d**. **b.** Representative FACS plots for IL-10^+^ expressing M-MDSCs corresponding to **Fig. 6e**.

**Supplementary Figure S10. The STING agonist c-di-AMP causes induction of macrophage activation.** Human macrophages were transfected with c-di-AMP for 24 h and phagocytosis of FITC-labeled IgG opsonized latex beads (green) was visualized using confocal microscopy on live cells. Hoechst was used for nuclear staining (blue). Images were acquired using LSM700 confocal microscope at 63X magnification. Images were process using Fiji software. Similar results were observed across two (n=2) independent biological replicate experiments.

Supplementary Figure S11. BCG-*disA*-OE elicits greater autophagy induction than BCG-WT in 5637 human urothelial carcinoma cells. Autophagy induction in the 5637 human urothelial carcinoma cells in representative confocal photomicrographs. Co-localization of FITC-labeled BCG strains (green), LC3B autophagic puncta (red) appears in yellow; nuclei are blue. Quantification of co-localized BCG and LC3b puncta is shown at right. Cells were fixed using 4% paraformaldehyde 3h after infection (MOI 10:1), and images obtained with an LSM700 confocal microscope and Fiji software processing. Statistical analyses done using 2-tailed Student’s t-test (** p < 0.01). Data shown are for BCG strains in the Tice background.

**Supplementary Figure S12. BCG induced differential glucose uptake in bone-marrow-derived macrophages (BMDMs). (a)** Experimental layout showing the strategy employed to determine intracellular uptake of fluorescent glucose. Briefly macrophages were infected at an MOI of 20:1 (BCG to macrophage ratio) in the presence of glucose-free medium followed by exogenous addition of 2-(*N*-(7-Nitrobenz-2-oxa-1,3-diazol-4-yl)Amino)-2-Deoxyglucose (2-NBDG). Macrophages were subsequently stained for GLUT1 and were investigated using flow cytometry. **(b-c)** Bar diagram showing induced expression of GLUT1 and intracellular fluorescent 2-NBDG in BMDMs following infection by BCG strains. Data are presented as mean values ± S.D. (n=2 independent biological replicate experiments). Data analyses were carried out using FACSDiva (v 9.0), Flowjo (v 10) and Graphpad Prism software (v 10.0.3). Statistical analysis employed a one-way ANOVA with Tukey’s test for multiple comparisons (* p < 0.05, ** p < 0.01, ***p < 0.001, **** p < 0.0001).

## REFERENCES

1. Saginala, K., et al. Epidemiology of Bladder Cancer. Med Sci (Basel) 8 (2020).

2. Grabe-Heyne, K., et al. Intermediate and high-risk non-muscle-invasive bladder cancer: an overview of epidemiology, burden, and unmet needs. Front Oncol 13, 1170124 (2023).

3. Morales, A., Eidinger, D. & Bruce, A.W. Intracavitary Bacillus Calmette-Guerin in the treatment of superficial bladder tumors. J Urol 116, 180–183 (1976).

4. Fankhauser, C.D., Teoh, J.Y. & Mostafid, H. Treatment options and results of adjuvant treatment in nonmuscle-invasive bladder cancer (NMIBC) during the Bacillus Calmette-Guérin shortage. Curr Opin Urol 30, 365–369 (2020).

5. Roumiguié, M., et al. International Bladder Cancer Group Consensus Statement on Clinical Trial Design for Patients with Bacillus Calmette-Guérin-exposed High-risk Non-muscle-invasive Bladder Cancer. Eur Urol 82, 34–46 (2022).

6. Tan, W.S., et al. Intermediate-risk Non-muscle-invasive Bladder Cancer: Updated Consensus Definition and Management Recommendations from the International Bladder Cancer Group. Eur Urol Oncol 5, 505–516 (2022).

7. Pettenati, C. & Ingersoll, M.A. Mechanisms of BCG immunotherapy and its outlook for bladder cancer. Nat Rev Urol 15, 615–625 (2018).

8. Lobo, N., et al. 100 years of Bacillus Calmette-Guérin immunotherapy: from cattle to COVID-19. Nat Rev Urol 18, 611–622 (2021).

9. Bowyer, L., Hall, R.R., Reading, J. & Marsh, M.M. The persistence of bacille Calmette-Guérin in the bladder after intravesical treatment for bladder cancer. Br J Urol 75, 188–192 (1995).

10. Durek, C., et al. The fate of bacillus Calmette-Guerin after intravesical instillation. J Urol 165, 1765–1768 (2001).

11. van Puffelen, J.H., et al. Trained immunity as a molecular mechanism for BCG immunotherapy in bladder cancer. Nat Rev Urol 17, 513–525 (2020).

12. Li, X.D., et al. Pivotal roles of cGAS-cGAMP signaling in antiviral defense and immune adjuvant effects. Science 341, 1390–1394 (2013).

13. Ablasser, A., et al. cGAS produces a 2’-5’-linked cyclic dinucleotide second messenger that activates STING. Nature 498, 380–384 (2013).

14. Woodward, J.J., Iavarone, A.T. & Portnoy, D.A. c-di-AMP secreted by intracellular Listeria monocytogenes activates a host type I interferon response. Science 328, 1703–1705 (2010).

15. Dey, B., et al. A bacterial cyclic dinucleotide activates the cytosolic surveillance pathway and mediates innate resistance to tuberculosis. Nat Med 21, 401–406 (2015).

16. Ahn, J. & Barber, G.N. STING signaling and host defense against microbial infection. Exp Mol Med 51, 1–10 (2019).

17. Dey, R.J., Dey, B., Singh, A.K., Praharaj, M. & Bishai, W. Bacillus Calmette-Guerin Overexpressing an Endogenous Stimulator of Interferon Genes Agonist Provides Enhanced Protection Against Pulmonary Tuberculosis. J Infect Dis 221, 1048–1056 (2020).

18. Singh, A.K., et al. Re-engineered BCG overexpressing cyclic di-AMP augments trained immunity and exhibits improved efficacy against bladder cancer. Nat Commun 13, 878 (2022).

19. Kates, M., et al. Preclinical Evaluation of Intravesical Cisplatin Nanoparticles for Non-Muscle-Invasive Bladder Cancer. Clin Cancer Res 23, 6592–6601 (2017).

20. Yoshida, T., et al. Ex vivo culture of tumor cells from N-methyl-N-nitrosourea-induced bladder cancer in rats: Development of organoids and an immortalized cell line. Urol Oncol 36, 160.e123–160.e132 (2018).

21. Kates, M., et al. Intravesical BCG Induces CD4(+) T-Cell Expansion in an Immune Competent Model of Bladder Cancer. Cancer Immunol Res 5, 594–603 (2017).

22. Kates, M., et al. Adaptive Immune Resistance to Intravesical BCG in Non-Muscle Invasive Bladder Cancer: Implications for Prospective BCG-Unresponsive Trials. Clin Cancer Res 26, 882–891 (2020).

23. Lérias, J.R., et al. Trained Immunity for Personalized Cancer Immunotherapy: Current Knowledge and Future Opportunities. Front Microbiol 10, 2924 (2019).

24. Gabrilovich, D.I. Myeloid-Derived Suppressor Cells. Cancer Immunol Res 5, 3–8 (2017).

25. Kumar, V., Patel, S., Tcyganov, E. & Gabrilovich, D.I. The Nature of Myeloid-Derived Suppressor Cells in the Tumor Microenvironment. Trends Immunol 37, 208–220 (2016).

26. Dahal, L.N., et al. STING Activation Reverses Lymphoma-Mediated Resistance to Antibody Immunotherapy. Cancer Res 77, 3619–3631 (2017).

27. Corrales, L., McWhirter, S.M., Dubensky, T.W., Jr. & Gajewski, T.F. The host STING pathway at the interface of cancer and immunity. J Clin Invest 126, 2404–2411 (2016).

28. Zhu, Y., et al. STING: a master regulator in the cancer-immunity cycle. Mol Cancer 18, 152 (2019).

29. Ohkuri, T., et al. Intratumoral administration of cGAMP transiently accumulates potent macrophages for anti-tumor immunity at a mouse tumor site. Cancer Immunol Immunother 66, 705–716 (2017).

30. Liu, D., et al. STING directly activates autophagy to tune the innate immune response. Cell Death Differ 26, 1735–1749 (2019).

31. Watson, R.O., et al. The Cytosolic Sensor cGAS Detects Mycobacterium tuberculosis DNA to Induce Type I Interferons and Activate Autophagy. Cell Host Microbe 17, 811–819 (2015).

32. Jagannath, C., et al. Autophagy enhances the efficacy of BCG vaccine by increasing peptide presentation in mouse dendritic cells. Nat Med 15, 267–276 (2009).

33. Crotzer, V.L. & Blum, J.S. Autophagy and its role in MHC-mediated antigen presentation. J Immunol 182, 3335–3341 (2009).

34. Arts, R.J.W., et al. BCG Vaccination Protects against Experimental Viral Infection in Humans through the Induction of Cytokines Associated with Trained Immunity. Cell Host Microbe 23, 89–100 e105 (2018).

35. Kaufmann, E., et al. BCG Educates Hematopoietic Stem Cells to Generate Protective Innate Immunity against Tuberculosis. Cell 172, 176–190 e119 (2018).

36. Kleinnijenhuis, J., et al. Bacille Calmette-Guerin induces NOD2-dependent nonspecific protection from reinfection via epigenetic reprogramming of monocytes. Proc Natl Acad Sci U S A 109, 17537–17542 (2012).

37. Arts, R.J., et al. Glutaminolysis and Fumarate Accumulation Integrate Immunometabolic and Epigenetic Programs in Trained Immunity. Cell Metab 24, 807–819 (2016).

38. Bekkering, S., et al. In Vitro Experimental Model of Trained Innate Immunity in Human Primary Monocytes. Clin Vaccine Immunol 23, 926–933 (2016).

39. Riksen, N.P. & Netea, M.G. Immunometabolic control of trained immunity. Mol Aspects Med, 100897 (2020).

40. Cheng, S.C., et al. mTOR-and HIF-1alpha-mediated aerobic glycolysis as metabolic basis for trained immunity. Science 345, 1250684 (2014).

41. Arts, R.J.W., et al. Immunometabolic Pathways in BCG-Induced Trained Immunity. Cell Rep 17, 2562–2571 (2016).

42. Gomes, M.T.R., et al. STING regulates metabolic reprogramming in macrophages via HIF-1α during Brucella infection. PLoS Pathog 17, e1009597 (2021).

43. Mbongue, J.C., et al. The Role of Indoleamine 2, 3-Dioxygenase in Immune Suppression and Autoimmunity. Vaccines (Basel) 3, 703–729 (2015).

44. Gautam, U.S., et al. In vivo inhibition of tryptophan catabolism reorganizes the tuberculoma and augments immune-mediated control of Mycobacterium tuberculosis. Proc Natl Acad Sci U S A 115, E62–e71 (2018).

45. Martini, A., Tholomier, C., Mokkapati, S. & Dinney, C.P.N. Interferon gene therapy with nadofaragene firadenovec for bladder cancer: from bench to approval. Front Immunol 14, 1260498 (2023).

46. Packiam, V.T., et al. An open label, single-arm, phase II multicenter study of the safety and efficacy of CG0070 oncolytic vector regimen in patients with BCG-unresponsive non-muscle-invasive bladder cancer: Interim results. Urol Oncol 36, 440–447 (2018).

47. Rentsch, C.A., et al. A Phase 1/2 Single-arm Clinical Trial of Recombinant Bacillus Calmette-Guérin (BCG) VPM1002BC Immunotherapy in Non-muscle-invasive Bladder Cancer Recurrence After Conventional BCG Therapy: SAKK 06/14. Eur Urol Oncol 5, 195–202 (2022).

48. Grode, L., et al. Increased vaccine efficacy against tuberculosis of recombinant Mycobacterium bovis bacille Calmette-Guérin mutants that secrete listeriolysin. J Clin Invest 115, 2472–2479 (2005).

49. Nieuwenhuizen, N.E., et al. The Recombinant Bacille Calmette-Guérin Vaccine VPM1002: Ready for Clinical Efficacy Testing. Front Immunol 8, 1147 (2017).

50. Singh, A.K., Srikrishna, G., Bivalacqua, T.J. & Bishai, W.R. Recombinant BCGs for tuberculosis and bladder cancer. Vaccine 39, 7321–7331 (2021).

51. Pieters, J. Entry and survival of pathogenic mycobacteria in macrophages. Microbes Infect 3, 249–255 (2001).

52. Vaugelade, J., Pinchinat, S., Guiella, G., Elguero, E. & Simondon, F. Non-specific effects of vaccination on child survival: prospective cohort study in Burkina Faso. Bmj 329, 1309 (2004).

53. Giamarellos-Bourboulis, E.J., et al. Activate: Randomized Clinical Trial of BCG Vaccination against Infection in the Elderly. Cell 183, 315–323.e319 (2020).

54. Singh, A.K., Netea, M.G. & Bishai, W.R. BCG turns 100: its nontraditional uses against viruses, cancer, and immunologic diseases. J Clin Invest 131 (2021).

55. Saeed, S., et al. Epigenetic programming of monocyte-to-macrophage differentiation and trained innate immunity. Science 345, 1251086 (2014).

56. Netea, M.G., et al. Defining trained immunity and its role in health and disease. Nat Rev Immunol 20, 375–388 (2020).

