## Supplemental Figures and Tables. Source Data for "Improved bladder cancer antitumor efficacy with a recombinant BCG that releases a STING agonist"

### Supplementary Figure S1

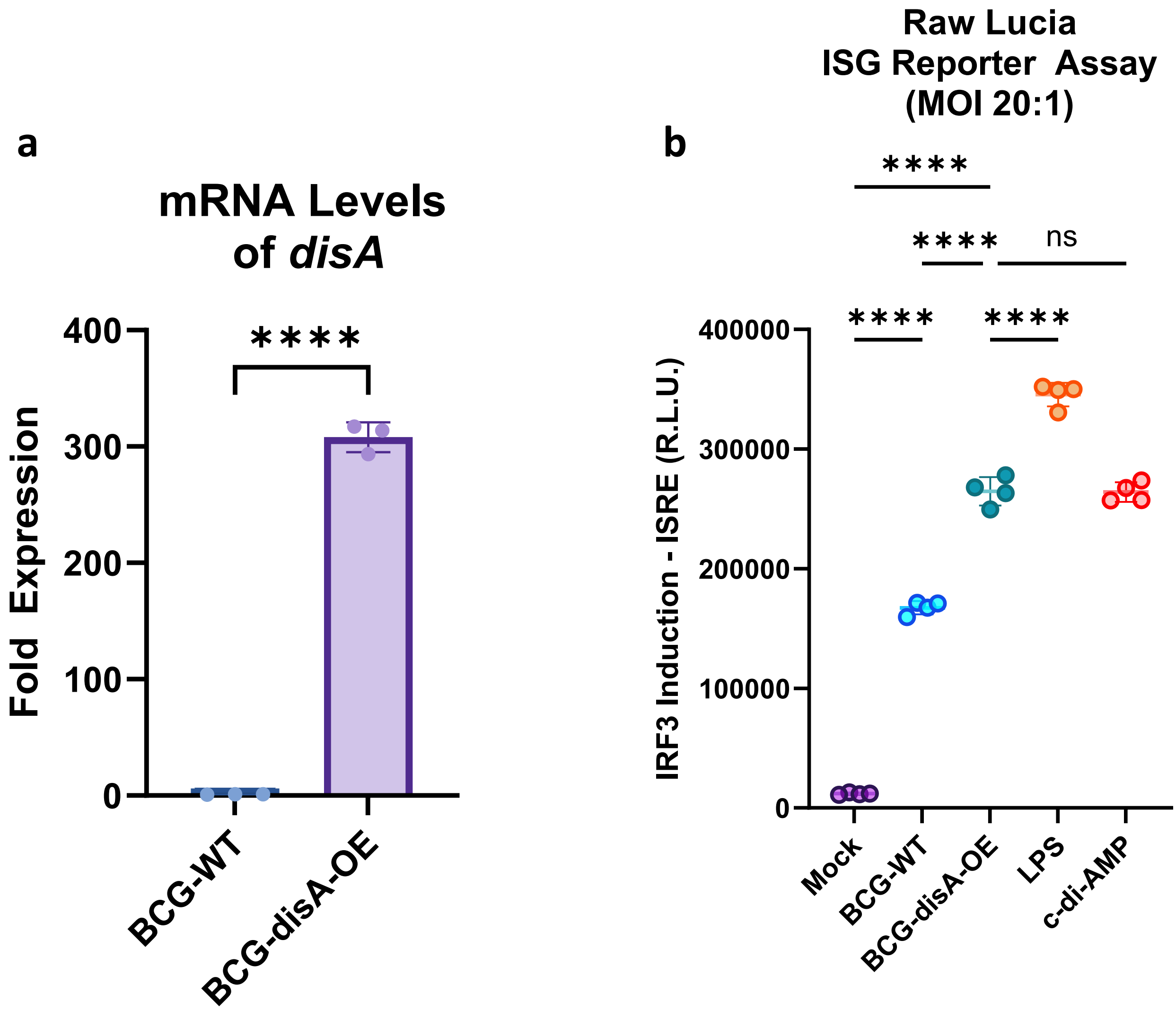

**Supplementary Figure S1. Validation of *disA* overexpression in BCG-*disA*-OE and induction of IRF3 signaling. a.** mRNA levels of *disA* in log-phase BCG cultures relative to *M. tuberculosis sigA* (Rv2703) (n=3 independent biological replicates). **b**. IRF3 induction measured in RAW-Lucia ISG reporter macrophages. IRF3 induction was quantified using culture supernatants of macrophages infected at an MOI of 20:1 for 24 hrs (n=4 independent biological replicates). Data reflect means values ± SD. Statistical analyses done using 2-tailed student's T-test in panel **a**; one-way ANOVA w/Tukey's test for multiple comparisons in panel **b** (**** p < 0.0001).

**Supplementary Figure S2**

**a**

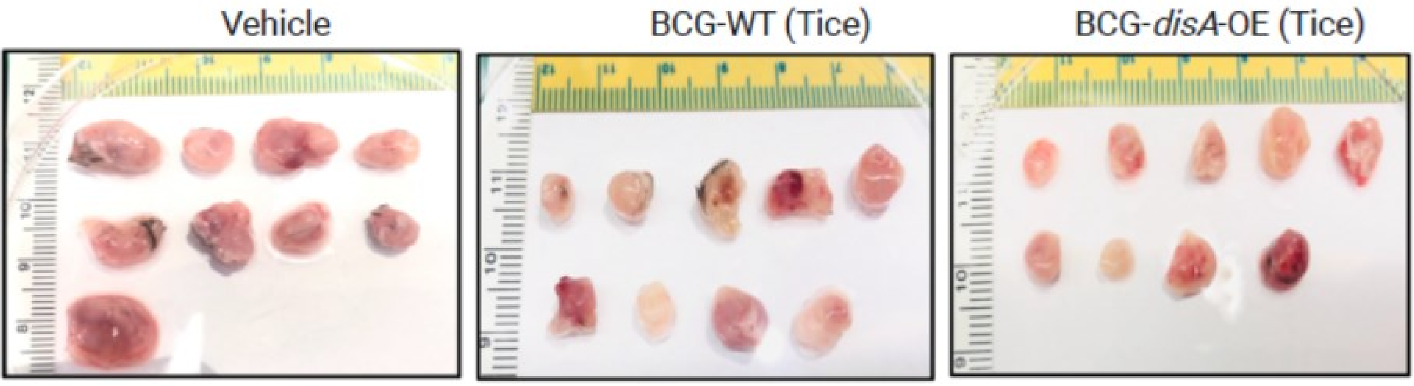

**b**

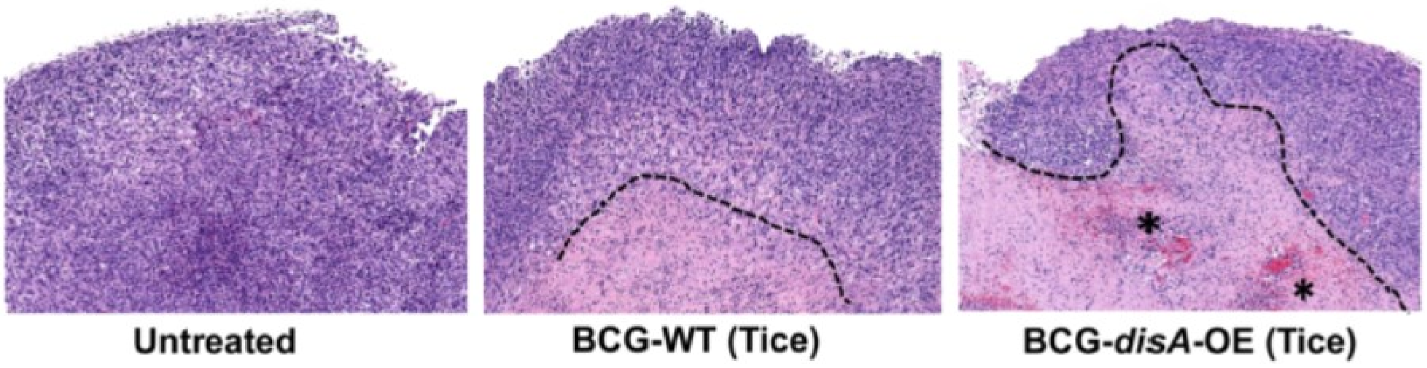

**Supplementary Figure S2. BCG-*disA*-OE causes reduced tumor growth and greater tumor-associated necrosis in the heterotopic syngeneic MB49 mouse model of urothelial cancer**. **a.** Tumors at necropsy on day 21 (n=9 animals/group). **b.** Representative H & E staining showing necrotic area and congestion in MB49 tumors. Similar observations were made in randomly selected 3 (n=3) tumor tissue slides per group. Untreated group shows densely packed tumor cells; BCG-WT (Tice) tumor cells with moderate necrosis (below dashed line), BCG-*disA*-OE (Tice) with extensive necrosis (below dashed line) and congestion (*). (Related to **Fig. 3a-b**).

### Supplementary Figure S3

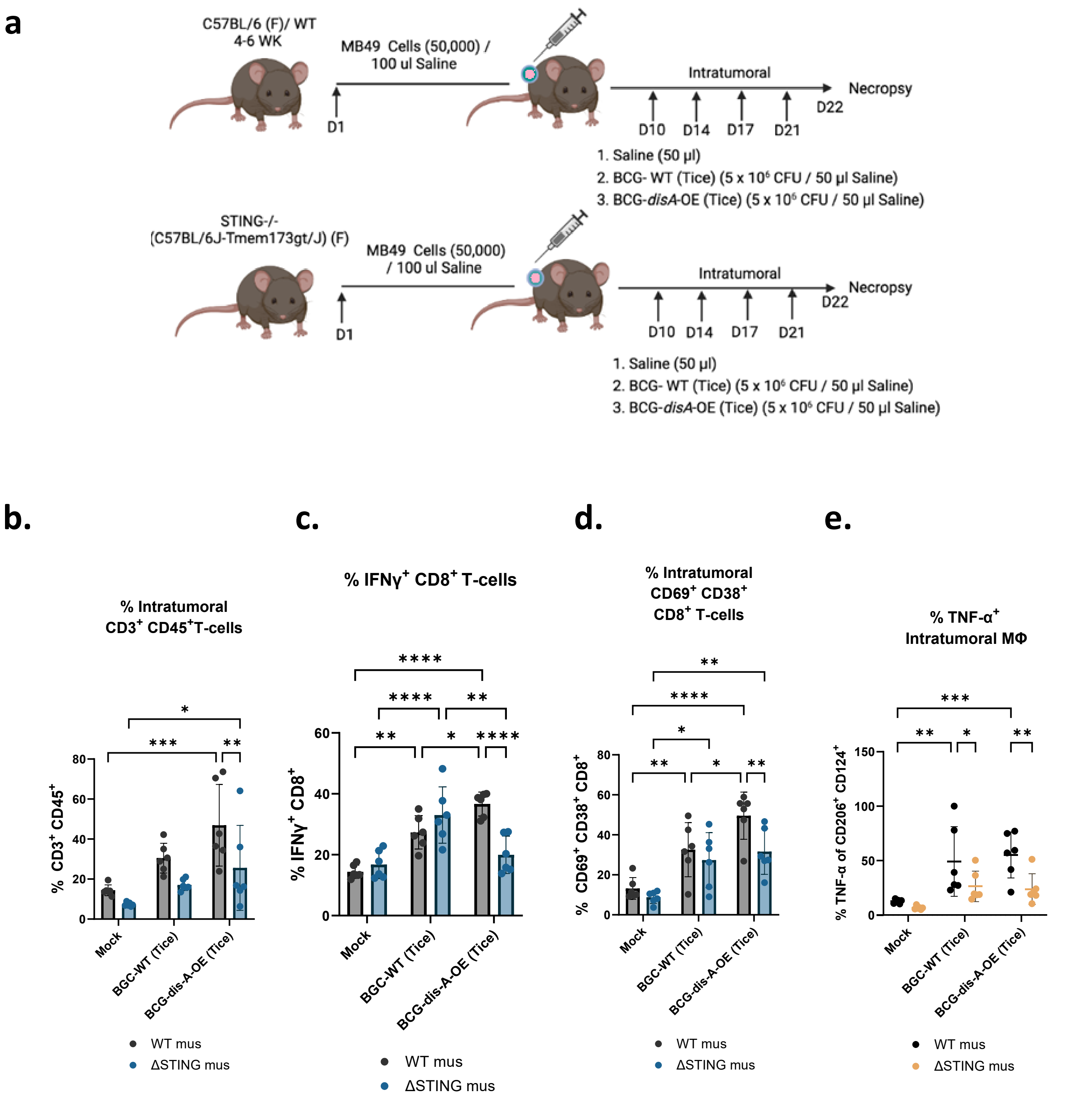

**Supplementary Figure S3. Improved antitumor efficacy of BCG-*disA*-OE is associated with differential recruitment of T cells and macrophages to tumors and is STING-dependent in the MB49 model** **a.** Schematic diagram of the MB49 syngeneic mouse model of urothelial tumors used in this experiment. **b.** Total $CD3^+$ T cells of all CD45+ leucocytes in tumors. **c.** $IFN\gamma^+$ tumor-infiltrating $CD8^+$ T cells. **d.** activated $CD8^+$ T cells (percent CD69+ CD38+ of CD8+). **e.** $TNF^+$-expressing immunosuppressive macrophages (percent TNFα+ of CD206+ CD124+ F4/80+ CD11b+) in MB49 tumors after necropsy. Data are presented as mean values ± S.D. (n=6 animals/group). Statistical analyses done using two-way ANOVA with Tukey's test for multiple comparisons. (* $p < 0.05$, ** $p < 0.01$, ***$p < 0.001$, **** $p < 0.0001$).

### Supplementary Figure S4

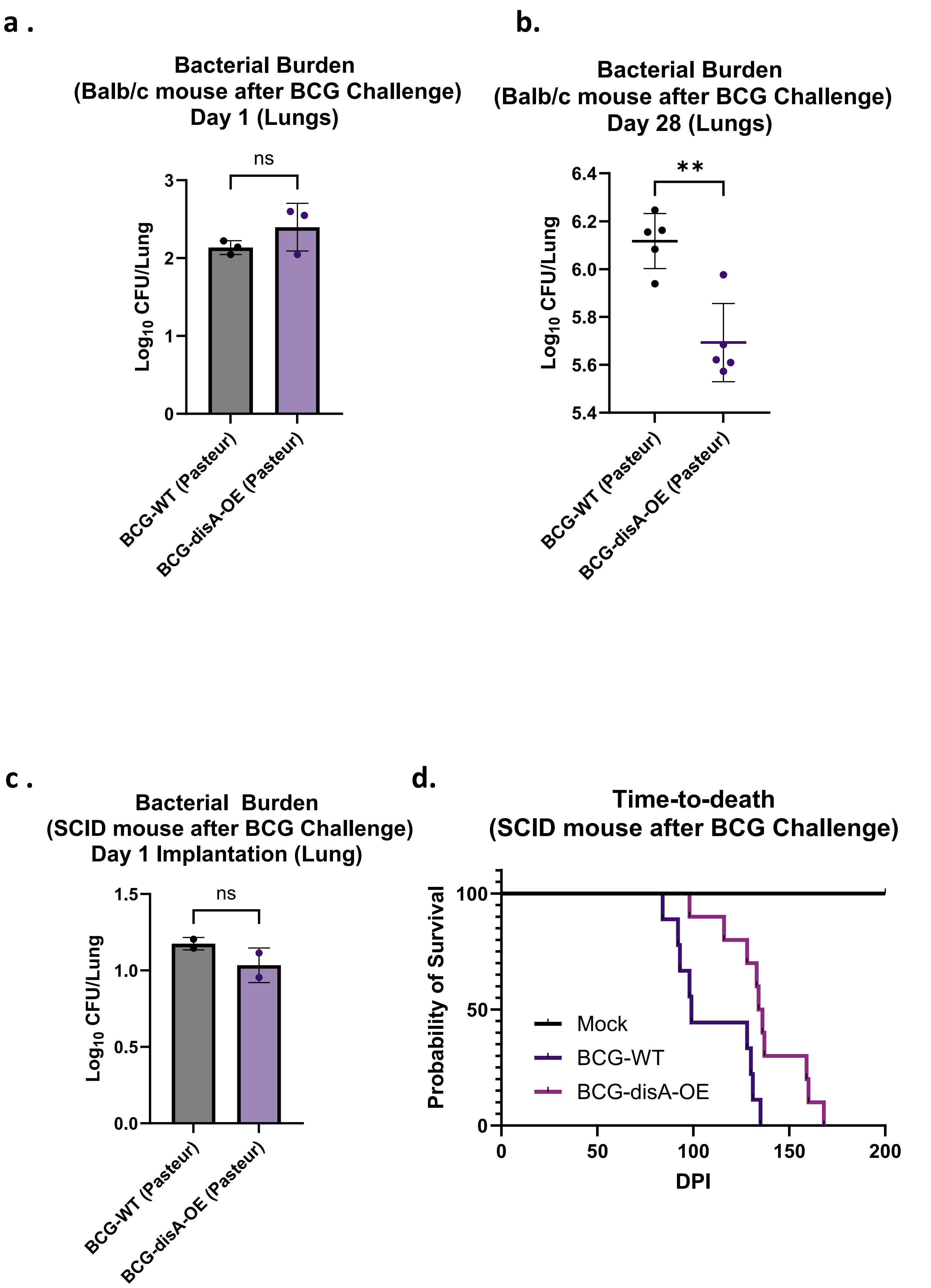

**Supplementary Figure S4. BCG-*disA*-OE (Pasteur) is less pathogenic that BCG-WT in two mouse models. a.** Using the same experimental scheme shown in Fig. 7a, BALB/c mice were aerosol infected and lung colony forming unit (CFU) counts at day 1 are shown (n=3 animals/group). **b.** Lung CFU counts for BALB/c mice at day 28 (n=5 animals/group). **c.** Using the same experimental scheme shown in Fig. 7c, SCID mice were aerosol infected and lung colony forming unit (CFU) counts at day 1 (n = 2 animals/group). **d.** Survival of SCID mice following low dose challenge (n=10 animals/group). The experiment was performed with BCG strains in the Pasteur background. Similar results were obtained with strains in the Tice background as shown in **Fig. 4**. Data are presented as mean values ± S.D. Statistical analyses done using 2-tailed Student's t-test (** $p < 0.01$).

### Supplementary Figure S5

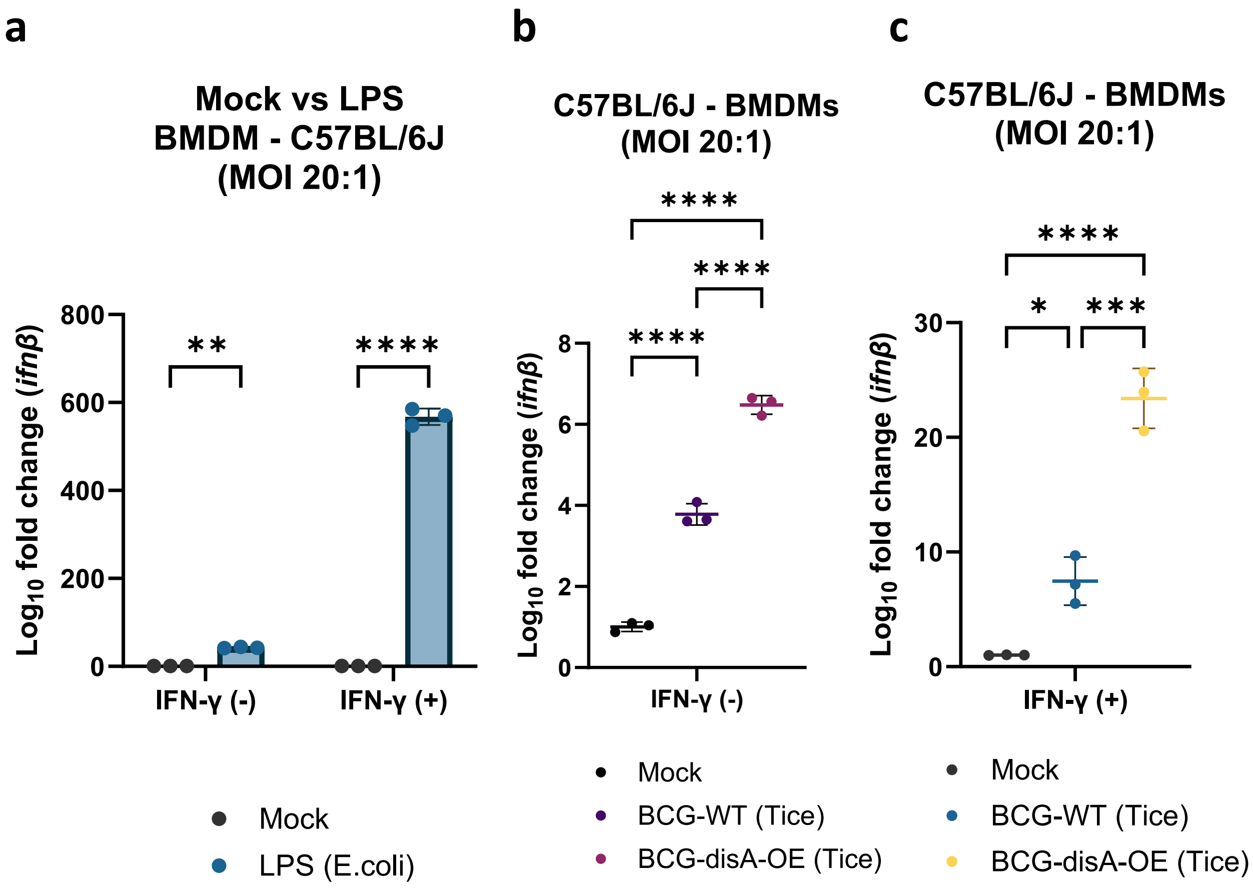

**Supplementary Figure S5. BCG-*disA*-OE elicits stronger IFN-β responses than BCG-WT in murine bone marrow-derived macrophages (BMDM). a.** IFN-β levels in resting and IFN-γ primed BMDMs (n=3 biological replicates). IFN-β levels were measured by RT-qPCR after a 6 hr exposure at a MOI of 20:1. Data are presented as mean values ± S.D. Gene expression analyses for cytokines and chemokines were performed 6 hr post-exposure. Statistical analyses performed using two-way (**Fig. S5a**) and one-way (**Fig. S2b-c**) ANOVA w/Tukey's multiple comparisons test in panel **a** (* $p < 0.05$, ** $p < 0.01$, ***$p < 0.001$, **** $p < 0.0001$).

### Supplementary Figure S6

**a**

**Immune responses in generated murine BMDMs**

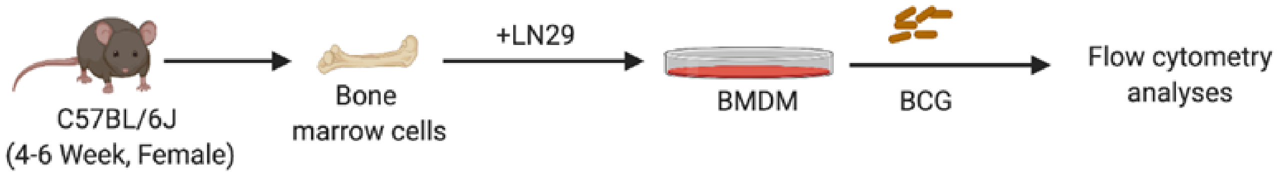

**b**

**Representative schematic of gating strategy for identifying various myeloid populations in murine BMDM**

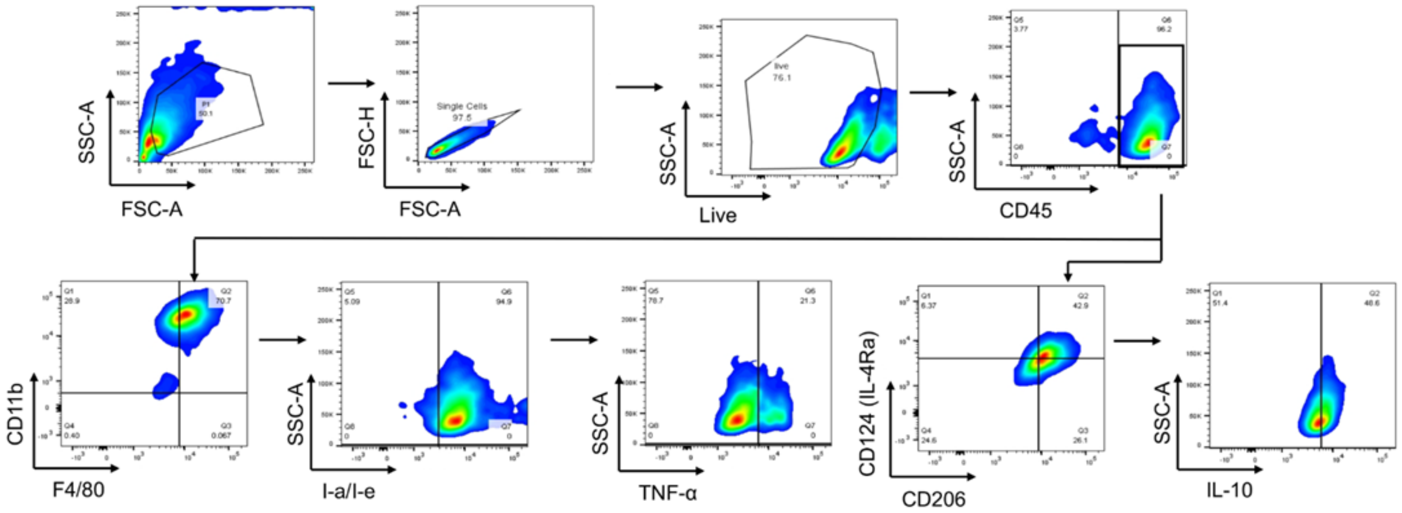

**Supplementary Figure S6. Representative schematic of gating strategy to identify various myeloid populations in murine BMDMs. a.** Schematic of generation of BMDMs. **b.** Representative gating scheme for identification of different myeloid cells. Briefly, leukocyte lineage was selected by gating SSC-A against CD45$^{+}$ populations on live cells. CD11b$^{+}$F4/80$^{+}$ macrophages were identified out of CD45$^{+}$ population. CD11b$^{+}$F4/80$^{+}$ macrophages were divided into MHC class II (I-a/I-e) and CD124+CD206+ populations. Expression of TNF$\alpha$ (M1-like macrophages) and IL-10 (M2-like macrophages) were determined on MHC class II subsets and CD124$^{+}$CD206$^{+}$ subsets respectively. (Related to Fig. **6a-e**).

### Supplementary Figure S7

**a**

**BMDM (MOI 5:1)**

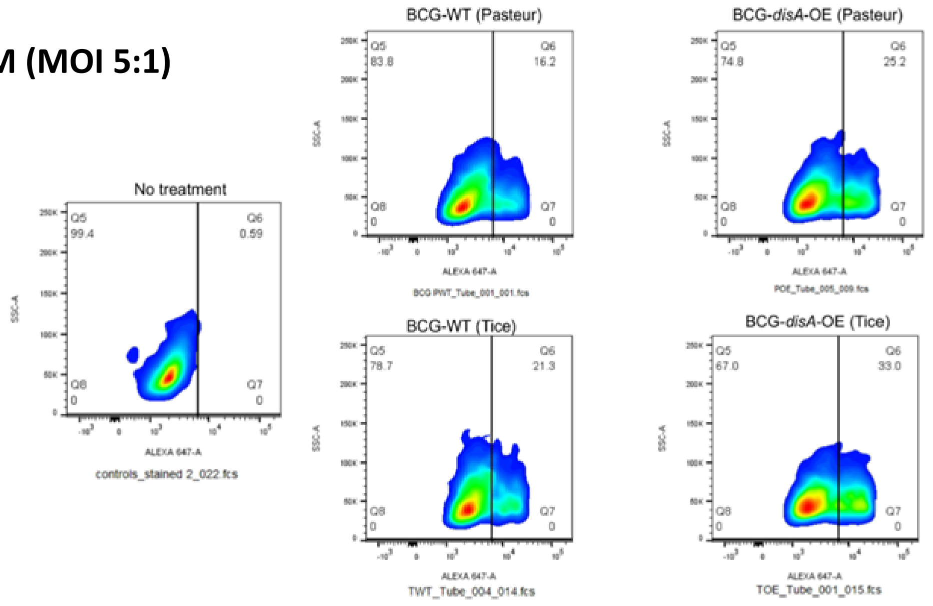

**b**

**BMDM (MOI 5:1)**

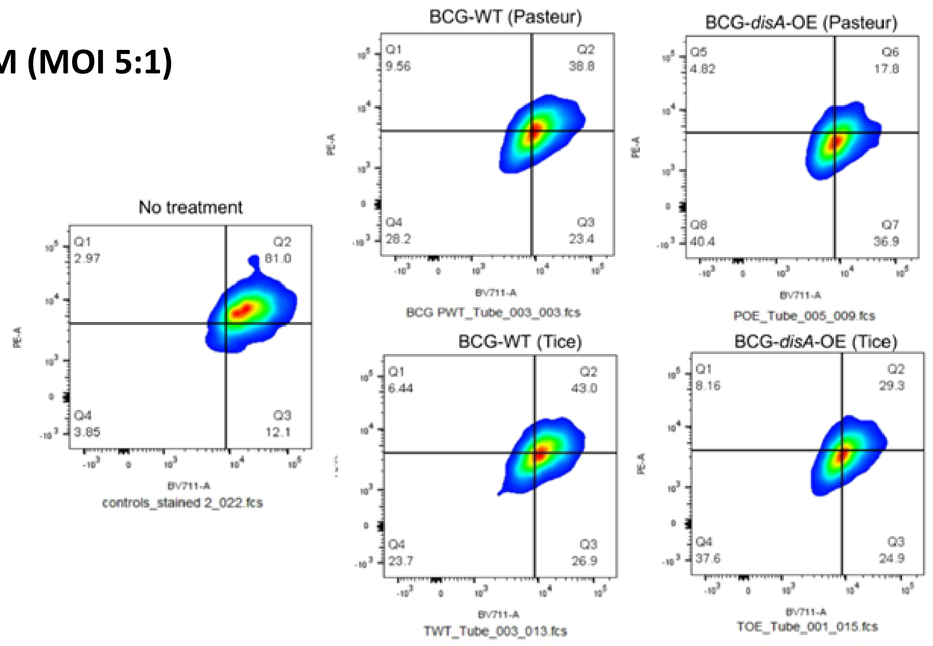

**c**

**BMDM (MOI 5:1)**

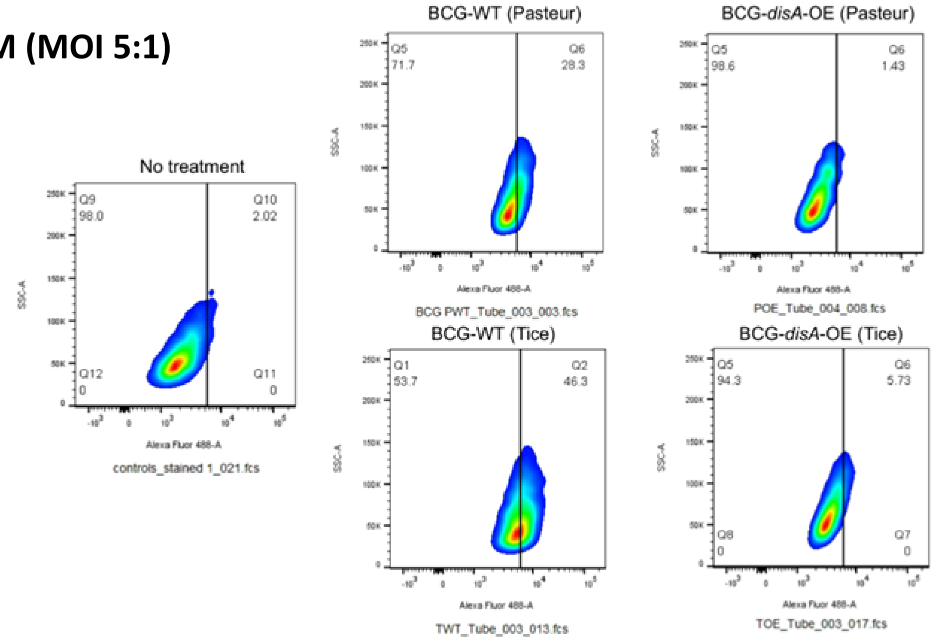

**Supplementary Figure S7. BCG-*disA*-OE induces macrophage reprogramming and favors a stronger inflammatory macrophage shift in murine BMDMs. a.** Representative FACS plots for TNF-a$^+$ M1-like macrophages (MHC Class II$^+$CD11b$^+$F4/80$^{+)}$ corresponding to **Fig. 6a**. **b.** Representative FACS plots for M2-like macrophages (CD206$^+$CD124$^+$) corresponding to **Fig. 6b**. representative FACS plots. **c.** Representative FACS plots for IL-10$^+$ M2-like macrophages (CD206$^+$CD124$^+$) corresponding to **Fig. 6c**.

### Supplementary Figure S8

**a**

**Representative gating strategy for identification of Myeloid-Derived Suppressor Cell BMDMs (C57BL/6J)**

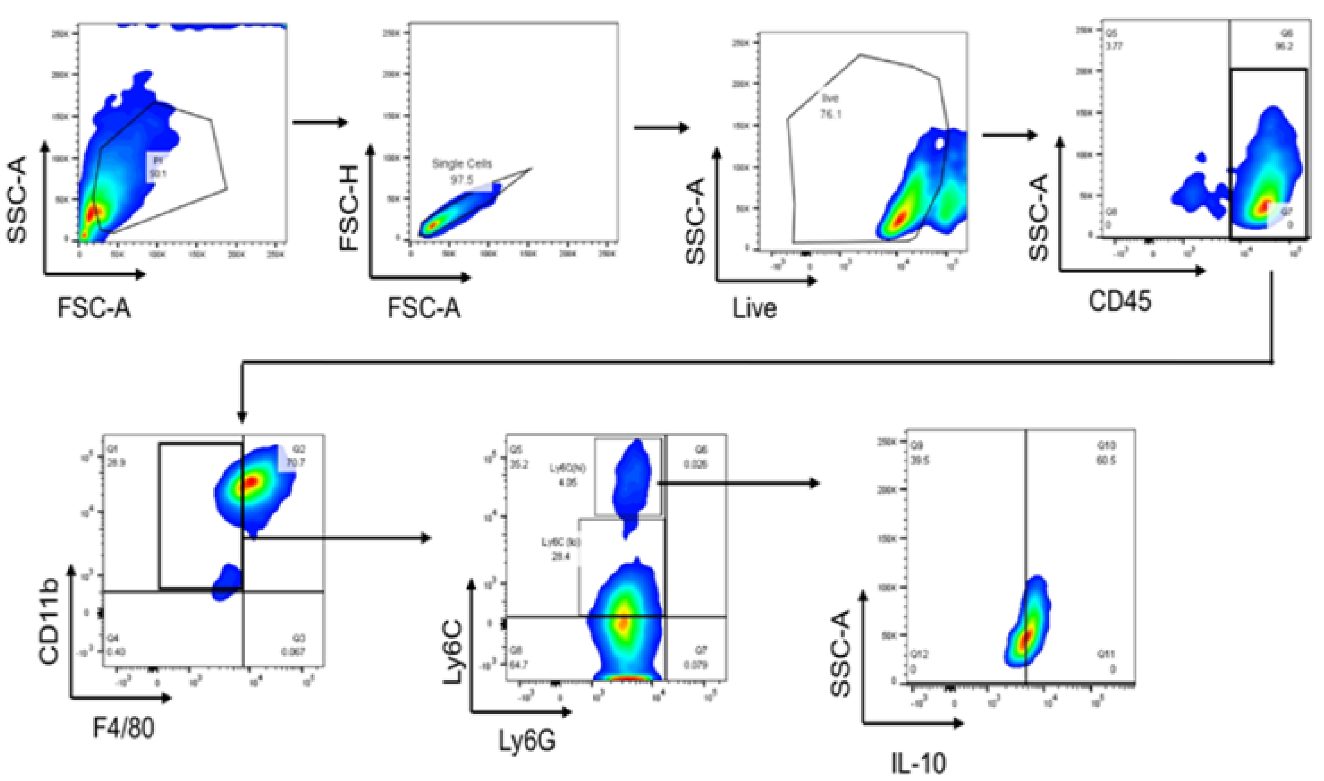

**Supplementary Figure S8. Gating scheme showing identification of myeloid-derived suppressor cell populations in primary mouse macrophages after BCG exposure.** Leukocyte lineage was determined on live cells by gating SSC-A against CD45+ myeloid cells. Myeloid cells were differentiated into $CD11b^{+}F4/80^{+}$ macrophages out of which $CD11b^{+}F4/80^{-}$ myeloid population was divided into Ly6C and Ly6G. Next, the $Ly6C^{(hi)}Ly6G^{-}$ immunosuppressive myeloid-derived suppressor cell populations were looked for IL-10 positivity (Related to **Fig. 6d-e**).

#### Supplementary Figure S9

**a**

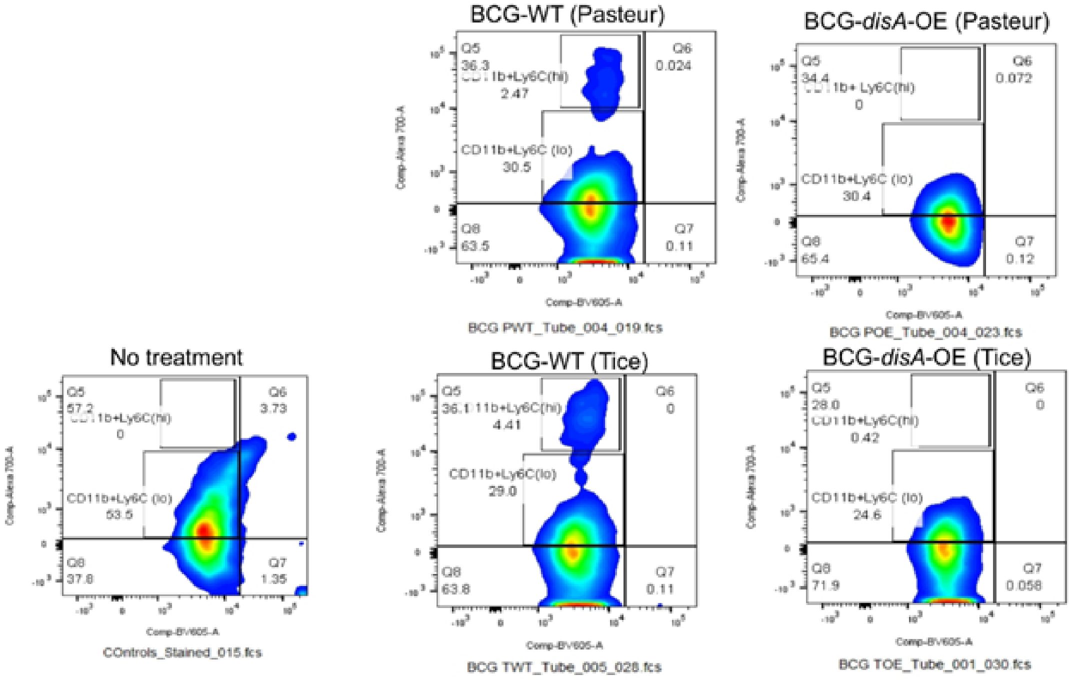

**b**

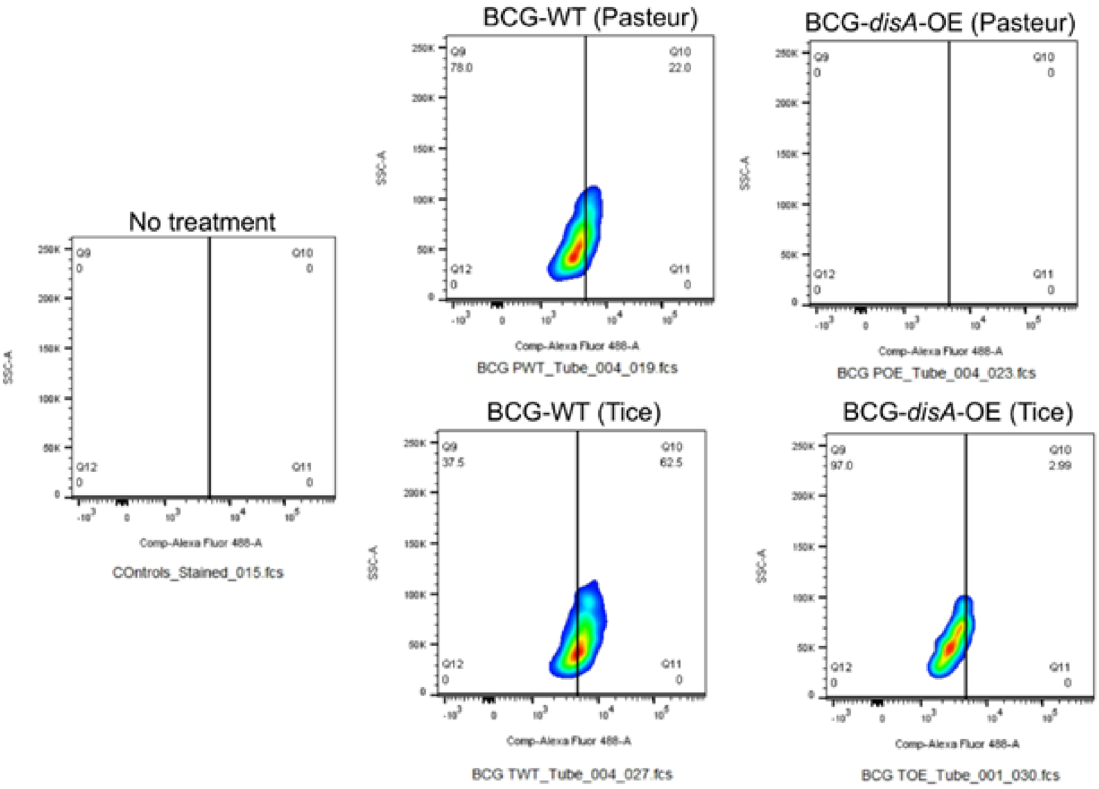

**Supplementary Figure S9. Immunosuppressive monocytic-MDSCs (M-MDSCs) populations murine primary macrophages after BCG exposure. a.** Representative FACS plots for M-MDSC measurements corresponding to **Fig. 6d**. **b.** Representative FACS plots for IL-10$^{+}$ expressing M-MDSCs corresponding to **Fig. 6e**.

### Supplementary Figure S10

**a**

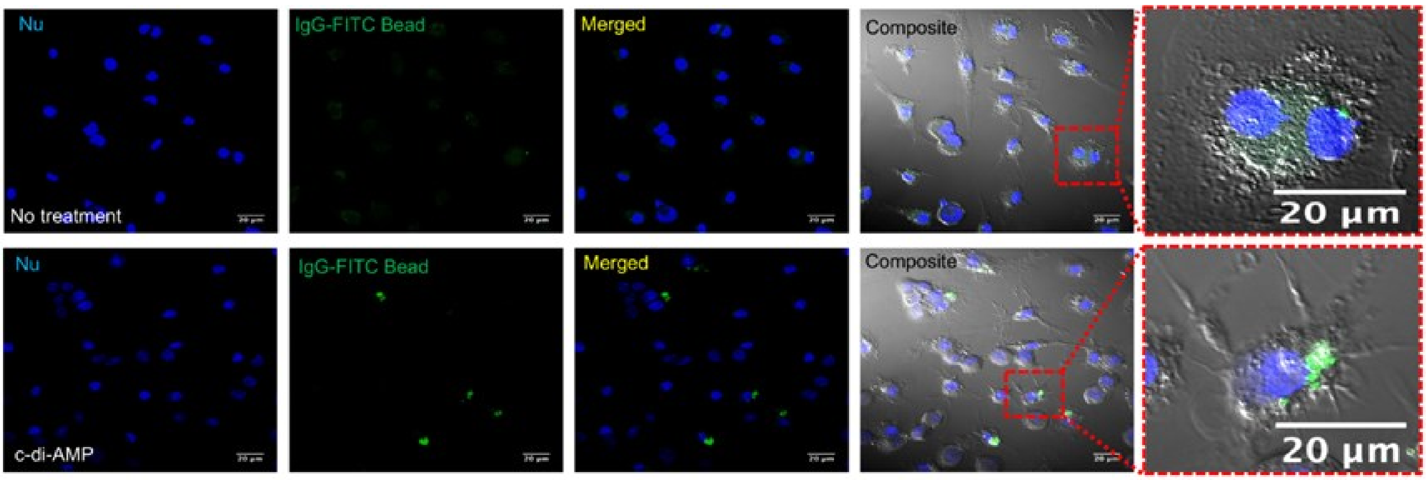

**Supplementary Figure S10. The STING agonist c-di-AMP causes induction of macrophage activation.** Human macrophages were transfected with c-di-AMP for 24 h and phagocytosis of FITC-labeled IgG opsonized latex beads (green) was visualized using confocal microscopy on live cells. Hoechst was used for nuclear staining (blue). Images were acquired using LSM700 confocal microscope at 63X magnification. Images were process using Fiji software. Similar results were observed across two (n=2) independent biological replicate experiments.

#### Supplementary Figure S11

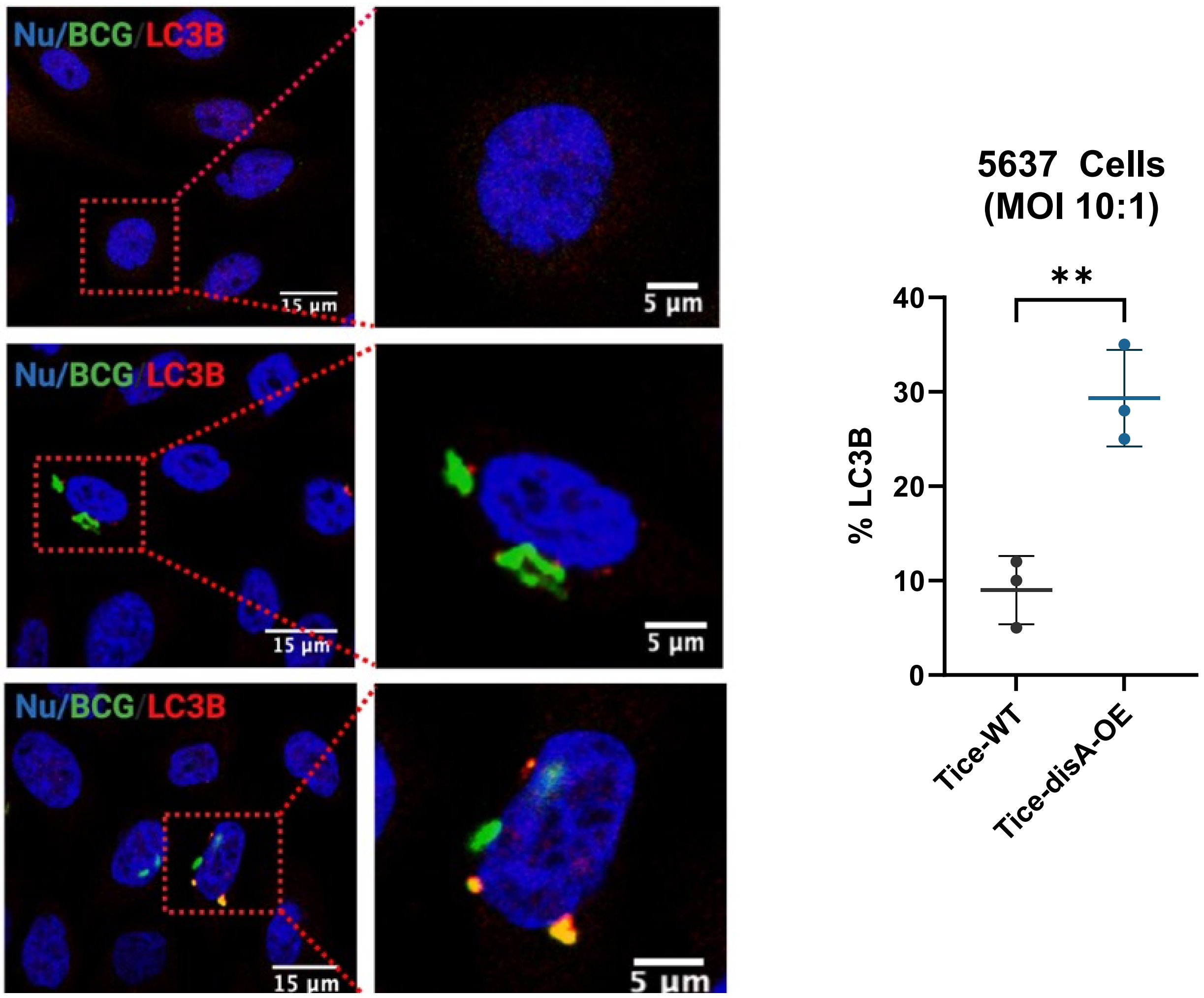

**Supplementary Figure S11. BCG-*disA*-OE elicits greater autophagy induction than BCG-WT in 5637 human urothelial carcinoma cells.** Autophagy induction in the 5637 human urothelial carcinoma cells in representative confocal photomicrographs. Co-localization of FITC-labeled BCG strains (green), LC3B autophagic puncta (red) appears in yellow; nuclei are blue. Quantification of co-localized BCG and LC3b puncta is shown at right. Cells were fixed using 4% paraformaldehyde 3h after infection (MOI 10:1), and images obtained with an LSM700 confocal microscope and Fiji software processing. Statistical analyses done using 2-tailed Student's t-test (** $p < 0.01$). Data shown are for BCG strains in the Tice background.

#### Supplementary Figure S12

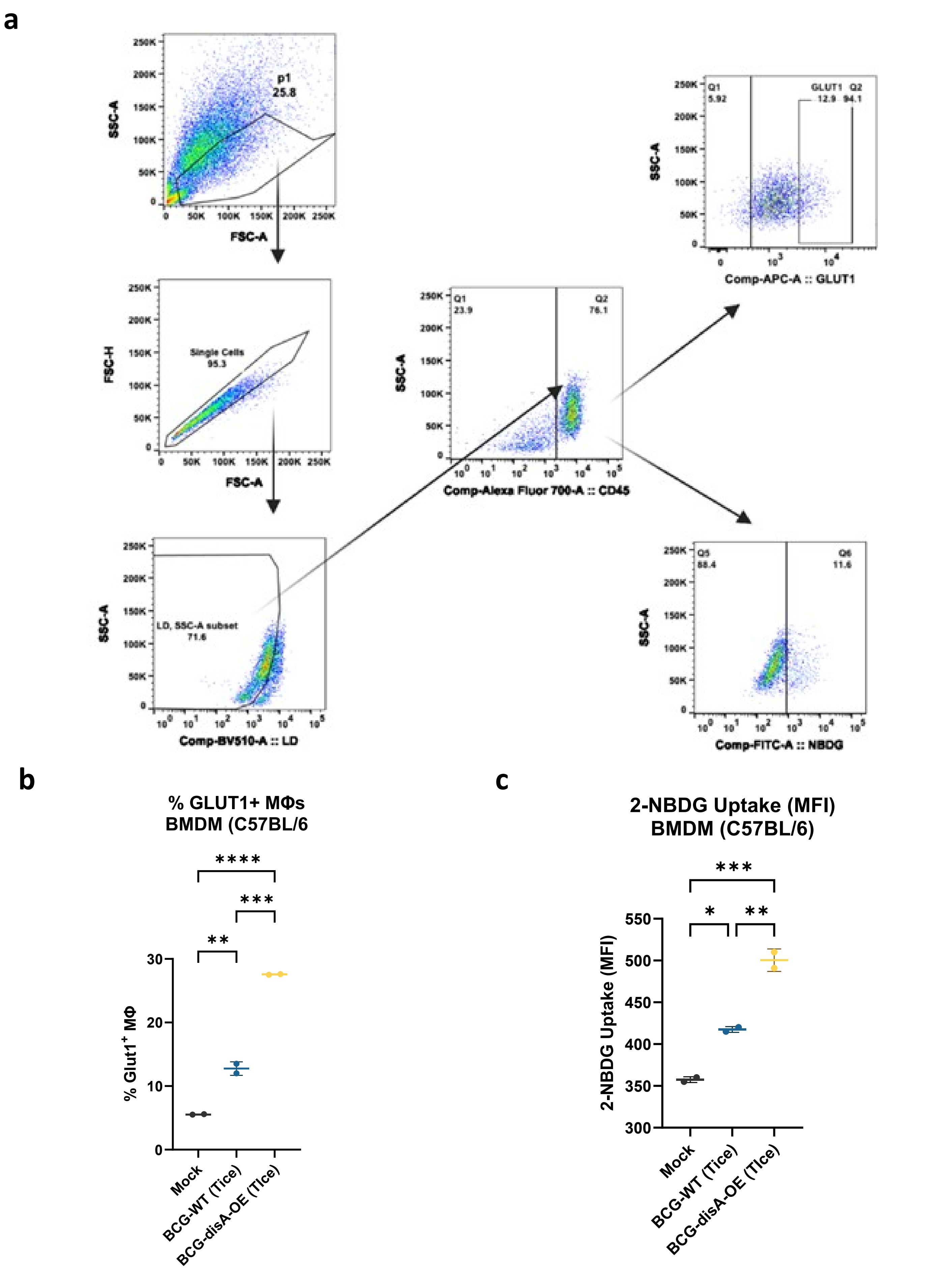

**Supplementary Figure S12. BCG induced differential glucose uptake in bone-marrow-derived macrophages (BMDMs). (a)** Experimental layout showing the strategy employed to determine intracellular uptake of fluorescent glucose. Briefly macrophages were infected at an MOI of 20:1 (BCG to macrophage ratio) in the presence of glucose-free medium followed by exogenous addition of 2-(*N*-(7-Nitrobenz-2-oxa-1,3-diazol-4-yl)Amino)-2-Deoxyglucose (2-NBDG). Macrophages were subsequently stained for GLUT1 and were investigated using flow cytometry. **(b-c)** Bar diagram showing induced expression of GLUT1 and intracellular fluorescent 2-NBDG in BMDMs following infection by BCG strains. Data are presented as mean values ± S.D. (n=2 independent biological replicate experiments). Data analyses were carried out using FACSDiva (v 9.0), Flowjo (v 10) and Graphpad Prism software (v 10.0.3). Statistical analysis employed a one-way ANOVA with Tukey's test for multiple comparisons (* $p < 0.05$, ** $p < 0.01$, ***$p < 0.001$, **** $p < 0.0001$).

**Supplementary Table. S1: List of bacterial strains, cell lines, plasmids and antibodies used in the study**

| Name | Description/Source |
|---|---|
| **Bacterial strains** | |
| ***M. tuberculosis* strain** | |
| Mtb-CDC1551 | Wild-type *M. tuberculosis* |
| ***M. bovis* BCG strains** | |
| BCG Pasteur | *M. bovis* BCG Pasteur |
| BCG-*disA*-OE (Pasteur) | BCG Pasteur strain overexpressing *disA* (MT3692) of *M.tb* |
| BCG Tice | *M. bovis* BCG Tice |
| BCG-*disA*-OE (Tice) | BCG Tice strain overexpressing *disA* (MT3692) of *M.tb* |
| ***E. coli* strain** | |
| DH5-$\alpha$ | Competent *E. coli* (High Efficiency) |
| **Cell lines** | |
| **Urinary bladder carcinoma cells** | |
| RT4 (ATCC ® HTB-2™) | Human low grade urothelial cancer |
| 5637 (ATCC® HTB-9™) | Human high-grade urothelial cancer |
| NBT-II (ATCC® CRL-1655™) | N-butyl-N-(4-hydroxybutyl) nitrosamine induced tumor cell line in Rattus norvegicus Nara Bladder Tumor No. 2 |
| MB49 (Cat. SSC148, EMD Millipore) | DMBA [7,12-dimethylbenz[a]anthracene] induced murine urothelial carcinoma cells, |
| UPPL-1595 | Luminal cell line established from a spontaneous primary bladder tumor in an Uroplakin-Cre driven PTEN/P53 knockout genetically engineered mouse model |
| BBN 975 | Basal- cell line established from 0.05% N-Butyl-N-(4-hydroxybutyl) nitrosamine (BBN) induced murine urothelial cancer model |
| J28 (ATCC® HTB-1™) | high grade urothelial cancer |
| **Reporter cells** | |
| RAW-Lucia ISG (InvivoGen) | IFN Reporter Raw 264.7 murine macrophages |
| **Macrophage cell lines** | |

| | | | |
|---|---|---|---|
| J774A.1 (ATCC® TIB67™) | Murine macrophage cell line | | |
| **Plasmids** | | | |
| pSD5.hsp60 | Mycobacterial expression plasmid with hsp60 promoter | | |
| pSD5hsp60.MT3692 | *disA* over-expression plasmid | | |
| **Confocal Microscopy Reagents** | | | |
| **Primary Antibodies** | | | |
| **Name** | **Source** | **#Cat.** | **Dilution** |
| LC3B | Novus Biologicals | NB100-2220 | 1:200 |
| P62/SQSTM1 | Sigma-Aldrich | P0067 | 1:100 |
| **Secondary Antibodies** | | | |
| Goat anti-Rabbit IgG Alexa Fluor Plus 647 | Thermo Fisher Scientific | A32733 | 1:1000 |
| **Chemicals/Probes** | | | |
| Fluorescein 5(6)-isothiocyanate (FITC) | Sigma-Aldrich | 46950 | |
| Hoechst 33342 | Thermo Fisher Scientific | 62249 | |
| **Flow Cytometry Reagents** | | | |
| **Antibodies (mouse BMDM study)** | | | |
| anti-CD45 (clone 30-F11) | Biolegend | 103128 | 1:200 |
| anti-CD124 (I clone 015F8) | Biolegend | 144804 | 1:50 |
| anti-I-A/I-E (clone M5/114.15.2) | Biolegend | 107645 | 1:200 |
| anti-Ly6C (clone HK1.4) | Biolegend | 128046 | 1:50 |
| anti-CD11b (clone M1/70) | Biolegend | 101206 | 1:200 |
| anti-F4/80 (clone BM8) | Biolegend | 123147 | 1:50 |
| anti-Ly6G (clone 1A8) | Biolegend | 127641 | 1:100 |
| anti CD206 (clone C068C2) | Biolegend | 141721 | 1:200 |
| anti-TNF (clone MP6-XT22) | Biolegend | 506341 | 1:100 |
| anti- IL-10 (clone JES5-16E3) | eBioscience | 505021 | 1:50 |

| | | | |
|---|---|---|---|
| Anti-Glut1 (clone EPR3915) | Abcam | ab195020 | 1:50 |
| **Antibodies (HMDM study)** | | | |
| anti-CD16 (clone 3G8) | Biolegend | 302028 | 1:50 |
| anti-CD14 (clone 63D3) | Biolegend | 367113 | 1:50 |
| anti-HLA-DR (clone L243) | Biolegend | 307615 | 1:50 |
| anti-CD11b (clone ICRF44) | Biolegend | 301351 | 1:100 |
| anti-TLR4 (clone HTA125) | Biolegend | 312811 | 1:50 |
| anti-CD206 (clone 15-2) | Biolegend | 321140 | 1:50 |
| anti-CD163 (clone GHI/61) | Biolegend | 333630 | 1:100 |
| anti-TNF (clone MAb11) | Biolegend | 502948 | 1:40 |
| anti-IL-6 (clone MQ2-13A5) | Biolegend | 501107 | 1:40 |
| **Antibodies (myeloid cell panel, Syngeneic MB49 urothelial cancer model)** | | | |
| CD45 (clone 30-F11) | Biolegend | 103128 | 1:100 |
| CD124 (IL-4Ra) (clone I015F8) | Biolegend | 144804 | 1:50 |
| I-a/I-e (clone M5/114.15.2) | Biolegend | 107645 | 1:100 |
| F4/80 (clone BM8) | Biolegend | 123147 | 1:100 |
| CD206 (clone C068C2) | Biolegend | 141721 | 1:200 |
| TNF (clone MP6-XT22) | Biolegend | 506345 | 1:40 |
| IL-10 (clone JES5-16E3) | Thermo Fisher | 505022 | 1:50 |
| **Antibodies (lymphoid cell panel, Syngeneic MB49 urothelial cancer model)** | | | |
| CD45 (clone 30-F11) | Biolegend | 103128 | 1:200 |
| CD25 (clone PC61) | Biolegend | 102033 | 1:100 |
| CD3 (clone 17A2) | Biolegend | 100248 | 1:50 |
| CD4 (clone GK1.5) | Biolegend | 100434 | 1:100 |
| CD8a (clone 53-6.7) | Biolegend | 100741 | 1:100 |
| FOXP3 (clone MF-14) | Biolegend | 126406 | 1:50 |
| Mouse IFN-γ (clone XMG1.2) | Biolegend | 505835 | 1:50 |

| | | | |
|---|---|---|---|
| CD69 (clone H1.2F3) | Biolegend | 104536 | 1:100 |
| CD38 (clone 90) | Biolegend | 102712 | 1:100 |
| **Reagents/Kits** | | | |
| **Name** | **Source** | **#Cat.** | |
| Protein transport inhibitor cocktail | eBioscience | 00-4980-03 | |
| Zombie Aqua™ Fixable Viability Kit | Biolegend | 423101 | |
| TruStain FcX™ | Biolegend | 101320 | |
| Fixation and Permeabilization Buffer Set | Biolegend | 421403 | |
| Human TruStain FcX™ | Biolegend | 422302 | |
| True-Stain Monocyte Blocker™ | Biolegend | 426102 | |
| 2-NBDG Glucose Uptake Assay Kit | Abcam | ab235976 | |
| **ELISA** | | | |
| **Mouse ELISA Kits** | | | |
| TNF- DuoSet | R and D Systems | DY410 | |
| IL-6 DuoSet | R and D Systems | DY406 | |
| IFN- DuoSet | R and D Systems | DY485 | |
| CCL2/JE/MCP-1 DuoSet | R and D Systems | DY479 | |
| LEGEND MAX™ Mouse IFN-β | Biolegend | 439407 | |
| **Human ELISA Kits** | | | |
| TNF- DuoSet | R and D Systems | DY210 | |
| IL-6 DuoSet | R and D Systems | DY206 | |
| IFN-β ELISA Kit | PBL Assay Science | 41410-2 | |
| **Rat ELISA Kits** | | | |
| IFN- Quantikine | R and D Systems | RIF00 | |
| TNF- Quantikine | R and D Systems | RTA00 | |

| | | |
|---|---|---|
| IL-2 Quantikine | R and D Systems | R2000 |
| **Chromatin Immunoprecipitation** | | |
| **ChIP Antibodies** | | |
| Histone H3K9me3 (H3K9 Trimethyl) Polyclonal Antibody | epigentek | A-4036-100 |
| Anti-Histone H3 (tri methyl K4) antibody - ChIP Grade | abcam | ab8580 |
| **ChIP Reagents** | | |
| BSA | Sigma-Aldrich | A3294 |
| Salmon Sperm DNA | Thermo Fisher Scientific | 15632011, |
| HEPES | Sigma-Aldrich | H3375 |
| Formaldehyde | Sigma-Aldrich | 252549 |
| EGTA | Sigma-Aldrich | 03777 |
| EDTA | Sigma-Aldrich | E6758 |
| TritonX-100 | Sigma-Aldrich) | T8787 |
| SDS | Sigma-Aldrich | 71736 |
| NaHCO3 | Sigma-Aldrich | 5761 |
| Nuclease free water | Thermo Fisher Scientific | AM9930 |
| SYBR green dye | Applied Biosystems | 4385614 |

**Supplementary Table S2: Cloning and PCR primers used in the study**

| **Cloning primers used in the study** | | |
|---|---|---|
| **Accession Number** | **Gene** | **Sequence (5’-3’)** |
| | pSD5hsp60.MT3692 (F) | GGGCATCATATGCACGCTGTGACTCGTC |
| | pSD5hsp60.MT3692 (R) | GGGACGCGTTATTGATCGCTGATGGTCGATT |
| | Kanamycin cassette (F) | GAGAAAACTCACCGAGGCAG |
| | Kanamycin cassette (R) | GTATTTCGTCTCGCTCAGGC |
| 32287254 | *M.tb* sigH (F) | GCGATGGTGGCTTCTCCCTCG |
| | *M.tb* sigH (R) | CCATCTTGCACAGCTCGCGTAG |
| **qPCR primers used in the study** | | |
| **Mouse Primers** | | |
| 11461 | Mouse.β actin (F) | TAAGGCCAACCGTGAAAAGATG |
| | Mouse.β actin (R) | CTGGATGGCTACGTACATGGCT |
| 21926 | Mouse.TNF-α (F) | GACCCTCACACTCAGATCATC |
| | Mouse.TNF-α (R) | GCTGCTCCTCCACTTGGT |
| 15977 | Mouse.IFN-β (F) | CCACAGCCCTCTCCATCAAC |
| | Mouse.IFN-β (R) | CTCCGTCATCTCCATAGGGA |
| 16193 | Mouse.IL6 (F) | CTGCAAGAGACTTCCATCCAG |
| | Mouse.IL6 (R) | CAGGTCTGTTGGGAGTGG |
| 15978 | Mouse.IFN (F) | AGCGGCTGACTGAACTCAGATTGT |
| | Mouse.IFN (R) | GTCACAGTTTTCAGCTGTATAGGG |
| 16176 | Mouse.IL1 (F) | GGAGAGTGTGGATCCCAA |
| | Mouse.IL1 (R) | GTGGAGTTTGAGTCTGCAG |
| 20296 | Mouse.MCP1 (F) | GGCTCAGCCAGATGCAGTTAAC |
| | Mouse.MCP1 (R) | GATCCTCTTGTAGCTCTCCAGC |
| 16160 | Mouse.IL12b (F) | GAAAGACGTTTATGTTGTAGAGG |
| | Mouse.IL12b (R) | GACTCCATGTCTCTGGTCTG |
| 17329 | Mouse.CXCL9 (F) | GGAGTTCGAGGAACCCTAGTG |

| | | |
|---|---|---|
| | Mouse.CXCL9 (R) | GGGATTTGTAGTGGATCGTGC |
| 15945 | Mouse.CXCL10 (F) | GTGGGACTCAAGGGATCCCTCTC |
| | Mouse.CXCL10 (R) | GCTTCCCTATGGCCCTCATTC |
| 18126 | Mouse.NOS2 (F) | GTTCTCAGCCCAACAATACAAG |
| | Mouse.NOS2 (R) | GGAACATTCTGTGCTGTCCC |
| 20299 | Mouse.CCL22 (F) | CTCTGATGCAGGTCCCTATGGTG |
| | Mouse.CCL22 (R) | GGCAGAGGGTGACGGATGTAG |
| **Human Primers** | | |
| 26827 | Human. RNU6A (F) | CTCGCTTCGGCAGCACATATAC |
| | Human. RNU6A (R) | AATATGGAACGCTTCACGAATTTG |
| 3456 | Human.IFNβ (F) | CAACTTGCTTGGATTCCTACAAAG |
| | Human.IFNβ (R) | TATTCAAGCCTCCCATTCAATTG |
| 3569 | Human.IL6 (F) | GGTACATCCTCGACGGCATCT |
| | Human.IL6 (R) | GTGCCTCTTTGCTGCTTTCAC |
| **Rat Primers** | | |
| 64367 | Rat.PPIB (F) | CAGGATTCATGTGCCAGGGT |
| | Rat.PPIB (R) | CCAAAGACCACATGCTTGCC |
| 24481 | Rat.IFN-β (F) | GAGTCTTCACACTCCTGGC |
| | Rat.IFN-β (R) | GTCCTTCAGGCATGAGACAG |
| 298210 | Rat.IFN–α (F) | GCGTTCCTGCTGTGCTTCTC |
| | Rat.IFN-α (R) | CCATTCAGCTGCCTCAGGAGC |
| 25712 | Rat.IFN–γ (F) | CGTCTTGGTTTTGCAGCTCT |
| | Rat.IFN-γ (R) | CGTCCTTTTGCCAGTTCCTC |
| 24599 | Rat. iNOS (F) | GGTGAGGGGACTGGACTTTTAG |
| | Rat. iNOS (R) | TTGTTGGGCTGGGAATAGCA |
| 245920 | Rat.IP10 (F) | TCCACCTCCCTTTACCCAGT |
| | Rat.IP10 (R) | AGAGCTAGGAGAGCCGTCAT |
| 24770 | Rat.MCP-1 (F) | CAGGTCTCTGTCACGCTTCTG |
| | Rat.MCP-1 (R) | GCCAGTGAATGAGTAGCAGCAG |
| 25542 | Rat.MIP-1α (F) | ACAAGCGCACCCTCTGTTAC |
| | Rat.MIP-1α (R) | GGTCAGGAAAATGACACCCG |
| 24494 | Rat.IL-1β (F) | GACTTCACCATGGAACCCGT |

| | | |
|---|---|---|
| | Rat.IL-1β (R) | GGAGACTGCCCATTCTCGAC |
| 24835 | Rat.TNF-α (F) | CGTCCCTCTCATACACTGG |
| | Rat.TNF-α (R) | CATGCTTTCCGTGCTCATG |
| 59086 | Rat.TGF-β (F) | TGACGTCACTGGAGTTGTCC |
| | Rat.TGF-β (R) | CCTCGACGTTTGGGACTGAT |
| 25325 | Rat.IL-10 (F) | CCTCTGGATACAGCTGCGAC |
| | Rat.IL-10 (R) | TGCCGGGTGGTTCAATTTTTC |
| ChIP-PCR Primers | | |
| | Human.GAPDH (F) | TACTAGCGGTTTTACGGGCG |
| | Human.GAPDH (R) | TCGAACAGGAGGAGCAGAGAGCGA |
| | Human.IL-6 (F) | CGGTGAAGAATGGATGACCT |
| | Human.IL-6 (R) | AAACGAGACCCTTGCACAAC |
| | Human.TNF-α (F) | ATCAGTCAGTGGCCCAGAAGACCC |
| | Human.TNF-α (R) | CCACGTCCCGGATCATGCTTCAG |

### Source Data

Improved bladder cancer antitumor efficacy with a recombinant BCG that releases a STING agonist

### Figure 2b (i)

**MNU Rat Bladder (Post BCG Immunotherapy) qPCR**
**Fold Expression (Log2)**
**IFN-β gene**

| | *Ifnb* | | | |
|---|---|---|---|---|
| | Control | Untreated | BCG-WT (Tice) | BCG-*disA* -OE (Tice) |
| Animal 1 | 0.9462 | 0.4323 | 0.3983 | 3.3258 |
| Animal 2 | 0.7949 | 0.8570 | 0.3026 | 4.0350 |
| Animal 3 | 1.4769 | 0.8532 | 0.3885 | 6.7943 |
| Animal 4 | 1.6185 | 0.8145 | 0.3947 | 2.4513 |
| Animal 5 | 0.7193 | 0.4325 | 0.4030 | 2.4153 |

**MNU Rat Bladder (Post BCG Immunotherapy) qPCR**
**Fold Expression (Log2)**
**IFN-γ gene**

| | *Ifng* | | | |
|---|---|---|---|---|
| | Control | Untreated | BCG-WT (Tice) | BCG-*disA* -OE (Tice) |
| Animal 1 | 1.3668 | 0.1644 | 0.5392 | 3.6505 |
| Animal 2 | 0.7741 | 0.1330 | 0.6280 | 4.7526 |
| Animal 3 | 1.5244 | 0.2770 | 0.4786 | 2.6071 |
| Animal 4 | 0.9933 | 0.1577 | 0.5626 | 2.5401 |
| Animal 5 | 0.9765 | 0.2184 | 0.6620 | 2.0162 |

**MNU Rat Bladder (Post BCG Immunotherapy) qPCR**
**Fold Expression (Log2)**
**TNF-α gene**

| | *Tnfa* | | | |
|---|---|---|---|---|
| | Control | Untreated | BCG-WT (Tice) | BCG-*disA* -OE (Tice) |
| Animal 1 | 1.0537 | 1.1987 | 2.2263 | 5.0408 |
| Animal 2 | 0.8277 | 1.5919 | 4.9580 | 5.0596 |
| Animal 3 | 1.1050 | 2.7742 | 4.7815 | 5.1397 |
| Animal 4 | 1.1006 | 2.8183 | 2.2567 | 4.6265 |
| Animal 5 | 0.9130 | 2.5765 | 4.9526 | 4.7647 |

### Figure 2b (ii)

**MNU Rat Bladder (Post BCG Immunotherapy) qPCR**
**Fold Expression (Log2)**
**IL-1β gene**

| | *Il1b* | | | |
|---|---|---|---|---|
| | Control | Untreated | BCG-WT (Tice) | BCG-*disA* -OE (Tice) |
| Animal 1 | 1.6453 | 1.3032 | 4.0929 | 3.8334 |
| Animal 2 | 1.4692 | 1.0493 | 1.5417 | 5.7733 |
| Animal 3 | 0.2676 | 2.3949 | 2.4456 | 5.4485 |
| Animal 4 | 1.3872 | 3.6081 | 1.8949 | 5.9633 |
| Animal 5 | 0.2308 | 3.6083 | 1.8901 | 5.9916 |

**MNU Rat Bladder (Post BCG Immunotherapy) qPCR**
**Fold Expression (Log2)**
**CXCL10 gene**

| | *Cxcl10* | | | |
|---|---|---|---|---|
| | Control | Untreated | BCG-WT (Tice) | BCG-*disA* -OE (Tice) |
| Animal 1 | 1.3507 | 0.1813 | 0.2843 | 2.4623 |
| Animal 2 | 0.8683 | 0.1763 | 0.2710 | 2.7094 |
| Animal 3 | 0.9417 | 0.1734 | 0.5572 | 2.3360 |
| Animal 4 | 0.8394 | 0.2330 | 0.3326 | 4.4101 |
| Animal 5 | 1.1473 | 0.3051 | 0.2727 | 2.9344 |

**MNU Rat Bladder (Post BCG Immunotherapy) qPCR**
**Fold Expression (Log2)**
**MCP-1 gene**

| | *Mcp1* | | | |
|---|---|---|---|---|
| | Control | Untreated | BCG-WT (Tice) | BCG-*disA* -OE (Tice) |
| Animal 1 | 1.2807 | 0.4347 | 0.3453 | 2.8489 |
| Animal 2 | 1.0746 | 0.5138 | 0.4674 | 3.1102 |
| Animal 3 | 1.0435 | 0.507 | 0.3142 | 4.7347 |
| Animal 4 | 0.8860 | 0.6558 | 0.2760 | 4.4665 |
| Animal 5 | 0.7153 | 0.6598 | 0.2898 | 2.1945 |

### Figure 2b (iii)

**MNU Rat Bladder (Post BCG Immunotherapy) qPCR**
**Fold Expression (Log2)**
**MIP-1$\alpha$ gene**

| | *Mip1a* | | | |
|---|---|---|---|---|
| | Control | Untreated | BCG-WT (Tice) | BCG-*disA* -OE (Tice) |
| Animal 1 | 1.1379 | 0.1872 | 0.3376 | 2.1696 |
| Animal 2 | 0.8621 | 0.1066 | 0.3569 | 1.8937 |
| Animal 3 | 0 | 0.1557 | 0.3246 | 1.0737 |
| Animal 4 | 0.8267 | 0.1587 | 0.2431 | 1.8990 |
| Animal 5 | 1.0137 | 0.1222 | 0.3550 | 1.2796 |

**MNU Rat Bladder (Post BCG Immunotherapy) qPCR**
**Fold Expression (Log2)**
**IL-10 gene**

| | *Il10* | | | |
|---|---|---|---|---|
| | Control | Untreated | BCG-WT (Tice) | BCG-*disA* -OE (Tice) |
| Animal 1 | 1.4684 | 6.1882 | 4.7483 | 2.0002 |
| Animal 2 | 0.6358 | 7.2067 | 4.1929 | 2.9396 |
| Animal 3 | 1.6621 | 6.8448 | 1.7800 | 2.5655 |
| Animal 4 | 0.6174 | 6.3872 | 3.0441 | 3.8789 |
| Animal 5 | 0.6163 | 6.2583 | 3.0442 | 3.8785 |

**MNU Rat Bladder (Post BCG Immunotherapy) qPCR**
**Fold Expression (Log2)**
**TGF-$\beta$ gene**

| | *Tgfb* | | | |
|---|---|---|---|---|
| | Control | Untreated | BCG-WT (Tice) | BCG-*disA* -OE (Tice) |
| Animal 1 | 1.0998 | 10.7662 | 2.2188 | 2.1424 |
| Animal 2 | 0.8343 | 6.5788 | 3.2697 | 2.2181 |
| Animal 3 | 1.1785 | 7.2088 | 2.7960 | 5.5651 |
| Animal 4 | 0.9414 | 6.7679 | 2.4813 | 2.5642 |
| Animal 5 | 0.9459 | 6.3653 | 2.4116 | 2.5671 |

### Figure 2b (iv)

**MNU Rat Bladder (Post BCG Immunotherapy) qPCR**
**Fold Expression (Log2)**
**Nos2 gene**

| | *Nos2* | | | |
|---|---|---|---|---|
| | Control | Untreated | BCG-WT (Tice) | BCG-*disA* -OE (Tice) |
| Animal 1 | 1.1946 | 4.9202 | 8.9481 | 49.8755 |
| Animal 2 | 0.1469 | 3.9075 | 7.8584 | 34.1369 |
| Animal 3 | 0.7264 | 5.7335 | 8.9911 | 48.7897 |
| Animal 4 | 1.4347 | 3.6855 | 5.9194 | 56.3878 |
| Animal 5 | 1.4974 | | 5.7084 | 56.1033 |

### Figure 2d

**Tumor Involvement Index (MNU Rat Bladder Post BCG Immunotherapy)**

| | Untreated | BCG-WT (Tice) | BCG-*disA* -OE (Tice) |
|---|---|---|---|
| **Animal 1** | 600 | 0 | 0 |
| **Animal 2** | 2500 | 2700 | 200 |
| **Animal 3** | 200 | 0 | 400 |
| **Animal 4** | 200 | 0 | 200 |
| **Animal 5** | 500 | 300 | 0 |
| **Animal 6** | 300 | 1500 | 0 |
| **Animal 7** | 3500 | 100 | 0 |
| **Animal 8** | | 0 | 300 |
| **Animal 9** | | 500 | 400 |
| **Animal 10** | | 400 | 0 |
| **Animal 11** | | 0 | 0 |

**Tumor staging was performed by 2 board-certified genitourinary pathologists (A.S.B., A.M.) blinded to treatment groups. Specimens were classified based on the percentage of involvement of abnormal tissue.**

### Figure 2e

Histopathology
(MNU Rat Model of NMIBC)

| Animal | Pathologic Stage |
|---|---|
| CTRL 1 | HGT2 |
| CTRL 2 | CIS |
| CTRL 4 | CIS |
| CTRL 5 | CIS |
| CTRL 6 | CIS |
| BCG WT 1 | dysplasia |
| BCG WT 2 | HGT1 |
| BCG WT 3 | dysplasia |
| BCG WT 4 | CIS |
| BCG WT 5 | dysplasia |
| BCG OE 1 | dysplasia |
| BCG OE 2 | dysplasia |
| BCG OE 3 | CIS |
| BCG OE 4 | CIS |
| BCG OE 5 | dysplasia |

| Animal | Pathologic Stage |
|---|---|
| Control 1 | LGTa |
| Control 2 | HGT2 |
| Control 3 | LGTa, squamous metaplasia |
| Control 4 | HGT1 |
| Control 5 | CIS |
| Control 6 | CIS |
| WT TICE 1 | HGT1 |
| WT TICE 2 | CIS |
| WT TICE 3 | CIS |
| WT TICE 4 | CIS |
| WT TICE 5 | CIS |
| WT TICE 6 | dysplasia |
| OE TICE 1 | LGTa |
| OE TICE 2 | dysplasia |
| OE TICE 3 | CIS |
| OE TICE 4 | LGTa |
| OE TICE 5 | dysplasia |
| WT Pasteur1 | HGT1 (90%) squamous differentiation |
| WT Pasteur2 | CIS |
| WT Pasteur3 | dysplasia |
| WT Pasteur4 | LGTa |
| WT Pasteur5 | CIS |
| OE Pasteur 1 | dysplasia |
| OE Pasteur 2 | LGTa |
| OE Pasteur 3 | CIS |
| OE Pasteur 4 | dysplasia |
| OE Pasteur 5 | dysplasia |

OE = BCG-*disA* -OE

| Staging was performed by a blinded by a board-certified pathologist |
|---|
| dysplasia= abnormal appearing urothelium falling short of criteria for carcinoma in situ |
| CIS= carcinoma in situ |
| LGTa= low grade non invasive |
| HGTa = high grade non invasive |
| HGT1 = high grade invasive into the lamina propria |
| HGT2 = high grade invasive into the muscle |

### Figure 2g

| | Ki67 Staining (IHC) (MNU Rat Bladder Post BCG Immunotherapy) | | |
|---|---|---|---|
| | Untreated | BCG-WT (Tice) | BCG-*disA* -OE (Tice) |
| **Animal 1** | 15 | 20 | 10 |
| **Animal 2** | 40 | 20 | 15 |
| **Animal 2** | 15 | 10 | 20 |
| **Animal 4** | 40 | 10 | 5 |
| **Animal 5** | 15 | 20 | 5 |
| **Animal 6** | 20 | 30 | 10 |
| **Animal 7** | 30 | 10 | 5 |
| **Animal 8** | 20 | 10 | 20 |
| **Animal 9** | 30 | 20 | 15 |
| **Animal 10** | 20 | 20 | 2 |
| **Animal 11** | 15 | 10 | 5 |
| **Animal 12** | 30 | 10 | 5 |
| **Animal 13** | 10 | 20 | 20 |
| **Animal 14** | 30 | 10 | 15 |
| **Animal 15** | 20 | 30 | 10 |
| **Animal 16** | 10 | 10 | 10 |
| **Animal 17** | | 10 | 10 |
| **Animal 18** | | 20 | 20 |
| **Animal 19** | | 10 | 15 |
| **Animal 20** | | 30 | 10 |
| **Animal 21** | | 10 | |

All slides were scored blinded by board certified pathologist
Ki67 IHC Scores represent % of positive cells in urothelium

### Figure 2h

**CD68 IHC Score (MNU Rat Bladder Post BCG Immunotherapy)**

| | Untreated | BCG-WT (Tice) | BCG-*disA* -OE (Tice) |
|---|---|---|---|
| **Animal 1** | 2 | 1 | 3 |
| **Animal 2** | 1 | 3 | 2 |
| **Animal 3** | 1 | 1 | 2 |
| **Animal 4** | 2 | 3 | 2 |
| **Animal 5** | 2 | 1 | 2 |

**CD86 IHC Score (MNU Rat Bladder Post BCG Immunotherapy)**

| | Untreated | BCG-WT (Tice) | BCG-*disA* -OE (Tice) |
|---|---|---|---|
| **Animal 1** | 0 | 0 | 1 |
| **Animal 2** | 0 | 0 | 0 |
| **Animal 3** | 1 | 0 | 0 |
| **Animal 4** | 0 | 1 | 1 |
| **Animal 5** | 1 | 0 | 1 |

**CD206 IHC Score (MNU Rat Bladder Post BCG Immunotherapy)**

| | Untreated | BCG-WT (Tice) | BCG-*disA* -OE (Tice) |
|---|---|---|---|
| **Animal 1** | 2 | 0 | 0 |
| **Animal 2** | 1 | 1 | 0 |
| **Animal 3** | 1 | 1 | 1 |
| **Animal 4** | 1 | 1 | 0 |
| **Animal 5** | 1 | 0 | 0 |

| |
|---|
| **All slides were scored blinded by board certified pathologist** |
| **CD68, CD206, CD86 IHC SCORES** |
| **1= Positive isolated cells** |
| **2= Clusters of up to 10 positive cells** |
| **3= Greater than 10 positive cells/cluster** |

### Figure 3b

**Tumor Volume (mm3) (Mouse MB49 Tumors, C57BL/6 Female Mice)**
**Syngeneic MB49 Model of Urothelial Cancer**

| | Vehicle Animal 1 | Vehicle Animal 2 | Vehicle Animal 3 | Vehicle Animal 4 | Vehicle Animal 5 | Vehicle Animal 6 | Vehicle Animal 7 | Vehicle Animal 8 |
|---|---|---|---|---|---|---|---|---|
| **D1** | 0 | 0 | 0 | 0 | 0 | 0 | 0 | 0 |
| **D10** | 87.99 | 78.96 | 28.42 | 57.14 | 47.07 | 47.01 | 40.95 | 60.00 |
| **D14** | 425.38 | 201.34 | 379.94 | 322.70 | 165.39 | 250.57 | 212.55 | 216.12 |
| **D18** | 669.83 | 221.31 | 660.34 | 495.39 | 165.58 | 514.45 | 496.02 | 248.55 |
| **D22** | 1041.45 | 316.99 | 783.46 | 463.34 | 250.82 | 643.67 | 446.31 | 210.12 |

| | BCG-WT (Pasteur) Animal 1 | BCG-WT (Pasteur) Animal 2 | BCG-WT (Pasteur) Animal 3 | BCG-WT (Pasteur) Animal 4 | BCG-WT (Pasteur) Animal 5 | BCG-WT (Pasteur) Animal 6 | BCG-WT (Pasteur) Animal 7 | BCG-WT (Pasteur) Animal 8 |
|---|---|---|---|---|---|---|---|---|
| **D1** | 0 | 0 | 0 | 0 | 0 | 0 | 0 | 0 |
| **D10** | 71.30 | 69.26 | 58.60 | 56.14 | 48.06 | 46.52 | 33.95 | 21.08 |
| **D14** | 155.32 | 158.67 | 140.94 | 168.40 | 180.66 | 166.57 | 160.94 | 136.87 |
| **D18** | 213.56 | 237.50 | 238.94 | 365.81 | 97.51 | 212.90 | 196.10 | 105.52 |
| **D22** | 252.24 | 280.65 | 251.85 | 444.25 | 32.76 | 221.53 | 213.04 | 181.35 |

| | BCG-*disA*-OE (Pasteur) Animal 1 | BCG-*disA*-OE (Pasteur) Animal 2 | BCG-*disA*-OE (Pasteur) Animal 3 | BCG-*disA*-OE (Pasteur) Animal 4 | BCG-*disA*-OE (Pasteur) Animal 5 | BCG-*disA*-OE (Pasteur) Animal 6 | BCG-*disA*-OE (Pasteur) Animal 7 | BCG-*disA*-OE (Pasteur) Animal 8 |
|---|---|---|---|---|---|---|---|---|
| **D1** | 0 | 0 | 0 | 0 | 0 | 0 | 0 | 0 |
| **D10** | 102.81 | 74.22 | 32.79 | 54.46 | 48.86 | 45.89 | 38.24 | 58.00 |
| **D14** | 79.38 | 109.66 | 110.68 | 43.96 | 104.04 | 122.23 | 200.26 | 63.15 |
| **D18** | 135.72 | 174.26 | 193.18 | 41.82 | 168.83 | 150.17 | 93.63 | 83.34 |
| **D22** | 72.14 | 156.35 | 174.09 | 56.27 | 194.91 | 109.07 | 151.79 | 198.25 |

### Figure 3c

**% CD3+ of CD45 cells (MB49 Tumors)**
**Syngeneic MB49 heterotopic tumor model)**
**Flow Cytometry Analyses**

| Animal | C57BL/6 WT + Saline | STING$^{-/-}$ (C57BL/6J-Tmem173gt/J) + Saline | C57BL/6 WT + BCG-WT (Tice) | STING$^{-/-}$ (C57BL/6J-Tmem173gt/J) + BCG-WT (Tice) | C57BL/6 WT + BCG-*disA* -OE (Tice) | STING$^{-/-}$ (C57BL/6J-Tmem173gt/J) + BCG-*disA* -OE (Tice) |
|---|---|---|---|---|---|---|
| **A1** | 13.7 | 6.9 | 42.2 | 16.0 | 42.7 | 64.1 |
| **A2** | 11.3 | 7.1 | 21.2 | 16.5 | 36.4 | 16.2 |
| **A3** | 19.5 | 6.3 | 25.0 | 13.5 | 73.6 | 6.4 |
| **A4** | 14.2 | 8.9 | 28.9 | 19.8 | 34.4 | 14.3 |
| **A5** | 13.6 | 6.8 | 30.6 | 15.8 | 70.5 | 36.0 |
| **A6** | 14.1 | 8.0 | 35.0 | 21.1 | 23.9 | 16.8 |

*A: Animal*

### Figure 3d

**% CD25+CD69+ of CD8 T cells (MB49 Tumors)**
**Syngeneic MB49 heterotopic tumor model)**
**Flow Cytometry Analyses**

| Animal | C57BL/6 WT + Saline | STING$^{-/-}$ (C57BL/6J-Tmem173gt/J) + Saline | C57BL/6 WT + BCG-WT (Tice) | STING$^{-/-}$ (C57BL/6J-Tmem173gt/J) + BCG-WT (Tice) | C57BL/6 WT + BCG-*disA* -OE (Tice) | STING$^{-/-}$ (C57BL/6J-Tmem173gt/J) + BCG-*disA* -OE (Tice) |
|---|---|---|---|---|---|---|
| **A1** | 1.88 | 1.76 | 12.9 | 2.54 | 30.5 | 7.82 |
| **A2** | 0.97 | 0.38 | 0.58 | 18.5 | 25.4 | 4.17 |
| **A3** | 9.20 | 0.64 | 9.75 | 0.28 | 26.6 | 0 |
| **A4** | 1.60 | 0.80 | 6.48 | 9.50 | 23.7 | 4.58 |
| **A5** | 2.13 | 1.14 | 8.61 | 2.36 | 27.3 | 22.8 |
| **A6** | 1.09 | 0.21 | 13.6 | 17.3 | 19.1 | 19.3 |

*A: Animal*

### Figure 3e

**% TNF-$\alpha$+ of F4/80+CD11b+ cells (MB49 Tumors)**
**Syngeneic MB49 heterotopic tumor model)**
**Flow Cytometry Analyses**

| **Animal** | **C57BL/6 WT + Saline** | **STING$^{-/-}$ (C57BL/6J-Tmem173gt/J) + Saline** | **C57BL/6 WT + BCG-WT (Tice)** | **STING$^{-/-}$ (C57BL/6J-Tmem173gt/J) + BCG-WT (Tice)** | **C57BL/6 WT + BCG-*disA* -OE (Tice)** | **STING$^{-/-}$ (C57BL/6J-Tmem173gt/J) + BCG-*disA* -OE (Tice)** |
|---|---|---|---|---|---|---|
| **A1** | 11.6 | 5.9 | 29.1 | 19.6 | 36.2 | 12.6 |
| **A2** | 7.1 | 3.8 | 18.8 | 28.1 | 39.2 | 17.9 |
| **A3** | 10.3 | 5.1 | 28.6 | 12.5 | 33.8 | 9.81 |
| **A4** | 12.3 | 4.6 | 26.1 | 24.4 | 38.0 | 15.6 |
| **A5** | 10.4 | 3.0 | 28.2 | 16.8 | 34.1 | 13.7 |
| **A6** | 10.3 | 7.8 | 23.6 | 35.9 | 41.0 | 17.3 |

*A: Animal*

### Figure 4b

**Lung Bacillary Burden in BALB/c Mice after BCG Infection**
**D1 Implantation: raw CFU per lung and log10 CFU per lung**

**Half of total lung homogenate plated on each of 2 plates**

| CFU counts per plate (each animal lung plated in duplicate) (D1 implantation) | | | | |
|---|---|---|---|---|
| **Strain** | **Plate 1** | **Plate 2** | **CFU (total)** | **Log10cfu** |
| **BCG-WT (Tice)** | | | | |
| **A1** | 209 | 195 | 404 | 2.607 |
| **A2** | 109 | 172 | 281 | 2.450 |
| **A3** | 110 | 352 | 462 | 2.666 |
| **BCG-*disA* -OE (Tice)** | | | | |
| A1 | 378 | 73 | 451 | 2.655 |
| A2 | 376 | 66 | 462 | 2.666 |
| A3 | 214 | 110 | 324 | 2.512 |

**Lung Bacillary Burden in BALB/c Mice after BCG.**
**Infection D28 (log10 CFU per lung)**

| | **BCG-WT (Tice)** | **BCG-*disA* -OE (Tice)** |
|---|---|---|
| **Animal 1** | 6.243 | 5.933 |
| **Animal 2** | 6.347 | 5.861 |
| **Animal 3** | 6.294 | 5.834 |
| **Animal 4** | 6.142 | 5.774 |
| **Animal 5** | 6.251 | 5.704 |

### Figure 4d

**Lung Bacillary Burden in SCID Mice after BCG Infection D1 Implantation (log10 CFU per lung)**

| | BCG-WT (Tice) | BCG-*disA* -OE (Tice) |
|---|---|---|
| **Animal 1** | 1.152 | 1.179 |
| **Animal 2** | 1.170 | 1.185 |

### Figure 4e (i)

**Time to Death (SCID Mice after BCG Infection)**

| Days after Infection | BCG-WT (Tice) | BCG-*disA* -OE (Tice) | No treatment |
|---|---|---|---|
| 60 | | | |
| 61 | | | |
| 62 | | | |
| 63 | | | |
| 64 | | | |
| 65 | | | |
| 66 | | | |
| 67 | | | |
| 68 | | | |
| 69 | | | |
| 70 | | | |
| 71 | | | |
| 72 | | | |
| 73 | 1 | | |
| 74 | | | |
| 75 | | | |
| 76 | | | |
| 77 | | | |
| 78 | 1 | | |
| 79 | | | |
| 80 | | | |
| 81 | | | |
| 82 | | | |
| 82 | 1 | | |
| 83 | | | |
| 84 | | | |
| 85 | | | |
| 86 | 1 | | |
| 87 | | | |
| 88 | | | |
| 89 | | | |
| 90 | | | |
| 91 | | | |
| 92 | | | |
| 92 | | | |
| 93 | | | |

### Figure 4e (ii)

| | | |
|---|---|---|
| 94 | | |
| 95 | | |
| 96 | | |
| 97 | | |
| 98 | | |
| 98 | | |
| 99 | | |
| 100 | | |
| 101 | | |
| 102 | | |
| 103 | | |
| 104 | | |
| 105 | | |
| 106 | | |
| 107 | | |
| 108 | | |
| 109 | | 1 |
| 110 | | |
| 111 | | |
| 112 | | |
| 113 | | |
| 114 | | |
| 115 | | |
| 116 | | |
| 117 | | |
| 118 | | |
| 119 | | |
| 120 | | |
| 121 | | 1 |
| 122 | | |
| 123 | | |
| 124 | | |
| 125 | | |
| 126 | | |
| 127 | | |
| 128 | | 1 |
| 128 | 1 | |
| 128 | | |
| 128 | | |
| 129 | | |
| 130 | | |

### Figure 4e (iii)

| | | |
|---|---|---|
| 131 | | 1 |
| 132 | 1 | |
| 133 | | 1 |
| 133 | 1 | |
| 133 | | |
| 134 | | |
| 134 | 1 | |
| 135 | | |
| 136 | | 1 |
| 136 | 1 | |
| 136 | | |
| 136 | | |
| 137 | | |
| 138 | 1 | |
| 139 | | |
| 140 | | |
| 141 | | |
| 142 | | |
| 143 | | |
| 144 | | |
| 145 | | |
| 146 | | |
| 147 | | |
| 148 | | |
| 149 | | |
| 150 | | |
| 151 | | |
| 152 | | |
| 153 | | |
| 154 | | |
| 155 | | 1 |
| 156 | | |
| 157 | | |
| 158 | | |
| 159 | | |
| 159 | | |
| 160 | | |
| 161 | | |
| 162 | | |
| 163 | | |
| 164 | | |

### Figure 4e (iv)

| | | | |
|---|---|---|---|
| 165 | | | |
| 166 | | | |
| 167 | | 1 | |
| 168 | | | 0 |
| 169 | | | 0 |
| 170 | | | 0 |
| 171 | | 1 | 0 |
| 172 | | | 0 |
| 173 | | | 0 |
| 174 | | | 0 |
| 175 | | | 0 |
| 176 | | | 0 |
| 177 | | 1 | 0 |

### Figure 5a

**Raw Lucia ISG Reporter Assay (IRF Induction; RLU)**

| | R1 | R2 | R3 | R4 |
|---|---|---|---|---|
| **No treatment** | 13110 | 11060 | 11560 | 12020 |
| **BCG-WT (Tice)** | 141140 | 130610 | 169150 | 174280 |
| **BCG-disA-OE (Tice)** | 298050 | 293190 | 288970 | 333100 |
| **LPS (E.coli)** | 352020 | 349970 | 349410 | 330760 |
| **c-di-AMP (50 $\mu$g/ml)** | 257170 | 257280 | 267580 | 273880 |

### Figure 5b

| | IFN-$\beta$ (BMDM) (ELISA) Concentration in pg/ml | | | | | |
|---|---|---|---|---|---|---|
| | **WT-1** | **WT-2** | **WT-3** | **STING KO-1** | **STING KO-2** | **STING KO-3** |
| **No treatment** | 10.0 | 8.5 | 9.0 | 5.0 | 4.8 | 6.1 |
| **BCG-WT (Tice)** | 19.9 | 22 | 18.9 | 12.3 | 15.6 | 16.8 |
| **BCG-*disA* -OE (Tice)** | 45.0 | 48.0 | 42.0 | 13.2 | 14.9 | 11.2 |
| **c-di-AMP (50 $\mu$g/ml)** | 60.7 | 61.9 | 59.9 | 7.3 | 6.8 | 7.9 |

*WT: BMDM C57BL/6 (F)*

*STING KO: BMDM "STING-/- (C57BL/6J-Tmem173gt/J)*

### Figure 5c

**ELISA (Bone Marrow-Derived Macrophages)**
**IFN-$\beta$ (ELISA) (pg/ml)**

| | NT-1 | NT-2 | NT-3 | WT-1 | WT-2 | WT-3 | OE-1 | OE-2 | OE-3 | LPS-1 | LPS-2 | LPS-3 |
|---|---|---|---|---|---|---|---|---|---|---|---|---|
| **J774.1** | 4.79 | 5.68 | 5.56 | 13.99 | 10.29 | 10.76 | 19.75 | 20.75 | 25.83 | 31.70 | 15.81 | 16.79 |
| **HMDM** | 3.57 | 6.00 | 5.89 | 13.66 | 15.18 | 11.86 | 38.50 | 41.37 | 35.89 | 57.65 | 62.96 | 56.58 |

*NT: No treatment*

*WT: BCG-WT (Tice)*

*OE: BCG-disA-OE (Tice)*

*LPS: LPS (E.coli)*

### Figure 6a

**Macrophage reprogramming (BMDM infection with BCG) (Flow cytomeric analyses)**
**MOI (20:1)**

| | NT-1 | NT-2 | NT-3 | NT-4 | WT (P)-1 | WT (P)-2 | WT (P)-3 | WT (P)-4 | OE (P)-1 | OE (P)-2 | OE (P)-3 | OE (P)-4 |
|---|---|---|---|---|---|---|---|---|---|---|---|---|
| **%TNFα+ MHCII+ of CD11b+ F4/80+** | 0.58 | 0.75 | 0.60 | 0.76 | 16.0 | 16.1 | 22.4 | 23.3 | 25.6 | 23.5 | 21.6 | 22.3 |

*NT: No treatment*
*WT (P): BCG-WT (Pasteur)*
*OE (P): BCG-disA-OE*

### Figure 6b

**Macrophage reprogramming (BMDM infection with BCG) (Flow cytomeric analyses)**
**MOI (20:1)**

| | NT-1 | NT-2 | NT-3 | NT-4 | WT (P)-1 | WT (P)-2 | WT (P)-3 | WT (P)-4 | OE (P)-1 | OE (P)-2 | OE (P)-3 | OE (P)-4 |
|---|---|---|---|---|---|---|---|---|---|---|---|---|
| **%CD206+ CD124+ of CD11b+ F4/80+** | 79.7 | 81.8 | 80.5 | 79.5 | 43.6 | 41.1 | 37.5 | 36.4 | 20.0 | 19.3 | 17.8 | 17.2 |

*NT: No treatment*
*WT (P): BCG-WT (Pasteur)*

#### Figure 6c

**Macrophage reprogramming (BMDM infection with BCG) (Flow cytomeric analyses) MOI (20:1)**

| | NT-1 | NT-2 | NT-3 | NT-4 | WT (P)-1 | WT (P)-2 | WT (P)-3 | WT (P)-4 | OE (P)-1 | OE (P)-2 | OE (P)-3 | OE (P)-4 |
|---|---|---|---|---|---|---|---|---|---|---|---|---|
| **% IL-10 of CD206 +CD124+** | 3.23 | 3.36 | 3.21 | 3.34 | 22.3 | 17.7 | 31.7 | 29.8 | 7.31 | 4.32 | 9.46 | 6.02 |

*NT: No treatment*
*WT (P): BCG-WT (Pasteur)*

### Figure 6d

**MDSC population in mouse BMDM after BCG infection (Flow cytomeric analyses)**
**MOI (20:1)**

| | NT-1 | NT-2 | NT-3 | NT-4 | WT(P) -1 | WT(P) -2 | WT(P) -3 | WT(P) -4 | OE(P) -1 | OE(P) -2 | OE(P) -3 | OE(P) -4 |
|---|---|---|---|---|---|---|---|---|---|---|---|---|
| **%M-MDSC of $CD45^+$** | 0 | 0 | 0 | 0 | 1.78 | 2.21 | 1.59 | 2.26 | 0.55 | 0.49 | 0 | 0 |

*NT: No treatment*
*WT (P): BCG-WT (Pasteur)*
*OE (P): BCG-disA-OE (Pasteur)*

### Figure 6e

**MDSC population in mouse BMDM after BCG infection (Flow cytomeric analyses)**
**MOI (20:1)**

| | NT-1 | NT-2 | NT-3 | NT-4 | WT(P)-1 | WT(P)-2 | WT(P)-3 | WT(P)-4 | OE(P)-1 | OE(P)-2 | OE(P)-3 | OE(P)-4 |
|---|---|---|---|---|---|---|---|---|---|---|---|---|
| % IL-10 of M-MDSC | 0 | 0 | 0 | 0 | 18.5 | 30.7 | 18.3 | 22.0 | 0 | 2.28 | 0 | 0 |

*NT: No treatment*
*WT (P): BCG-WT (Pasteur)*
*OE (P): BCG-disA-OE*

### Figure 6f

**In Vitro Phagocytic Activity**
**Bone Marrow-Derived Macrophages (BMDM)**
**Mean Fluorescence Intensity**

| NT-1 | NT-2 | NT-3 | WT-1 | WT-2 | WT-3 | OE-1 | OE-2 | OE-3 |
|---|---|---|---|---|---|---|---|---|
| 782 | 734 | 754 | 1363 | 1407 | 2129 | 6271 | 5497 | 2605 |

*NT: No treatment*

*WT: BCG-WT (Tice)*

*OE: BCG-disA-OE (Tice)*

### Figure 6h

**Autophagic Flux (LC3B Staining)**
**Bone Marrow-Derived Macrophages (BMDM)**

| NT-1 | NT-2 | NT-3 | WT-1 | WT-2 | WT-3 | OE-1 | OE-2 | OE-3 |
|---|---|---|---|---|---|---|---|---|
| 0.19 | 0.20 | 0.12 | 2.50 | 2.40 | 1.90 | 3.16 | 2.85 | 2.45 |

**Autophagic Targeting (LC3B-BCG colocalization)**
**Bone Marrow-Derived Macrophages (BMDM)**

| WT-1 | WT-2 | WT-3 | OE-1 | OE-2 | OE-3 |
|---|---|---|---|---|---|
| 27.8 | 25.0 | 30.6 | 42.5 | 40.0 | 39.2 |

*NT: No treatment*

*WT: BCG-WT (Tice)*

*OE: BCG-disA-OE (Tice)*

### Figure 6j

**Autophagic Targeting (p62-BCG colocalization)**
**Bone Marrow-Derived Macrophages (BMDM)**

| Group | Nuclei | BCG bacilli | Colocalized BCG bacilli with p62 | % Colocalization |
|---|---|---|---|---|
| WT-1 | 4 | 12 | 3 | 25.0 |
| WT-1 | 4 | 8 | 2 | 25.0 |
| WT-1 | 4 | 12 | 2 | 16.6 |
| OE-1 | 4 | 14 | 7 | 50.0 |
| OE-2 | 4 | 14 | 6 | 42.9 |
| OE-3 | 4 | 12 | 4 | 33.3 |

### Figure 7a

**TNF-$\alpha$ qPCR (Human Monocytes from Healthy Donors)**
**Fold-change in Expression**

| | WT | OE |
|---|---|---|
| **D1** | 7.1 | 17.5 |
| **D2** | 6.8 | 17.7 |
| **D3** | 5.2 | 16.4 |
| **D4** | 10.0 | 15.4 |
| **D5** | 9.8 | 13.7 |
| **D6** | 10.8 | 14.9 |

*WT: BCG-WT (Tice)*

*OE: BCG-disA-OE (Tice)*

**IL-6 qPCR (Human Monocytes from Healthy Donors)**
**Fold-change in Expression**

| | WT | OE |
|---|---|---|
| **D1** | 1237 | 4169 |
| **D2** | 793 | 2806 |
| **D3** | 730 | 3168 |
| **D4** | 765 | 1562 |
| **D5** | 633 | 1457 |
| **D6** | 1037 | 2083 |

*WT: BCG-WT (Tice)*

*OE: BCG-disA-OE (Tice)*

### Figure 7c

**Chromatin Immunoprecipitation-Polymerase Chain Reaction (ChIP-PCR)**
**(Human Monocytes)**
**Chromatin Activation Mark (H3K4Me3) after BCG Training**
**Human Monocytes**
**IL-6 Promoter (% Input)**

| | NT-1 | NT-2 | NT-3 | NT-1 +PAM3CK4 | NT-2 +PAM3CK4 | NT-3 +PAM3CK4 |
|---|---|---|---|---|---|---|
| Donor | 0.024 | 0.054 | 0.14 | 0.63 | 0.12 | 0.22 |

| | WT-1 +RPMI | WT-2 +RPMI |
|---|---|---|
| Donor | 0.69 | 1.12 |

| WT-1 + PAM3CSK4 | WT-2 + PAM3CSK4 | WT-3 + PAM3CSK4 |
|---|---|---|
| 6.56 | 6.87 | 7.26 |

| | OE-1 +RPMI | OE-2 +RPMI | OE-3 +RPMI | OE-1 +PAM3CSK4 | OE-2 +PAM3CSK4 | OE-3 +PAM3CSK4 |
|---|---|---|---|---|---|---|
| Donor | 2.29 | 1.78 | 2.32 | 19.90 | 19.60 | 17.10 |

### Figure 7d

**In Vitro BCG Training**
**IL-6 ELISA (pg/ml) (Human Monocytes after BCG Training and Stimulation)**

| NT-1 | NT-2 | NT-3 | Rest-1 | Rest-2 | Rest-3 | Rest-1 + PAM3CSK 4 (Stim.) | Rest-2 + PAM3CSK 4 (Stim.) | Rest-3 + PAM3CS K4 (Stim.) |
|---|---|---|---|---|---|---|---|---|
| 15.6 | 12.7 | 19 | 13.7 | 14.2 | 14.3 | 408 | 372 | 396 |

| WT-1 + Rest | WT-2 +Rest | WT-3 +Rest | WT-1 +Rest +PAM3CS K4 (Stim.) | WT-2 +Rest +PAM3CS K4 (Stim.) | WT-3 +Rest +PAM3CS K4 (Stim.) |
|---|---|---|---|---|---|
| 268 | 226 | 231 | 431 | 445 | 449 |

| OE-1 +Rest | OE-2 +Rest | OE-3 +Rest | OE-1 +Rest +PAM3CS K4 (Stim.) | OE-2 +Rest +PAM3CS K4 (Stim.) | OE-3 +Rest +PAM3CS K4 (Stim.) |
|---|---|---|---|---|---|
| 313 | 388 | 376 | 555 | 565 | 541 |

*NT: No treatment*

*WT: BCG-WT (Tice)*

*OE: BCG-disA-OE (Tice)*

*Stim.: Stimulation*

### Figure 7e

**In Vitro BCG Training**
**TNF-$\alpha$ ELISA (pg/ml) (Human Monocytes after BCG Training and Stimulation)**

| NT-1 | NT-2 | NT-3 | Rest-1 | Rest-2 | Rest-3 | Rest-1+ PAM3CSK4 (Stim.) | Rest-2+ PAM3CSK4 (Stim.) | Rest-3+ PAM3CSK4 (Stim.) |
|---|---|---|---|---|---|---|---|---|
| 49.9 | 55.5 | 76.4 | 9.0 | 39.5 | 42.4 | 1251 | 1238 | 1322 |

| WT-1 +Rest | WT-2 +Rest | WT-3 +Rest | WT-1 +Rest +PAM3CSK4 (Stim.) | WT-2 +Rest +PAM3CSK4 (Stim.) | WT-3 +Rest +PAM3CSK4 (Stim.) |
|---|---|---|---|---|---|
| 254 | 292 | 273 | 1736 | 1793 | 1760 |

| OE-1 +Rest | OE-2 +Rest | OE-3 +Rest | OE-1 +Rest +PAM3CSK4 (Stim.) | OE-2 +Rest +PAM3CSK4 (Stim.) | OE-3 +Rest +PAM3CSK4 (Stim.) |
|---|---|---|---|---|---|
| 371 | 378 | 379 | 2000 | 2266 | 2381 |

*NT: No treatment*

*WT: BCG-WT (Tice)*

*OE: BCG-disA-OE (Tice)*

*Stim.: Stimulation*

#### Figure 8a

**Relative Abundance of Intracellular Metabolite (HMDM)**

| | WT-1 | WT-2 | OE-1 | OE-2 |
|---|---|---|---|---|
| **Glucose** | 68002 | 61340 | 103234 | 102895 |
| **Lactate** | 45899539 | 30163300 | 81380397 | 87077835 |
| **Tryptophan** | 1159366 | 1636769 | 1945251 | 1898404 |
| **Kynurenine** | 649069 | 745098 | 41300 | 40942 |

*WT: BCG-WT (Tice)*

*OE: BCG-disA-OE (Tice)*

### Figure 8b

**Relative Abundance of Intracellular Metabolite (BMDM)**

| | WT-1 | WT-2 | WT-3 | WT-4 | OE-1 | OE-2 | OE-3 | OE-4 |
|---|---|---|---|---|---|---|---|---|
| **Glucose** | 144972 | 94300 | 35240 | 112712 | 223466 | 223982 | 155474 | 154959 |
| **Lactate** | 309000000 | 367000000 | 420000000 | 408000000 | 546000000 | 539000000 | 537000000 | 535000000 |
| **Tryptophan** | 5095958 | 6000815 | 3667625 | 6288585 | 23414256 | 11263863 | 10891419 | 7355621 |

| | HK-WT-1 | HK-WT-2 | HK-WT-3 | HK-WT-4 | HK-OE-1 | HK-OE-2 | HK-OE-3 | HK-OE-4 |
|---|---|---|---|---|---|---|---|---|
| **Glucose** | 21397 | 83754 | 51961 | 97683 | 58805 | 128042 | 83549 | 91445 |
| **Lactate** | 366000000 | 352000000 | 295000000 | 210000000 | 265000000 | 359000000 | 314000000 | 319000000 |
| **Tryptophan** | 4930796 | 5327760 | 6979848 | 10649448 | 4408661 | 3109284 | 3977229 | 6137930 |

*WT: BCG-WT (Tice)*

*OE: BCG-disA-OE (Tice)*

*HK-WT: Heat-Killed  BCG-WT (Tice)*

*HK-OE: Heat-Killed  BCG-disA-OE (Tice)*

### Figure S1a

***disA*-qPCR in log-phase BCG cultures of BCG-WT (Tice and BCG-*disA*-**

| | WT-1 | WT-2 | WT-3 | OE-1 | OE-2 | OE-3 |
|---|---|---|---|---|---|---|
| **Fold expression** | 1.01 | 0.94 | 1.04 | 313.9 | 317.0 | 293.4 |

*WT: BCG-WT (Tice)*
*OE: BCG-disA-OE (Tice)*

#### Figure S1b

**RAW Lucia ISG Reporter Assay**
**(IRF3 Induction Assay)**

| | NT-1 | NT-2 | NT-3 | NT-4 | WT-1 | WT-2 | WT-3 | WT-4 |
|---|---|---|---|---|---|---|---|---|
| ISRE (R.L.U.) | 13114 | 11064 | 11560 | 12017 | 171552 | 159639 | 167460 | 170955 |

| | OE-1 | OE-2 | OE-3 | OE-4 | LPS-1 | LPS-2 | LPS-3 | LPS-4 |
|---|---|---|---|---|---|---|---|---|
| ISRE (R.L.U.) | 267912 | 263221 | 249400 | 277987 | 352021 | 349971 | 349409 | 330755 |

| | c-di-AMP-1 | c-di-AMP-2 | c-di-AMP-3 | c-di-AMP-4 |
|---|---|---|---|---|
| ISRE (R.L.U.) | 257169 | 257280 | 267583 | 273879 |

*NT: No treatment*
*WT: BCG-WT (Pasteur)*
*OE: BCG-disA-OE (Pasteur)*
*LPS: LPS (E.coli)*
*c-di-AMP: Cyclic di-AMP*

### Figure S3b

**% CD3+ of CD45 cells (MB49 Tumors)**
**Syngeneic MB49 heterotopic tumor model)**
**Flow Cytometry Analyses**

| Animal | C57BL/6 WT + Saline | STING$^{-/-}$ (C57BL/6J-Tmem173gt/J) + Saline | C57BL/6 WT + BCG-WT (Tice) | STING$^{-/-}$ (C57BL/6J-Tmem173gt/J) + BCG-WT (Tice) | C57BL/6 WT + BCG-*disA* -OE (Tice) | STING$^{-/-}$ (C57BL/6J-Tmem173gt/J) + BCG-*disA* -OE (Tice) |
|---|---|---|---|---|---|---|
| **A1** | 13.7 | 6.9 | 42.2 | 16.0 | 42.7 | 64.1 |
| **A2** | 11.3 | 7.1 | 21.2 | 16.5 | 36.4 | 16.2 |
| **A3** | 19.5 | 6.3 | 25.0 | 13.5 | 73.6 | 6.36 |
| **A4** | 14.2 | 8.9 | 28.9 | 19.8 | 34.4 | 14.3 |
| **A5** | 13.6 | 6.8 | 30.6 | 15.8 | 70.5 | 36.0 |
| **A6** | 14.1 | 8.1 | 35.0 | 21.1 | 23.9 | 16.8 |

*A: Animal*

#### Figure S3c

**% IFN-$\gamma$+ of CD8+ T cells (MB49 Tumors)**
**Syngeneic MB49 heterotopic tumor model)**
**Flow Cytometry Analyses**

| Animal | C57BL/6 WT + Saline | STING$^{-/-}$ (C57BL/6J-Tmem173gt/J) + Saline | C57BL/6 WT + BCG-WT (Tice) | STING$^{-/-}$ (C57BL/6J-Tmem173gt/J) + BCG-WT (Tice) | C57BL/6 WT + BCG-*disA* -OE (Tice) | STING$^{-/-}$ (C57BL/6J-Tmem173gt/J) + BCG-*disA* -OE (Tice) |
|---|---|---|---|---|---|---|
| **A1** | 13.4 | 12.3 | 23.9 | 48.2 | 38.1 | 27.5 |
| **A2** | 13.3 | 21.3 | 19.7 | 33.0 | 39.8 | 20.3 |
| **A3** | 13.6 | 17.8 | 27.4 | 21.6 | 38.7 | 15.2 |
| **A4** | 11.9 | 13.7 | 35.0 | 37.7 | 32.3 | 13.8 |
| **A5** | 16.5 | 12.7 | 26.2 | 30.8 | 31.1 | 15.8 |
| **A6** | 17.6 | 22.9 | 31.9 | 26.8 | 40.0 | 27.3 |

*A: Animal*

#### Figure S3d

**% CD69+CD38+ of CD8 T cells (MB49 Tumors)**
**Syngeneic MB49 heterotopic tumor model)**
**Flow Cytometry Analyses**

| Animal | C57BL/6 WT + Saline | STING$^{-/-}$ (C57BL/6J-Tmem173gt/J) + Saline | C57BL/6 WT + BCG-WT (Tice) | STING$^{-/-}$ (C57BL/6J-Tmem173gt/J) + BCG-WT (Tice) | C57BL/6 WT + BCG-*disA* -OE (Tice) | STING$^{-/-}$ (C57BL/6J-Tmem173gt/J) + BCG-*disA* -OE (Tice) |
|---|---|---|---|---|---|---|
| **A1** | 10.1 | 10.5 | 42.5 | 26.3 | 52.9 | 29.3 |
| **A2** | 11.4 | 7.3 | 10.2 | 45.3 | 26.8 | 27.4 |
| **A3** | 23.3 | 8.1 | 34.3 | 9.0 | 55.7 | 16.1 |
| **A4** | 9.7 | 10.9 | 27.3 | 36.3 | 47.4 | 27.0 |
| **A5** | 15.4 | 11.9 | 31.6 | 14.0 | 58.9 | 43.3 |
| **A6** | 9.2 | 3.7 | 49.5 | 33.1 | 55.6 | 46.6 |

*A: Animal*

#### Figure S3e

| | % TNF-$\alpha$+ of CD206+ CD124+ cells (MB49 Tumors) Syngeneic MB49 heterotopic tumor model) Flow Cytometry Analyses | | | | | |
|---|---|---|---|---|---|---|
| **Animal** | **C57BL/6 WT + Saline** | **STING$^{-/-}$ (C57BL/6J-Tmem173gt/J) + Saline** | **C57BL/6 WT + BCG-WT (Tice)** | **STING$^{-/-}$ (C57BL/6J-Tmem173gt/J) + BCG-WT (Tice)** | **C57BL/6 WT + BCG-*disA* -OE (Tice)** | **STING$^{-/-}$ (C57BL/6J-Tmem173gt/J) + BCG-*disA* -OE (Tice)** |
| **A1** | 13.5 | 6.6 | 27.9 | 19.4 | 21.0 | 20.4 |
| **A2** | 10.2 | 5.1 | 27.2 | 36.4 | 74.9 | 24.1 |
| **A3** | 13.2 | 6.8 | 39.2 | 14.6 | 57.1 | 10.4 |
| **A4** | 13.3 | 6.2 | 77.9 | 18.8 | 77.0 | 18.0 |
| **A5** | 15.1 | 5.6 | 100.0 | 18.6 | 59.4 | 18.9 |
| **A6** | 10.7 | 9.5 | 22.9 | 50.3 | 41.8 | 51.2 |

*A: Animal*

#### Figure S4a

**Lung Bacillary Burden BALB/c Mice**
**D1 Implantation**
**Raw CFU per lung and log10 CFU per lung**
**(half of total lung homogenate plated on each of 2 plates).**

| CFU counts per plate (each animal lung plated in duplicate) (D1 implantation) | | | | |
|---|---|---|---|---|
| **Strain** | **Plate 1** | **Plate 2** | **CFU (total)** | **Log10cfu** |
| **BCG-WT (Tice)** | | | | |
| **A1** | 53 | 57 | 110 | 2.045 |
| **A2** | 56 | 81 | 137 | 2.140 |
| **A3** | 66 | 99 | 165 | 2.220 |
| **BCG-*disA* -OE (Tice)** | | | | |
| A1 | 69 | 41 | 110 | 2.045 |
| A2 | 253 | 142 | 395 | 2.598 |
| A3 | 242 | 110 | 352 | 2.548 |

### Figure S4b

**Lung Bacillary Burden**
**(Day 28 Implantation in Balb/c Mice (Female))**

| | BCG-WT (Pasteur) | BCG-*disA* -OE (Pasteur) |
|---|---|---|
| **Animal1** | 6.155 | 5.685 |
| **Animal2** | 6.083 | 5.621 |
| **Animal3** | 5.939 | 5.610 |
| **Animal4** | 6.162 | 5.976 |
| **Animal5** | 6.246 | 5.573 |

### Figure S4c

**Lung Bacillary Burden**
**(Day 1 Implantation in SCID Mice (Female)**

| | BCG-WT (Pasteur) | BCG-*disA* -OE (Pasteur) |
|---|---|---|
| **Animal1** | 1.204 | 1.114 |
| **Animal2** | 1.146 | 0.954 |

### Figure S4d (i)

| Survival of SCID Mice infected with BCG<br>Time to Death Analyses | | | |
|---|---|---|---|
| Days | BCG-WT (Pasteur) | BCG-*disA* -OE (Pasteur) | No treatment |
| 60 | | | |
| 61 | | | |
| 62 | | | |
| 63 | | | |
| 64 | | | |
| 65 | | | |
| 66 | | | |
| 67 | | | |
| 68 | | | |
| 69 | | | |
| 70 | | | |
| 71 | | | |
| 72 | | | |
| 73 | | | |
| 74 | | | |
| 75 | | | |
| 76 | | | |
| 77 | | | |
| 78 | | | |
| 79 | | | |
| 80 | | | |
| 81 | | | |
| 82 | | | |
| 82 | | | |
| 83 | | | |
| 84 | 1 | | |
| 85 | | | |
| 86 | | | |
| 87 | | | |
| 88 | | | |
| 89 | | | |
| 90 | | | |
| 91 | | | |
| 92 | 1 | | |
| 92 | | | |
| 93 | 1 | | |

#### Figure S4d (ii)

| | | |
|---|---|---|
| 94 | | |
| 95 | | |
| 96 | | |
| 97 | | |
| 98 | | 1 |
| 98 | 1 | |
| 99 | 1 | |
| 100 | | |
| 101 | | |
| 102 | | |
| 103 | | |
| 104 | | |
| 105 | | |
| 106 | | |
| 107 | | |
| 108 | | |
| 109 | | |
| 110 | | |
| 111 | | |
| 112 | | |
| 113 | | |
| 114 | | |
| 115 | | |
| 116 | | 1 |
| 117 | | |
| 118 | | |
| 119 | | |
| 120 | | |
| 121 | | |
| 122 | | |
| 123 | | |
| 124 | | |
| 125 | | |
| 126 | | |
| 127 | | |
| 128 | | |
| 128 | | |
| 128 | | 1 |
| 128 | 1 | |
| 129 | | |
| 130 | 1 | |

### Figure S4d (iii)

| | | |
|---|---|---|
| 131 | | |
| 131 | 1 | |
| 132 | | |
| 133 | | |
| 133 | | |
| 133 | | 1 |
| 134 | | 1 |
| 134 | | |
| 135 | 1 | |
| 136 | | |
| 136 | | |
| 136 | 1 | |
| 136 | | 1 |
| 137 | | 1 |
| 138 | | |
| 139 | | |
| 140 | | |
| 141 | | |
| 142 | | |
| 143 | | |
| 144 | | |
| 145 | | |
| 146 | | |
| 147 | | |
| 148 | | |
| 149 | | |
| 150 | | |
| 151 | | |
| 152 | | |
| 153 | | |
| 154 | | |
| 155 | | |
| 156 | | |
| 157 | | |
| 158 | | |
| 159 | | 1 |
| 159 | | |
| 160 | | 1 |
| 161 | | |
| 162 | | |
| 163 | | |

### Figure S4d (iv)

| | | | |
|---|---|---|---|
| 164 | | | |
| 165 | | | |
| 166 | | | |
| 167 | | | |
| 168 | | 1 | 0 |
| 169 | | | 0 |
| 170 | | | 0 |
| 171 | | | 0 |
| 172 | | | 0 |
| 173 | | | 0 |
| 174 | | | 0 |
| 175 | | | 0 |
| 176 | | | 0 |
| 177 | | | 0 |

### Figure S5

**IFN-$\beta$ q-PCR in resting and IFN-$\gamma$ primed bone-marrow-derived macrophages**

| | NT-1 | NT-2 | NT-3 |
|---|---|---|---|
| IFN-$\gamma$ (-) | 0.874 | 1.041 | 1.099 |
| IFN-$\gamma$ (+) | 0.990 | 1.020 | 1.040 |

| | WT (P)-1 | WT (P)-2 | WT (P)-3 | OE (P)-1 | OE (P)-2 | OE (P)-3 |
|---|---|---|---|---|---|---|
| IFN-$\gamma$ (-) | 1.855 | 1.857 | 1.635 | 5.861 | 5.413 | 4.671 |
| IFN-$\gamma$ (+) | 1.922 | 0.872 | 1.371 | 12.804 | 16.244 | 10.175 |

| | WT (T)-1 | WT (T)-2 | WT (T)-3 | OE (T)-1 | OE (T)-2 | OE (T)-3 |
|---|---|---|---|---|---|---|
| IFN-$\gamma$ (-) | 3.610 | 3.647 | 4.084 | 6.652 | 6.565 | 6.217 |
| IFN-$\gamma$ (+) | 7.179 | 5.510 | 9.697 | 25.711 | 23.933 | 20.550 |

| | LPS-1 | LPS-2 | LPS-3 |
|---|---|---|---|
| IFN-$\gamma$ (-) | 44.494 | 41.608 | 42.005 |
| IFN-$\gamma$ (+) | 585.049 | 547.881 | 570.034 |

*NT: No treatment*
*WT (P): BCG-WT (Pasteur)*
*OE (P): BCG-disA-OE (Pasteur)*
*WT (T): BCG-WT (Tice)*

### Figure S11

| Quantification lof LC3B-BCG colocalization in 5637 cells MOI (20:1) | | | | | | |
|---|---|---|---|---|---|---|
| | **WT (T)-1** | **WT (T)-2** | **WT (T)-3** | **OE (T)-1** | **OE (T)-2** | **OE (T)-3** |
| **% LC3B-BCG colocalization** | 10 | 5 | 12 | 25 | 28 | 35 |

*WT (T): BCG-WT (Tice)*
*OE (T): BCG-disA-OE (Tice)*

### Figure S12b

| GLUT1 expression on BMDMs after BCG infection (Flow cytomeric analyses) MOI (20:1) | | | | | | |
|---|---|---|---|---|---|---|
| | **NT-1** | **NT-2** | **WT (T)-1** | **WT (T)-2** | **OE (T)-1** | **OE (T)-2** |
| **Glut1+ macrophages** | 5.6 | 5.5 | 13.5 | 12.0 | 27.5 | 27.6 |

*NT: No treatment*
*WT (T): BCG-WT (Tice)*
*OE (T): BCG-disA-OE (Tice)*

### Figure S12c

| | GLUT1 expression on BMDMs after BCG infection (Flow cytomeric analyses) MOI (20:1) | | | | | |
|---|---|---|---|---|---|---|
| | **NT-1** | **NT-2** | **WT (T)-1** | **WT (T)-2** | **OE (T)-1** | **OE (T)-2** |
| **2-NBDG Uptake (MFI)** | 355 | 360 | 420 | 415 | 491 | 510 |

*NT: No treatment*
*WT (T): BCG-WT (Tice)*
*OE (T): BCG-disA-OE (Tice)*
